# Discovery and biosynthesis of biffamycin A – a novel glycotetrapeptide antibiotic

**DOI:** 10.1101/2025.09.22.677794

**Authors:** Michael W. Brigham, Edward S. Hems, Daniel C. L. Van, Justin E. Clarke, Sergey Nepogodiev, Christian Bassi, Michael E. Webb, Glyn R. Hemsworth, Barrie Wilkinson, Ryan F. Seipke

## Abstract

The clinical deployment of antibiotics is undermined by antimicrobial resistance. Without new agents to treat antibiotic resistant bacterial infections, mortality rates are predicted to reach 10 million people per year by 2050. Most antibiotics are derived from natural products (NPs) produced by bacteria; however, this resource was abandoned by industry because of high rediscovery rates. We are amid a natural product renaissance fuelled by inexpensive access to genome sequencing and sophisticated bioinformatic tools, which have highlighted that most of the biosynthetic pathways for NPs are not expressed in the laboratory. Here, we engineered the expression of a silent biosynthetic gene cluster harboured by an environmental isolate of *Streptomyces albidoflavus*. By using a bioinformatics-guided approach, we isolated and structurally characterised a novel glycopeptide antibiotic (GPA) named biffamycin A, which is the smallest GPA known and harbours unprecedented 5-chloro-4-methoxy tryptophan and 3-hydroxy(α-d-mannoysl)-d-lysine moieties. Biffamycin A possesses antimycobacterial and antistaphylococcal bioactivity, including methicillin-vancomycin-resistant *Staphylococcus aureus*.

## Introduction

A key challenge in antimicrobial drug development is the discovery of agents that circumvent clinically prevalent resistance mechanisms, which are particularly problematic for the ESKAPE group of pathogens (*Enterococcus faecium, Staphylococcus aureus, Klebsiella pneumoniae, Acinetobacter baumannii, Pseudomonas aeruginosa* and *Enterobacter* species). Most antibiotics in use are derived from microbial natural products, particularly those produced by *Streptomyces* bacteria and other filamentous Actinobacteria.^[1]^ *Streptomyces* species harbour many biosynthetic pathways, but only a handful of them are typically productive in a laboratory setting.^[2]^ Accessing the chemical diversity encoded by such biosynthetic pathways is widely believed to be the best route to a second golden era of antibiotic discovery.

Nonribosomal peptides (NRPs) are a structurally complex and diverse family of natural products that often exhibit therapeutically relevant activities. NRPs are produced by multifunctional enzymes called nonribosomal peptide synthetases (NRPSs), which are large assembly line-like machines organised into modules whose biochemical function is to incorporate a single monomeric building block into the growing polypeptide. NRPS biosynthetic modules can be grouped into two categories, loading modules and elongation modules.^[3]^ A loading module generally consists of two active domains, an adenylation (A) domain that activates an amino acid substrate and loads it onto the second domain, and a peptidyl carrier protein (PCP) domain, which possess a 4′-phosphopantetheinyl prosthetic group to which the growing peptide chain remains covalently linked. Elongation modules also harbour A and PCP domains, but additionally contain a condensation (C) domain, which precedes the A domain (C-A-PCP), and catalyzes peptide bond formation between two PCP-bound peptide units. Both loading and elongation modules can harbour additional tailoring domains such as epimerase (E) domains amongst others that modify peptide intermediates. The terminal elongation module usually possesses a C-terminal thioesterase (TE) domain, which transfers the polypeptide intermediate from the final PCP domain onto a conserved Ser residue, after which either a hydrolytic or macrocyclisation reaction occurs to produce the mature peptide or depsipeptide.

Genome mining enabled us and others to discover a novel standalone peptide cyclase (SurE) from the surugamide biosynthetic system.^[4–6]^ SurE is the archetypical member of what has been dubbed the penicillin-binding protein (PBP)-TE family, orthologues of which are widespread in *Streptomyces* species with about 15% of sequenced strains harbouring at least one NRPS system with a SurE offloading strategy.^[4]^ Our continued characterisation of the surugamide-producing strain *S. albidoflavus* S4^[4]^ led us to identify a silent/cryptic NRPS system that, instead of harbouring a standalone SurE cyclase, contains a *cis-*encoded (*i*.*e*., embedded) SurE domain at the C-terminus of the terminal biosynthetic module, a unique observation within NRPS megaenzymes. This intriguing observation was the impetus for a targeted campaign to access the product of this pathway, which is revealed here to be a novel glycotetrapeptide antibiotic with unprecedented 5-chloro-4-methoxy tryptophan and 3-hydroxy(α**-**D-mannoysl)-d-lysine moieties.

## Results and Discussion

### *Identification and analysis of the biffamycin* (bif) *biosynthetic gene cluster (BGC)*

The *bif* BGC harboured by *Streptomyces albidoflavus* S4^[7]^ is composed of 17 genes (*bifABCDEFGHIJKLMNOP*) predicted to encode the production of a modified glycotetrapeptide (Figure 1, Table S1). Bioinformatic analyses were used to predict the putative product of the *bif* BGC, which served as a framework for its experimental identification. This framework considered the following: the two dimodular nonribosomal peptide synthetases, BifC (domain organisation: A1-PCP1-C1-A2-PCP2) and BifB (domain organisation: C2-A3-PCP3-E3-C3-A4-PCP4-SurE) assemble a peptide scaffold, which, based on analysis of the predicted adenylation domain active site residues using PARAS (Figure S1)^[8]^, is predicted to be cyclo[L-Val-L-Trp-D-Lys-Gly]. The terminal biosynthetic intermediate is likely cyclised by a domain that shows high predicted structural similarity to a standalone peptide cyclase named SurE (Figure S2-S3).^[4-6]^ The presence of an E-domain within the third module (Figure S4) indicates an amino acid in the D-configuration is likely present at the third position of the final molecule. The resulting cyclotetrapeptide scaffold is predicted to be mono-hexosylated by a putative hexosyl transferase (BifP), the substrate for which is provided by the polyprenyl-P-hexose synthase (BifO). Analysis of other proteins encoded within the BGC indicated that two amino acids, one basic polar, and the other aromatic, are likely modified either on-board or prior to activation by their cognate A-domains. BifEF share 49% and 60% sequence identity with OhmKJ, respectively, which were recently proposed to introduce a methoxy group to the Trp moiety of ohmyungsamycin^[11]^, while BifKI constitute a predicted Trp halogenase/flavin reductase pair. BifN is a putative α-ketoglutarate (α-KG)-dependent amino acid hydroxylase with 41% shared amino acid identity to VioC, a structurally characterised standalone L-Arg 3*S*-hydroxylase.^[8–10]^ Thus, BifEFIK likely generates Cl-MeO-Trp, and BifN may hydroxylate Lys, the presumed substrates for the second and third biosynthetic modules, respectively. Taken together, these analyses allowed us to predict that biffamycin would consist of a glycopeptide composed of hexosylated cyclo[L-Val-L-(Cl-MeO-Trp)-(3-OH-D-Lys)-Gly].

**Figure 1.**
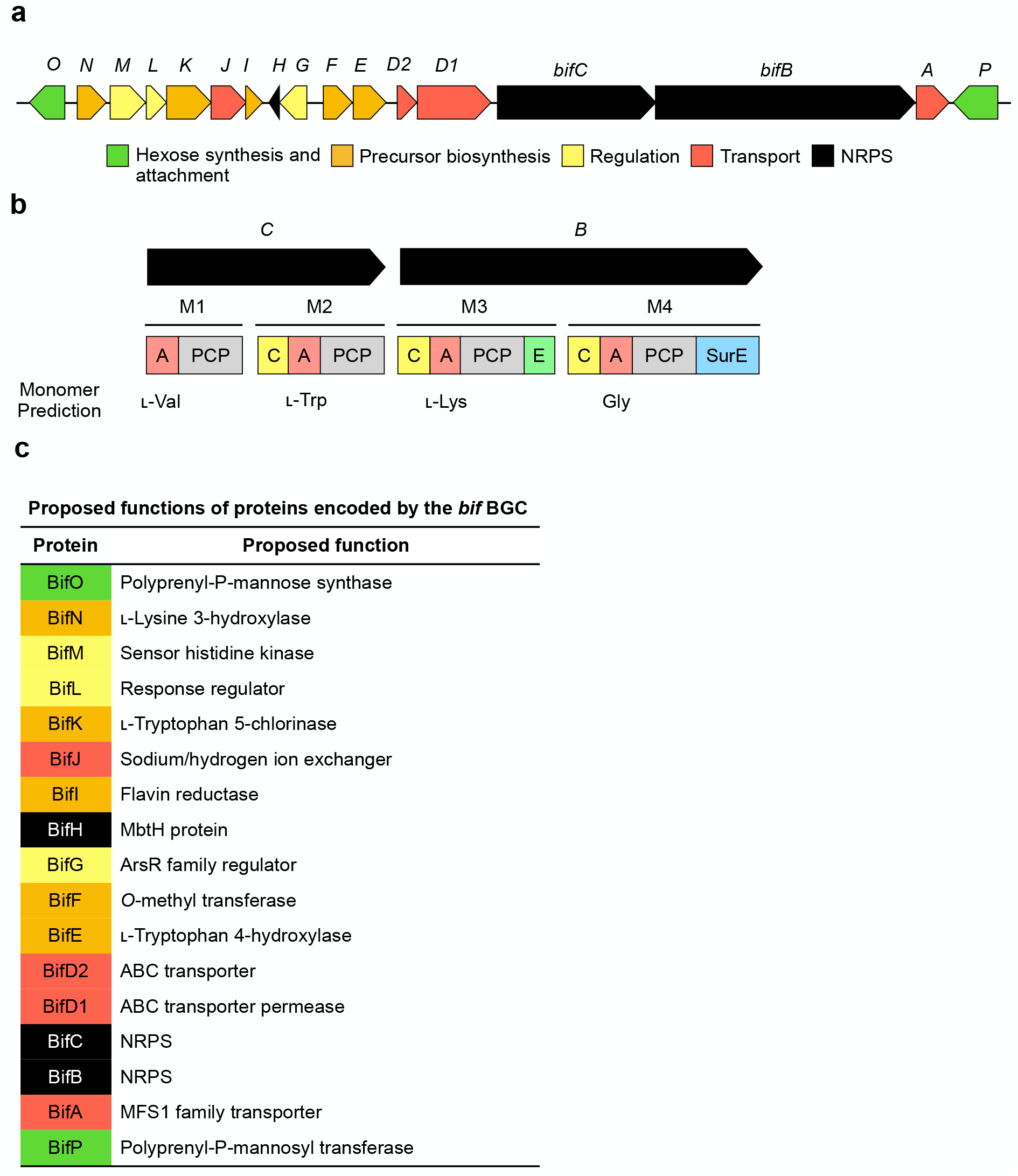
The biffamycin *(bif)* biosynthetic gene cluster (BGC). (a) Gene organisation of the *bif* BGC colour-coded by deduced function as indicated. (b) Biosynthetic module organisation of BifC and BifB. A, adenylation domain; T, thiolation domain; C, condensation domain; E, epimerase domain; SurE, macrocyclisation domain. Domain organisation was predicted using antiSMASH v6.1.1^[14]^ and A-domain specificities were predicted using PARAS.^[15]^ (c) Proposed function of proteins encoded by the *bif* BGC based upon bioinformatic analyses and experiments performed during this study. The *bif* BGC is available under Genbank accession PV425438.

### *Deregulation of the* bif *BGC and chemical analysis of biffamycin A*

To access the product(s) of the *bif* BGC, we cultivated a variant of *S. albidoflavus* S4 (*S. albidoflavus* S4Δ5, in which five endogenous BGCs are inactivated^[12]^), on a small suite of agar growth media (see methods). Methanolic extracts were prepared from these cultures, concentrated, and analysed by LC-HRMS. Minute quantities of two metabolites broadly consistent with the molecular weight associated with our bioinformatic prediction were detected in the extract prepared from minimal medium (MM)-grown culture; however, low production titre precluded their detailed analysis (Figure 2a). Thus, a promoter engineering strategy was used to de-regulate transcription of the *bif* BGC. The genes *bifO-*to-*bifD1* were replaced with a hygromycin resistance gene harbouring an *ermE*\* promoter to facilitate constitutive expression of *bifCBA* on the chromosome of *S. albidoflavus* S4 Δ5 (Δ*bifO-bifD1*). Next, an integrative plasmid harbouring the genes *bifPOKJIFENHD2D1* constitutively expressed from *ermE*\* and *rpsL(XC)* promoters was constructed and introduced into the Δ*bifO-bifD1* strain to create Δ*bifO-bifD1*/pBiff (Figure S5). The Δ*bifO-bifD1*/pBiff strain was cultivated in MM as above and crude methanolic extracts were prepared and analysed by LC-HRMS. De-regulation of the *bif* BGC led to reliable detection of two compounds with an isotopic distribution pattern consistent with a singly chlorinated species, and a mass consistent with the predicted hexosylated compound ((calc. *m/z* 713.2908, obs. *m/z* 713.2811, [M+H]^+^ C_31_H_45_ClN_6_O_11_) and (calc. *m/z* 727.3064, obs. *m/z* 727.2974, [M+H]^+^ C_32_H_47_ClN_6_O_11_)) (Figure 2ab). The latter compound was the more abundant product and therefore was named biffamycin A (**1**) and is the focus of this study; we named the former compound biffamycin B (**2**). It is common for NRPS systems to produce a small suite of congeners differing in their aliphatic amino acid content^[13]^, so we hypothesised the mass increase of *m/z* 14.0066 for **1** compared to our bioinformatic prediction indicated that the compound may harbour an amino acid with an additional methylene group compared to Val, presumably Ile or Leu. To simplify purification, we sought to determine if we could skew the production profile of **1** relative to **2** by supplementing growth media with specific amino acids. Δ*bifO-bifD1*/pBiff was therefore cultivated in MM supplemented with either Val, Ile or Leu. Interestingly, supplementation of MM with Ile or Leu led to a marked increase in **1** and no detectable **2**, and supplementation of MM with Val led to increased **2** and only a trace level of **1** (Figure S6). Taken together, these data establish that the *bif* BGC encodes production of at least two metabolites whose production can be modulated by the composition of culture media. As we could not disambiguate whether **1** possessed Ile or Leu by mass spectrometry alone, we chose to characterise **1** from cultures supplemented with Ile.

**Figure 2.**
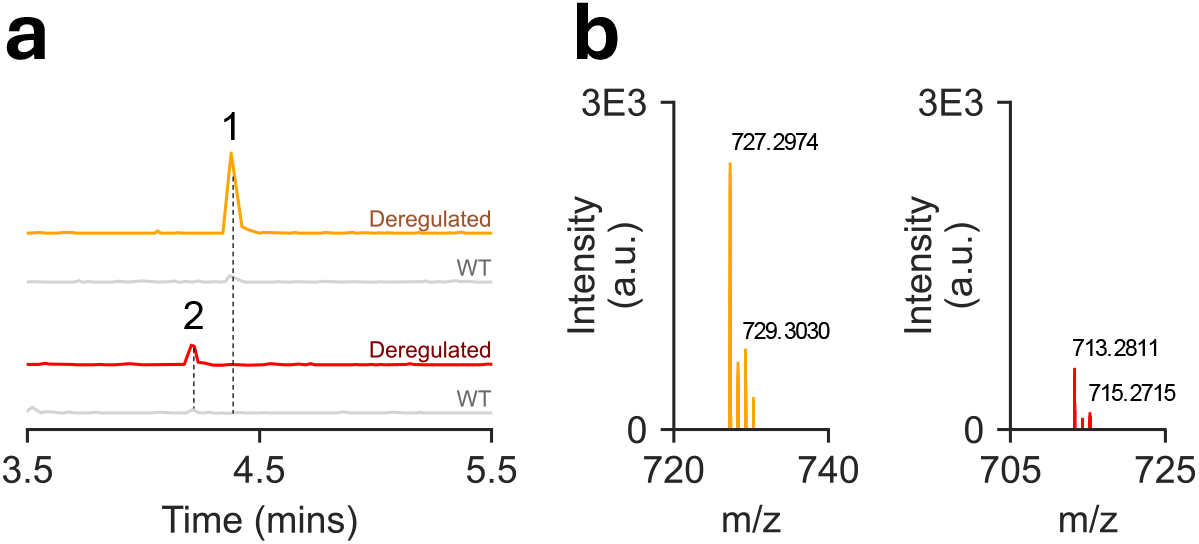
De-regulation of the *bif* BGC results in production of biffamycins. (a) Production of biffamycin A (**1**) and B (**2**) in minimal medium (MM) by either the WT or the deregulated *S. albidoflavus* S4Δ5 Δ*bifO-bifD1*/pBiff strain. Displayed are extracted ion chromatograms (EICs) for the monoisotopic masses corresponding to the [M+H]^+^ adduct for **1** (C_32_H_47_ClN_6_O_11_) ± 0.01 and **2** (C_31_H_45_ClN_6_O_11_) ± 0.01. (b) Mass spectra for **1** (left) and **2** (right); a.u., arbitrary units. Colours coordinated with companion Figure S6.

### Purification and structure elucidation of biffamycin A (1)

To purify **1**, the producer organism was first cultivated in 8 L of MM supplemented with 5 mM Ile production medium. Following centrifugation and filtration, the clarified supernatant was passed through a C_18_ flash chromatography cartridge to capture **1**, after which the retained components were eluted. The resulting fractions were analysed by LCMS and those containing target compounds were further purified by preparative HPLC to yield biffamycin A (**1**, 3.5 mg), *des-*hydroxy-biffamycin A aglycon (**3**, 180 µg), the precursor 5-Cl-4-MeO-L-Trp (**4**, 15.6 mg) and an acetylated derivative, *N*-acetyl-5-Cl-4-MeO-L-Trp (**5**, 4.4 mg). In addition, the cell pellet was extracted with methanol, and the resulting extract purified by preparative HPLC to yield the biffamycin A aglycon (**6**, 230 µg) and additional **3** (230 µg).

Unfortunately, **1** did not have sufficient solubility in common deuterated solvents for NMR analysis and, as such, the structure of **1** was elucidated using complementary approaches. First, we considered the LCHRMS^2^ data for **1, 3** and **6** (Figure 3a, Figures S7,S9-10 and Tables S2,S4-5). **1** gave reliable in-source fragmentation across several MS instruments with a loss of *m/z* 162, characteristic of a hexose moiety.

**Figure 3.**
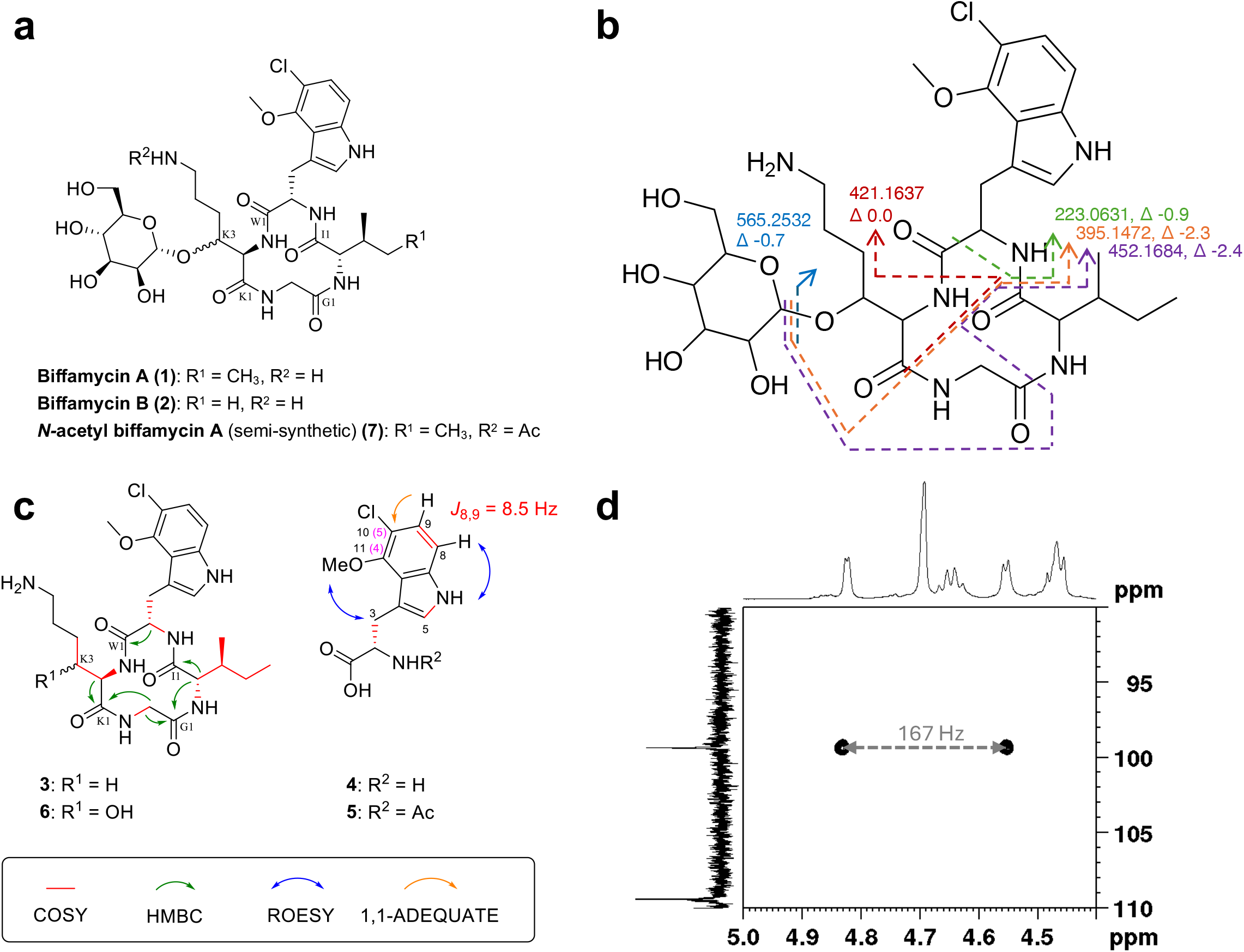
Chemical characterisation of **1**. (a) Structures of biffamycin A (**1**), biffamycin B (**2**) and semi-synthetic *N*-acetyl-biffamycin A (**7**). (b) LCHRMS^2^ fragmentation pattern for **1** showing *m/z* and error (ppm) of key fragments used to determine amino acid connectivity. (c) Key 2D NMR correlations used in the structural assignment of compounds **3**-**6**. The IUPAC numbering for the indole ring on **4** and **5** is shown in magenta. (d) Zoomed coupled HSQCed spectrum of **7** showing the anomeric signal as a doublet with ^1^*J*_CH_ = 167.4 Hz.

Comparison of the MS^2^ spectra of **1** with that of **6** showed that after the loss of a hexose, the fragmentation data were essentially identical. This gave strong evidence that **1** and **6** share the same tetrapeptide backbone, with **1** being glycosylated with a hexose. The predicted connectivity of the cyclotetrapeptide backbone of **1** from bioinformatics was supported by detailed MS^2^ analysis of both **1** and **6**, as well as full assignment of the 1D and 2D NMR spectra of **6** (Figure 3b, Figure S34-40 and Table S7). Here, COSY was utilised to assign the Gly, Ile and 3-OH-Lys side chains of **6**. The HMBC cross peaks between the α-hydrogens of Gly and the amide carbon of 3-OH-Lys, and the a-hydrogen of Ile and amide carbon of Gly established the (3-OH-D-Lys)-Gly-L-Ile sequence which, in combination with MS^2^, confirmed the amino acid connectivity as cyclo[L-Ile-(5-Cl-4-MeO-L-Trp)-(3-OH-D-Lys)-Gly]. Using the same MS^2^ analysis methods suggested a structure for **3** of cyclo[L-Ile-(5-Cl-4-MeO-L-Trp)-D-Lys-Gly] (Figure 3b, Figures S10-16 and Table S6). Comparison of the ^1^H and ^13^C NMR of **6** and **3** showed great similarities between the corresponding NMR spectra except for a clear downfield shift at the C3 resonance of the D-Lys residue for **6** (δ_H_ 3.75, δ_C_ 68.9) when compared with **3** (δ_H_ 1.63 & 1.50, δ_C_ 29.4), indicating **6** is hydroxylated at the C3 on the D-Lys side chain. The stereochemistry of this hydroxyl group could not be resolved, and in all cases the stereochemistry of D-Lys was inferred from the bioinformatic analysis.

Next, the regiochemistry of the Cl and MeO substituents on the presumed NRPS substrate **4** were assigned by NMR (Figure 3b, Figure S20-26 and Table S7). First the aromatic protons were considered, with a singlet at δ_H_ 7.21 assigned to H5 and two doublets at δ_H_ 7.15 and δ_H_ 7.07 with a coupling constant of 8.5 Hz suggesting a pair of protons with an *ortho* orientation on the indole ring. Next, analysis of the ROESY NMR showed an interaction between the indole N*H* proton at δ_H_ 11.21 and the aromatic doublet of H8 at δ_H_ 7.15, which indicated the *ortho* aromatic protons were on C8 and C9 (Figure S26). Additional ROESY cross coupling between the methoxy protons (H13) and H2 and H3’ of the tryptophan group indicated the methoxy group was on C11 (δ_C_ 149.3). Further analysis by 1,1-ADEQUATE NMR showed a cross coupling between H9 and C10 (δ_C_ 115.8 ppm) indicating the position of the chlorine atom (Figure S25). Taken together, **4** was assigned as 5-chloro-4-methoxy-L-tryotophan using IUPAC nomenclature. *N*-acetyl-5-chloro-4-methoxy-L-tryptophan (**5**) was characterised in an analogous manner (Figure 3b, Figures S27-33 and Table S7).

To determine the hexose moiety on biffamycin A (**1**), 50 µg of **1** were hydrolysed with 1M aqueous trifluoracetic acid. The liberated hexose was identified as mannose by co-injection/migration of standards by high performance anion exchange chromatography with pulsed amperometric detection (HPAEC-PAD) to be mannose (Figure S12). We are unaware of a microbial source of L-mannose^[16]^ and therefore presume mannose to be in the D-configuration.

Finally, to address the problem of solubility in NMR solvents, a sample of **1** was *N*-acetylated using acetic anhydride in pyridine to give **7** which was partially soluble in DMSO-d_6_ (Figure 3a, Figures S41-446,S56 and Table S7). LCHRMS^2^ analysis of **7** confirmed the compound was *N*-acetylated on the 3-hydroxy-D-lysine side chain (Figure S11 and Table S6). The solubility of **7** allowed us to analyse the mannose glycoside. HMBC was used to determine that **7** was glycosylated at the hydroxyl group on C3 of the D-Lys side chain, with a cross peak between the mannose anomeric proton (δ_H_ 4.70) and K3 (δ_C_ 75.2) (Figure S45). To determine the anomeric configuration of mannose, we ran a coupled HSQCedited spectrum to measure the one-bond carbon-proton coupling constants. For the anomeric carbon ^1^*J*_CH_ = 167.4 Hz, which would suggest an α-D-mannopyranose residue on **7** (Figure 3).^[17]^ For comparison, we also ran the coupled HSQCedited spectrum for methyl α-D-mannopyranoside in DMSO-d_6_ for which the anomeric carbon gave ^1^*J*_CH_ = 167.5 Hz (Figure S47). Thus, the final proposed structure of **1** is assigned as: cyclo[L-Ile-(5-Cl-4-MeO-L-Trp)-(3-O-α-D-mannosyl-D-Lys)-Gly].

### Reconstitution of 5-Cl-4-MeO-L-Trp and 3-OH-L-Lys biosynthesis

The chemical structure of **1** revealed the presence of two modified amino acid residues (5-Cl-4-MeO-L-Trp, and 3-OH-D-Lys) in the cyclotetrapeptide scaffold. We purified the former from culture supernatant and elucidated its structure above. To the best of our knowledge, chloro-methoxy-tryptophans have not previously been reported from a natural or synthetic source, we therefore sought to reconstitute the biosynthesis of 5-Cl-4-MeO-L-Trp (**4**) *in vitro*. However, despite considerable effort, the BifK halogenase could not be produced as a soluble protein from *E. coli*. Thus, we established the biosynthetic origin of **4** through a combination of *in vitro* reconstitution and *in vivo* heterologous expression studies. (His)_6_-BifE was heterologously produced and purified from *E. coli* (Figures S48-S49), and the hydroxylation activity of BifE and heat-inactivated BifE were assessed *in vitro* using L-Trp and commercially available 5-Cl-L-Trp, which established BifE utilises Trp and not 5-Cl-L-Trp as a substrate (Figure 4a). Next, (His)_6_-BifF was overproduced and purified from *E. coli* (Figures S48 and S50), and a one-pot reaction was developed to successfully reconstitute the methoxylation of L-Trp *in vitro* (Figure 4b). This enzymatic route for L-Trp methoxylation is intriguing; while tryptophan hydroxylases are known to catalyse oxygenation of the indole ring in animals, an equivalent enzyme in bacteria remained enigmatic until recently. During **1** biosynthesis, C4 oxygenation of L-Trp is catalysed by a rare heme-dependent Trp hydroxylase (BifE) that belongs to a new sub-family of the Trp-2,3-dioxygenases reported while our study was in progress.^[18]^

**Figure 4.**
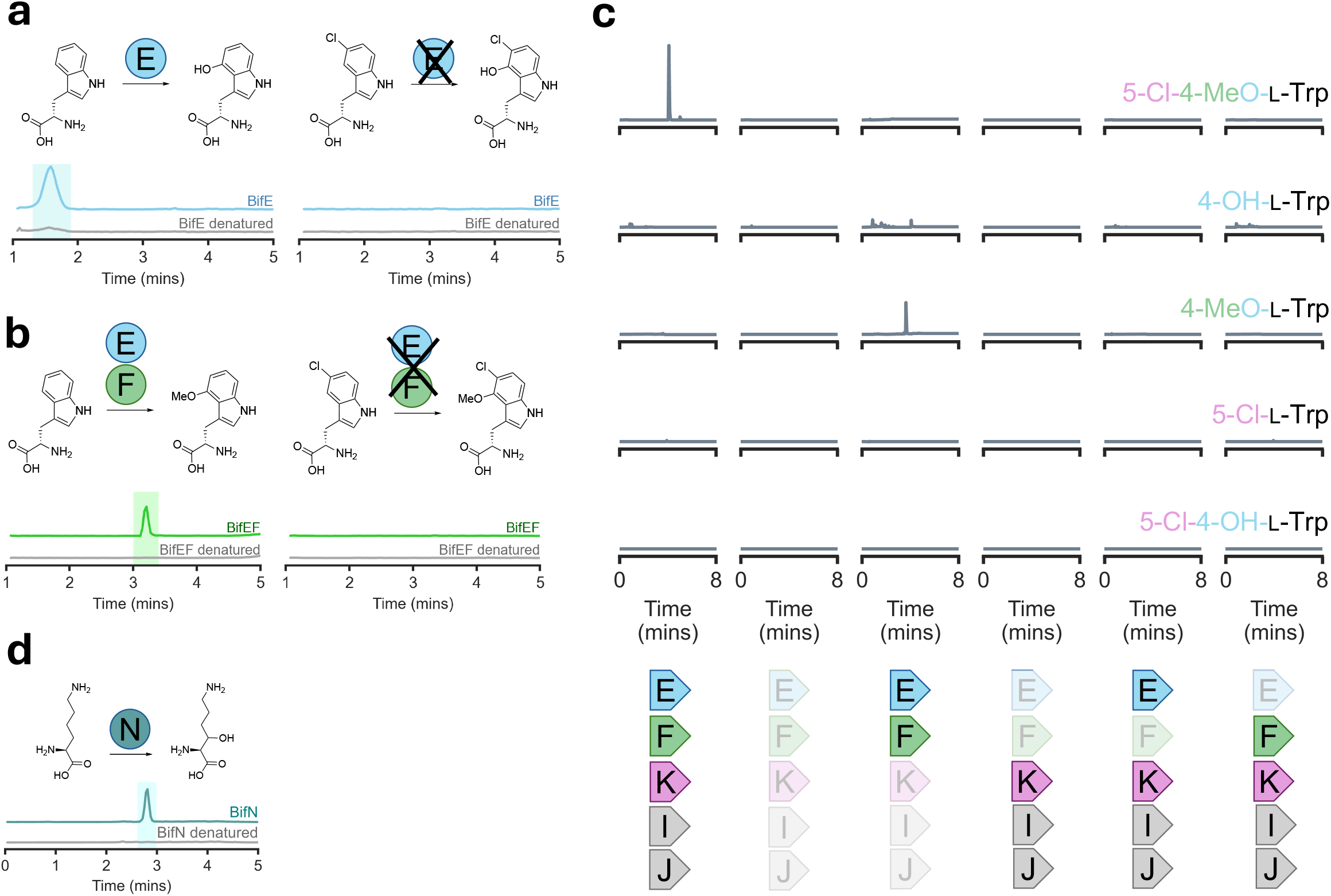
Reconstitution of 5-Cl-4-MeO-L-Trp and 3-OH-L-Lys biosynthesis. (a) *In vitro* reconstitution of 4-OH-L-Trp biosynthesis by BifE. Shown is the LCMS analysis of reactions mixtures of BifE or heat-inactivated BifE incubated with L-Trp (left) or 5-Cl-L-Trp (right); shown are EICs for the monoisotopic mass corresponding to the [M+H]^+^ adduct derived from 4-OH-L-Trp (*m/z* 221.1 ± 0.5) or 5-Cl-OH-L-Trp (*m/z* 255.1 ± 0.5), respectively. (b) *In vitro* reconstitution of 4-MeO-L-Trp biosynthesis in a one-pot reaction of BifEF or heat-inactivated BifEF with Trp (left) or 5-Cl-L-Trp (right); shown are EICs for the monoisotopic mass corresponding to the [M+H]^+^ adduct derived from 4-MeO-L-Trp (*m/z* 235.1 ± 0.5) or 5-Cl-4-MeO-L-Trp (*m/z* 269.1 ± 0.5), respectively. (c) *S. coelicolor* M1152 Heterologous expression of 5-Cl-4-OH-L-Trp biosynthetic genes (indicated by coloured circles; faded circles indicate absence of gene(s). Shown are EICs for the monoisotopic mass corresponding to the [M+H]^+^ adduct for the compound indicated. 5-Cl-4-MeO-L-Trp (*m/*z 269.0687 ± 0.01); 4-OH-L-Trp (*m/z* 255.0531 ± 0.01); 4-MeO-L-Trp (*m/z* 235.1077 ± 0.01); 5-Cl-L-Trp (*m/z* 239.0582 ± 0.01); 5-Cl-4-OH-L-Trp (*m/z* 254.0458 ± 0.01). (d) *In vitro* reconstitution of 3-OH-L-Lys biosynthesis by BifN or heat-inactivated BifN with L-Lys; the EIC corresponds to the [M+H]^+^ adduct for Fmoc-OH-L-Lys (*m/z* 385.2 ± 0.5). For (b) and (d) the X denotes that the indicated enzyme(s) do not perform the specified reaction.

**Figure 5.**
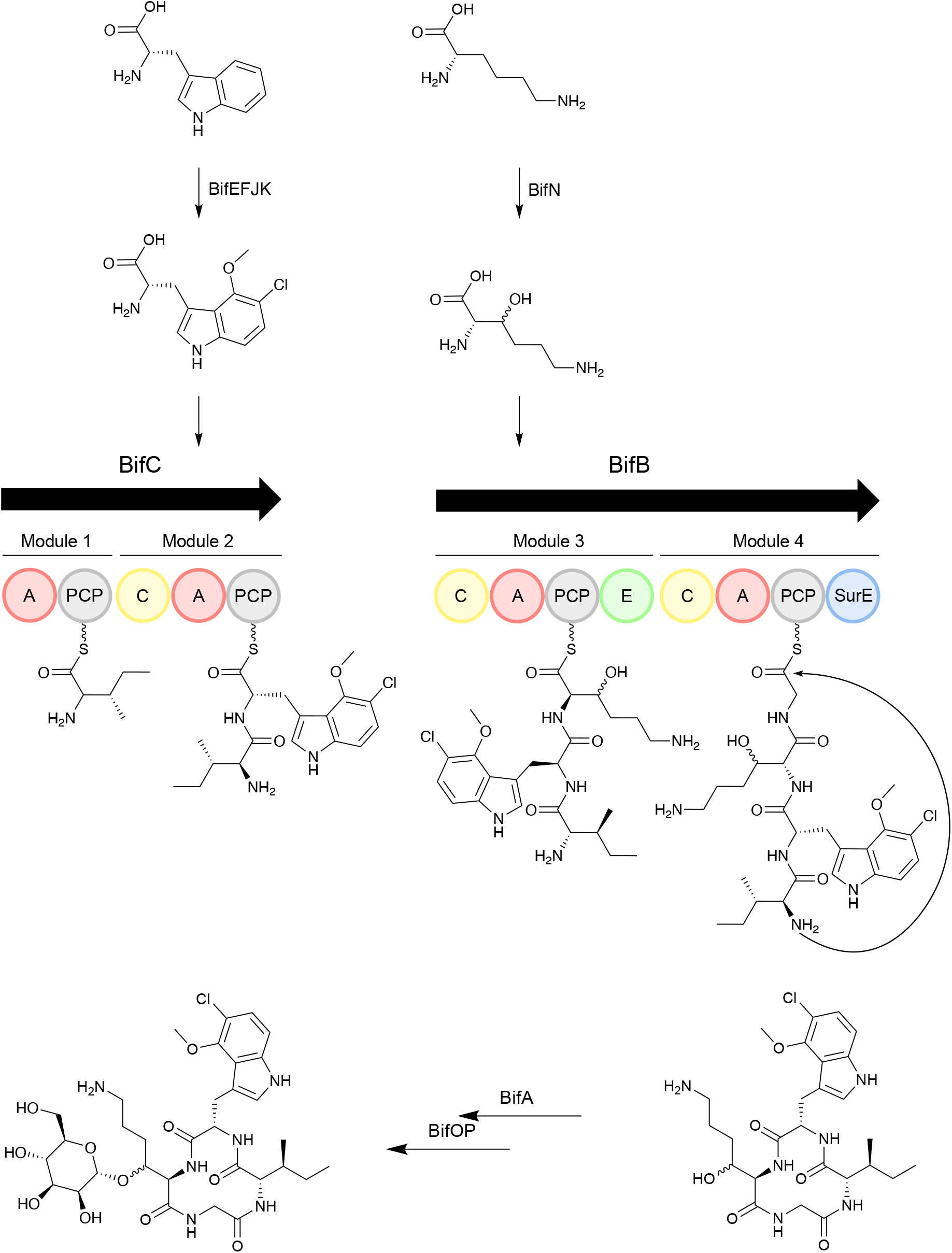
Proposed biosynthetic scheme for **1**

Through sequence analysis, BifK appears to be a conventional flavin-dependent Trp halogenase, which are well studied.^[19–21]^ To assess the halogenation activity of BifK, varying combinations of *bifEFJIK* were introduced into *Streptomyces coelicolor* M1152, which lacks the *bif* BGC. Metabolites were extracted from the resulting M1152 heterologous expression strains and analysed for the presence of **4** and relevant pathway intermediates by LCHRMS. The results are displayed in Figure 4c and indicate that expression of *bifEF* is sufficient for the heterologous production of 4-MeO-L-Trp and that expression of *bifEFIK* is essential for the heterologous production of **4**. The putative sodium/hydrogen exchanger encoded by *bifJ* was not essential for production of 4-MeO-L-Trp, at least not in *S. coelicolor* M1152. 4-OH-L-Trp was not observed, even in the absence of *bifE*, suggesting 4-OH-L-Trp is either unstable during cultivation or possibly consumed by another physiological process. Interestingly, 5-Cl-L-Trp was not observed in any experiment, suggesting that the BifK halogenase exhibits a tight specificity for 4-MeO-L-Trp. Taken together, these data support a model in which BifE hydroxylates C4 of L-Trp, the product of which is then methylated by BifF prior to halogenation at C5 by BifK and its cognate reductase partner BifI.

Hydroxylation of Lys C3 is presumed to be carried out by BifN, which is an α-KG-dependent amino acid hydroxylase, an enzyme type that is well-known within NRPS pathways^[22,23]^, and for which there are two subfamilies: those that work on-board with the NRPS assembly line (*i*.*e*., act on PCP*-S-*bound intermediates) and those that act on free amino acids.^[10,24]^ Our bioinformatics analysis of BifN suggested these two enzyme classes can be resolved phylogenetically (Figure S52), which led us to hypothesise that BifN acts upon free L-Lys. To experimentally test this hypothesis, (His)_6_-BifN was overproduced and purified from *E. coli* (Figures S48 and S51) and incubated with L-Lys alongside a heat-inactivated control. Reactions were derivatised with fluorenylmethyoxycarbonyl (Fmoc) to enable more reliable chromatography prior to analysis by LCMS. As anticipated, Fmoc-OH-L-Lys was detected for reaction mixtures containing (His)_6_-BifN but not heat-inactivated (His)_6_-BifN (Figure 4d). Taken together, these data suggest that BifN acts on free L-Lys and does not act on an NRPS bound substrate during biosynthesis. Although glidobactin harbours 4-OH-L-Lys^[25]^; to our knowledge a natural product harbouring 3-OH-L/D-Lys has not been observed previously. The stereochemistry of the hydroxyl group appended to Lys C3 is not known, however, only three L-Lys 3-hydroxylases have been characterised and they all catalyse the formation of (3*S*)-3-OH-L-Lys. ^[26,27]^ One of these hydroxylases, KDO1 from *Catenulispora acidiphila* DSM44928 (30% shared amino acid identity with BifN), is structurally characterised (PDB IDs 6F2A, 6F2B, 6F2E and 6F6J).^[27]^ An AlphaFold3 structural model of BifN was therefore generated and compared to 6F6J (KDO1 in complex with Fe^2+^, succinate and (3*S*)-3-OH-L-Lys), which revealed the active sites of the two enzymes are conserved and align well geometrically (Figure S53), suggesting BifN likely catalyses the formation of (3*S*)-3-OH-L-Lys. For **1**, the BifN catalysed 3-hydroxylation of L-Lys is critical because it is the attachment point for the D-mannose. It is interesting to note that only a trace amount of (**3**) was observed during fermentation of the producer strain, which suggests that either the BifB^A3^ has a substrate preference for 3-OH-L-Lys and/or that BifB^C4^ may have a gatekeeping function to minimise incorporation of L-Lys. It also suggests that the **3** may not be a substrate for the aglycon exporter.

### BifB^SurE^ is essential for production of 1

Intriguingly, the biffamycin A scaffold is predicted to be cyclised by an embedded SurE cyclase domain harboured within the terminal biosynthetic module of BifB. SurE is a promiscuous peptide cyclase belonging to the newly identified family of PBP-TEs^[4–6]^, which have thus far only been observed as standalone was essential for biffamycin A production, we deleted the BifB^SurE^ coding sequence in the Δ*bifO*-Δ*bifD1* strain using CRISPR-Cas9 editing and reintroduced the pBiff plasmid (Δ*bifO-bifD1*Δ*bifB*^*surE*^/pBiff). The resulting mutant strain and its parent were cultivated under biffamycin A production conditions and chemical extracts were analysed by LC-HRMS. As anticipated, biffamycin A was only detected in extract generated from the parental strain and absent from the Δ*bifO-bifD1*Δ*bifB*^*surE*^/pBiff strain, indicating BifB^SurE^ is essential for biffamycin A production (Figure S54).

### Proposed biosynthetic pathway for biffamycin A (1)

Structural elucidation of **1**, combined with enzymology and bioinformatics analyses of enzymes encoded by the *bif* BGC, led us to propose a biosynthetic pathway that begins with the modification of L-Trp and L-Lys prior to utilisation as substrate for an NRPS assembly line system. Specifically, C4 on the L-Trp indole ring is methoxylated by the concerted action of a heme-dependent Trp hydroxylase (BifE) and *O-*methyltransferase (BifF). Following methoxylation, C5 of 4-MeO-L-Trp is chlorinated by a chlorinase/flavin reductase pair (BifKI), possibly supplied with chlorine by the putative sodium-chlorine exchange protein, BifJ. Separately, L-Lys undergoes C3 hydroxylation catalysed by the α-KG-dependent hydroxylase, BifN. The cyclotetrapeptide aglycon is then assembled by the dimodular NRPSs BifC and BifB, plus BifH which is an MbtH-family protein that may enhance the activity of one or more of the NRPS adenylation domain(s). BifC possess two modules organised as follows: A1-PCP1-C2-A2-PCP2. The A1 domain initiates the chain by activating and loading L-Ile onto PCP1. A2 activates and loads 5-Cl-4-MeO-L-Trp onto PCP2, which is subsequently condensed with Ile by C2. BifB possesses two modules organised as follows: C3-A3-PCP3-E3-C4-A4-PCP4-SurE. A3 activates and loads 3-OH-L-Lys onto PCP3 where it is epimerised to the D-configuration by E3. C3 then catalyses the condensation of 3-OH-D-Lys with the dipeptide aminoacylthioester on PCP2. A4 activates and loads Gly onto PCP4, which is subsequently condensed with the tripeptide aminoacylthioester on PCP3 by C4 prior to macrocylisation of the resulting tetrapeptide aminoacylthioester by the embedded SurE cyclase domain of BifB. The cyclotetrapeptide aglycon is presumably exported by the MFS1 family exporter, BifA, which shares 49% amino acid identity with MppL, an MFS1 family transporter experimentally linked to export of the mannopeptimycin aglycon,^[29]^ where the 3-OH moiety of D-Lys within the aglycon is mannosylated by a membrane-bound mannosyl transferase (BifP), the substrate for which is provided by the BifO, a membrane-bound polyprenyl-P-hexose synthase in a manner analgous to that proposed for mannopeptimycin, ramoplanin and tecicoplanin.^[29-31]^

### The antibacterial activity of biffamycin A (1)

Glycopeptide antibiotics (GPAs) are a major antibacterial class that includes cell wall biosynthesis inhibitors such as vancomycin and teicoplanin.^[32,33]^ GPAs are relatively large, highly crosslinked natural products, and apart from type V GPAs, possess more than one carbohydrate moiety, sometimes composed of chains up to five sugars in length.^[34,35]^ In this respect, **1** is remarkable because it is approximately half the size of other GPAs (*i*.*e*., 726 Da compared to 1,449 Da for vancomycin), does not contain crosslinks, and to our knowledge, is the mallest naturally occurring glycosylated cyclotetrapeptide. To evaluate whether **1** might exhibit antibacterial activity, its minimum inhibitory concentrations (MIC) was determined for a small cross section of bacteria, including *Mycobacterium smegmatis* and clinical isolates of *S. aureus* resistant to methicillin (MRSA) or vancomycin (VRSA) as well as clinical isolates of *S. pyogenes, E. coli* and *P. aeruginosa*. The resulting data revealed that **1** possessed antibacterial activity against Gram-positive organisms with an MIC of 8 µg mL^-1^ for both MRSA and VRSA, 32 µg mL^-1^ for *S. pyogenes* ATCC19615, and 16 µg mL^-1^ for *M. smegmatis*, whereas **1** was inactive at the concentrations used against Gram-negative organisms tested (Table 1).

**Table 1.** Minimum inhibitory concentration^[a]^ of 1 and intermediates against microorganisms and human cells.

| Organism | Biffamycin A<br>($\mu\text{g mL}^{-1}$) | Biffamycin A aglycon<br>($\mu\text{g mL}^{-1}$) | des-hydroxy biffamycin<br>A aglycon ($\mu\text{g mL}^{-1}$) |
| --- | --- | --- | --- |
| **Gram-positive** |  |  |  |
| *Staphylococcus aureus* USA300 (MRSA) | 8 | 64 | > 128 |
| *Staphylococcus aureus* VRSA-5 (VRSA) | 8 | n.d. | n.d. |
| *Streptococcus pyogenes* ATCC19615 | 32 | n.d. | n.d. |
| *Mycobacterium smegmatis* ATCC101 | 16 | n.d. | n.d. |
| **Gram-negative** |  |  |  |
| *Escherichia coli* IMHA 659045 | > 128 | n.d. | n.d. |
| *Pseudomonas aeruginosa* PA14 | > 128 | n.d. | n.d. |
| **Mammalian cells** |  |  |  |
| Human embryonic kidney (HEK) 293 | > 128 | n.d. | n.d. |

<sup>[a]</sup> The MIC was tested by broth microdilution in triplicate. MRSA, methicillin-resistant *S. aureus*; VRSA, vancomycin-resistant *S. aureus*.

Interestingly, the biffamycin A aglycon (**6**) retained bioactivity against MRSA, albeit the MIC was 64 µg mL^-1^, eight times higher than that for **1**, while the *des-*hydroxy biffamycin A aglycon (**3**) was inactive at 128 µg mL^-1^. At the highest concentration tested (128 µg mL^-1^), **1** did not show cytotoxicity when incubated for 48 h with human cell line HEK293 (Table 1, Figure S55). The bioactivity of **1** against only Gram-positive bacteria and its ineffectiveness against Gram-negative bacteria is consistent with the profile of other GPAs to which Gram-negative bacteria possess intrinsic resistance because the molecules cannot cross the outer membrane.^[36]^ The mode of action of **1** remains to be determined, however given that it is not cytotoxic to HEK293 cells, it is tempting to speculate **1** may inhibit cell wall biosynthesis, like other GPAs.^[37]^ However, the ability of **1** to inhibit the growth of *S. aureus* VSRA-5, which harbours *vanA* and is highly resistant to vancomycin with an MIC >256µg mL^-1^ compared to an MIC of 2 µg mL^-1^ for susceptible strains^[35,38-40]^ suggests that **1** either has a mode of action different to vancomycin or that it circumvents VanA-mediated resistance.

## Conclusion

Antibiotic resistant poses a severe threat to global health. While most antibiotics are derived from microbial natural products, many if not most biosynthetic pathways are not expressed/productive in the laboratory. To overcome this challenge, we used promoter engineering to de-regulate the expression of a silent biosynthetic gene cluster, which enabled isolation and characterisation of its product – a novel glycopeptide antibiotic (GPA) we named biffamycin A. Biffamycin A contains two unprecedented chemical moieties, 3-hydroxy(α-D-mannoysl)-D-Lys and 5-chloro-4-methoxy L-Trp, whose biosynthesis we characterised through *in vitro* and *in vivo* reconstitution experiments. SurE/PBP-TE cyclases are emerging as biocatalysts to improve yield and diversity during chemical synthesis of cyclopeptide therapeutics^[41–43]^. A particular challenge is the cyclisation of tetrapeptides where current methods are inadequate^[44,45]^. BifB^SurE^ is the first member of the SurE/PBP-TE family whose native substrate is a tetrapeptide, which together with a recently characterised hexapeptide cyclase with tetrapeptide cyclase activity^[46]^, provides a foundation to generate diverse libraries of cyclotetrapeptides for drug discovery. Moreover, biffamycin A stands out as the smallest known member of its class, lacking the complex structural features typically associated with GPAs. Despite its simplicity, biffamycin A demonstrates potent activity against drug-resistant bacteria, including MRSA and VRSA, as well as the model organism, *M. smegmatis*. This discovery presents a unique opportunity to leverage the scaffold of biffamycin A for developing novel therapeutics against *M. tuberculosis*, a pathogen of critical priority for the World Health Organization.^[47]^

## Acknowledgements

We thank Stuart Warriner and Jeanine Williams for training and maintenance of instrumentation in the School of Chemistry and the JIC Metabolomics platform for excellent mass spectrometry support. We also thank Thomas Billinge for initial insights into BifN hydroxylation, Divya Thankachan and Asif Fazal for constructive discussions throughout the project, and Alex O’Neill for providing clinical isolates. **M.B**. was supported by a PhD studentship funded by the White Rose Mechanistic Biology Doctoral Training Programme funded by BBSRC (BB/T007222/1). **E.H**. and **B.W**. were supported by the BBSRC via the Institute Strategic Programme Grant ‘Harnessing Biosynthesis for Sustainable Food and Health’ (HBio) (BB/X01097X/1). **D.C.L.V**. was supported by a PhD studentship funded by the Wellcome Trust (215064/Z/18/Z). **J.C**. and **R.F.S**. were supported by a BBSRC responsive mode grant BB/T014962/1.

## Supplementary Information

Supplementary information includes the materials and methods, Tables S1-S9 and Figures S1-S59.

## Notes

### Competing Interest Statement

The authors have declared no competing interest.

