## Supplementary Information for "Discovery and biosynthesis of biffamycin A – a novel glycotetrapeptide antibiotic"

[b] Prof. Dr. M. E. Webb
Astbury Centre for Structural Molecular Biology, School of Chemistry, University of Leeds, Leeds, LS2 9JT (UK)

[c] Dr. E. S. Hems, Prof. Dr. B. Wilkinson
Centre for Microbial Interactions, Department of Molecular Microbiology, John Innes Centre, Norwich, NR4 7UH (UK)

[d] Dr. S. Nepogodiev
NMR platform, John Innes Centre, Norwich, NR4 7UH (UK)

**Contents**

**Materials and methods**
**Table S1.** Bioinformatic summary of biffamycin A BGC gene products (separate Excel file).
**Figure S1.** Analysis of adenylation domain active site residues with PARAS.
**Figure S2.** Clustal O alignment of the BifB-PCP-SurE di-domain and the SurE (PDB ID 6KSU).
**Figure S3.** Structural analysis of $BifB^{SurE}$.
**Figure S4.** Multisequence alignment of nonribosomal peptide synthetase epimerase (E) domains.
**Figure S5.** Deregulation of the biffamycin (*bif*) biosynthetic gene cluster.
**Figure S6.** Analysis of **1** and **2** production by Δ*bifO-bifD1*/pBiff when cultivated under amino acid supplementation.
**Figure S7.** $MS^2$ spectra of ion 727.3051 biffamycin A (**1**).
**Table S2.** Mapping of $MS^2$ spectra for biffamycin A (**1**).
**Figure S8.** $MS^2$ spectra of ion 713.2905 biffamycin B (**2**).
**Table S3.** Mapping of $MS^2$ spectra for biffamycin B (**2**).
**Figure S9.** $MS^2$ spectra of ion 565.2535 biffamycin A aglycone (**6**).
**Table S4.** Mapping of $MS^2$ spectra for biffamycin A aglycone (**6**).
**Figure S10.** $MS^2$ spectra of ion 549.2585, *des*-hydroxy biffamycin A aglycone (**3**).
**Table S5.** Mapping of $MS^2$ spectra for *des*-hydroxy biffamycin A aglycone (**3**).
**Figure S11.** $MS^2$ spectra of ion 786.3433, *N*-acetyl biffamycin (**7**).
**Table S6.** Mapping of $MS^2$ spectra for *N*-acetyl biffamycin A (**7**).
**Figure S12.** The sugar hydrolysed from biffamycin (**1**) was mannose.
**Table S7.** Resonance assignments in the $^1H$ and $^{13}C$ NMR spectra.
**Figure S13.** $^1H$ NMR of **3** (DMSO-$d_6$, 600 MHz, 298 K).
**Figure S14.** $^{13}C$ NMR of compound **3** (DMSO-$d_6$, 151 MHz, 298 K).
**Figure S15.** $^1H$-$^1H$ COSY spectrum of **3** (DMSO-$d_6$, 298 K).
**Figure S16.** 2D TOCSY spectrum of **3** (DMSO-$d_6$, 298 K).
**Figure S17.** $^1H$-$^{13}C$ HSQC-edited spectrum of **3** (DMSO-d6, 298 K).
**Figure S18.** $^1H$-$^{13}C$ HMBC spectrum of **3** (DMSO-d6, 298 K).
**Figure S19.** $^1H$-$^1H$ ROESY spectrum of **3** (DMSO-d6, 298 K).
**Figure S20.** $^1H$ NMR spectrum of **4** (DMSO-d6, 600 MHz, 298 K).
**Figure S21.** $^{13}C$ NMR spectrum of **4** (DMSO-d6, 151 MHz, 298 K).
**Figure S22.** $^1H$-$^1H$ COSY spectrum of **4** (DMSO-d6, 298 K).
**Figure S23.** $^1H$-$^{13}C$ HSQC-edited spectrum of **4** (DMSO-d6, 298 K).
**Figure S24.** $^1H$-$^{13}C$ HMBC spectrum of **4** (DMSO-d6, 298 K).
**Figure S25.** 1,1-ADEQUATE spectrum of **4** (DMSO-d6, 298 K).
**Figure S26.** $^1H$-$^1H$ ROESY spectrum of **4** (DMSO-$d_6$, 298 K).
**Figure S27.** $^1H$ NMR spectrum of compound **5** (DMSO-$d_6$, 600 MHz, 298 K).
**Figure S28.** $^{13}C$ NMR of compound **5** (DMSO-$d_6$, 151 MHz, 298 K).
**Figure S29.** $^1H$-$^1H$ COSY spectrum of **5** (DMSO-$d_6$, 298 K).
**Figure S30.** 2D TOCSY spectrum of **5** (DMSO-$d_6$, 298 K).
**Figure S31.** $^1H$-$^{13}C$ HSQC-edited spectrum of **5** (DMSO-$d_6$, 298 K).
**Figure S32.** $^1H$-$^{13}C$ HMBC spectrum of **5** (DMSO-$d_6$, 298 K).
**Figure S33.** $^1H$-$^1H$ ROSEY spectrum of **5** (DMSO-$d_6$, 298 K).
**Figure S34.** $^1H$ NMR of **6** (DMSO-$d_6$, 600 MHz, 298 K).
**Figure S35.** $^{13}C$ NMR of **6** (DMSO-$d_6$, 151 MHz, 298 K).
**Figure S36.** $^1H$-$^1H$ COSY spectrum of **6** (DMSO-$d_6$, 298 K).
**Figure S37.** 2D TOCSY NMR spectrum of **6** (DMSO-$d_6$, 298 K).
**Figure S38.** $^1H$-$^{13}C$ HSQC-edit spectrum of **6** (DMSO-$d_6$, 298 K).
**Figure S39.** $^1H$-$^{13}C$ HMBC spectrum of **6** (DMSO-$d_6$, 298 K).
**Figure S40.** $^1H$-$^1H$ ROESY spectrum of **6** (DMSO-$d_6$, 298 K).
**Figure S41.** $^1H$ NMR of **7** (DMSO-$d_6$, 600 MHz, 298 K).
**Figure S42.** $^{13}C$ NMR of **7** (DMSO-$d_6$, 151 MHz, 298 K).
**Figure S43.** $^1H$-$^1H$ COSY spectrum of **7** (DMSO-$d_6$, 298 K).
**Figure S44.** $^1H$-$^{13}C$ HSQC-edit spectrum of **7** (DMSO-$d_6$, 298 K).
**Figure S45.** $^1H$-$^{13}C$ HMBC spectrum of **7** (DMSO-$d_6$, 298 K).
**Figure S46.** $^1H$-$^1H$ ROESY spectrum of **7** (DMSO-$d_6$, 298 K).

**Figure S47.** Sections of coupled 2D HSQC spectra (DMSO-$d_6$ at 298 K) of (**A**) compound **7** and (**B**) methyl α-D-mannopyranoside containing cross-peaks of the mannose residue.
**Figure S48.** Purification and analysis of recombinant proteins.
**Figure S49**. Deconvoluted intact mass spectrometry for $(His)_6$-BifE.
**Figure S50.** Deconvoluted intact mass spectrometry for $(His)_6$-BifF.
**Figure S51.** Deconvoluted intact mass spectrometry for $(His)_6$-BifN.
**Figure S52.** Bioinformatic analysis of experimentally characterised α-ketoglutarate-dependent hydroxylases (αKHOs).
**Figure S53.** Structural analysis of BifN.
**Figure S54.** BifBSurE is required for biffamycin A production.
**Figure S55.** Biffamycin A is not cytotoxic against HEK293 cells.
**Figure S56.** Semi-synthesis of *N*-acetyl biffamycin A (**7**).
**Table S8.** Bacterial strains and plasmids used in this study.
**Table S9.** Oligonucleotides used in this study.

### METHODS

**Growth media, strains and other reagents.** *Escherichia coli* strains were cultivated using Lennox agar (LA) or broth (LB) and *Streptomyces* strains were propagated on mannitol-soya flour agar or TSB broth.[48] Recipes for agar media: SFM, ISP2, YEMETSB and MM are described at www.actinobase.org.[49] MM production broth was the same as MM but without agar and supplemented with 5 mM Ile and 10 mM NaCl. Culture media was supplemented with antibiotics as required at the following concentrations: apramycin (50 μg $mL^{-1}$), carbenicillin (100 μg $mL^{-1}$), hygromycin (75 μg $mL^{-1}$), kanamycin (50 μg $mL^{-1}$), nalidixic acid (25 μg $mL^{-1}$). Enzymes were purchased from New England Biolabs unless otherwise stated. Q5 DNA polymerase was used for all PCR reactions unless otherwise stated. The DNA constructs and bacterial strains used in this study are listed in Table S8. Oligonucleotides were purchased from Integrated DNA Technologies and are described in Table S9. Solvents used for extractions, purifications and LCMS analysis were purchased from Fisher Scientific (UK) or Sigma-Aldrich.

**Construction of *S. albidoflavus* S4 Δ5 Δ*bifO-bifD1*.** The *bifO-bifD1* was deleted from the chromosome of *S. albidoflavus* S4 using the ReDirect PCR targeting system.[50] CosBif was identified by screening a previously constructed *S. albidoflavus* S4 cosmid library[51] by PCR using primers RFS88 and RFS99. CosBif was insert-end sequenced using primers RFS184 and RFS185 and the resulting reads were mapped to the *S. albidoflavus* S4 genome[7] to establish the sequence of the cloned insert. A deletion cassette was PCR amplified from pHygP[52] using primers RFS764 and RFS778, and the resulting gel-purified and DpnI-restricted PCR product was used to delete *bifO-BifD1* from CosBif using recombineering with *E. coli* GB05-red.[53]The resulting mutagenised cosmid was electroporated into *E. coli* ET12567/pUZ8002[54] and subsequently mobilised to *S. albidoflavus* S4 Δ5 by conjugation as described previously.[48] Transconjugants were selected for hygromycin resistance and kanamycin sensitivity. The integrity of the mutant strain was verified by PCR using RFS766 and RFS779 and sequencing of the resulting PCR product.

**Construction of pBiff.** pBiff was constructed by five rounds of NEB HiFi Assembly. First, the *bifKJI* operon was PCR amplified from CosBif using RFS772 and RFS773 and assembled with NdeI/KpnI-restricted pSETermEp[55], to result in pBiff-temp-1. Second, pBiff-temp-1 was linearized with KpnI and assembled with two PCR fragments: (1) *rpsL(XC)*p amplified from pCRISPomyces-2[56] with RFS774 and RFS775; (2) *bifFE* amplified from CosBif using RFS776 and RFS777. The resulting plasmid was named pBiff-temp-2. Third, pBiff-temp-2 was linearized with SpeI and assembled with four PCR fragments: (1) *ermE**p amplified from pSETermEp[55] with RFS788 and RFS789; (2) *bifN* amplified from CosBif with RFS790 and RFS791; (3) *rpsL(XC)*p amplified from pCRISpomyces-2[56] with RFS792 and RFS793; (4) *bifH* amplified from CosBif with RFS794 and RFS795. The resulting plasmid was named pBiff-temp-3. Fourth, pBiff-temp-3 was assembled with *bifD1D2.* To achieve this, an Integrated DNA Technologies gBlock (Table S6) was designed to harbour *ermE**p[57] and the first 124bp of the *bifD2* permease to enable recoding of the endogenous leucyl TTA codon to CTC. pBiff-temp-3 was then linearized with EcoRI and assembled with two PCR fragments: (1) the above gBlock amplified with RFS800 and RFS801; (2) *bifD1D2* amplified with RFS786 and RFS787. The resulting plasmid was named pBiff-temp-4. Fifth, pBiff-temp-4 was linearised with EcoRI and was assembled with three PCR fragments: (1) *rpsL(XC)*p amplified from pCRISPomyces-2[56] with RFS802 and RFS803; (2) *bifO* amplified from CosBif with RFS804 and RFS854; (3) *bifP* amplified from CosBif with RFS855 and RFS856. The final plasmid was designated pBiff and its deduced nucleotide sequence is available at https://doi.org/10.6084/m9.figshare.27958413.v2.

**Heterologous production of Cl-MeO-Trp.** Circular Polymerase Extension Cloning[58] was used to systematically delete genes from the starting plasmid, pBiff-temp-2 from above. In brief, PCR fragments were generated from pBiff-temp-2 that when assembled omitted the target gene. To construct pBifFIJK, two PCR fragments were assembled from PCR fragments generated with MWB126 and MWB138, and MWB127 and MWB137; *bifKJI* is expressed from *ermE**p and *bifF* is expressed from *rpsL*(XL)p. To construct pBifEIJK, one PCR fragment was generated using MWB124 and MWB135 and assembled; *bifKJI* is expressed from *ermE**p and *bifE* is expressed from *rpsL*(XL)p. To construct pBifEF, two PCR fragments were assembled from PCR fragments generated with MWB126 and MWB136, and MWB127 and MWB135; *bifEF* is expressed from *rpsL*(XL)p.

**Deletion of the BifB$^{SurE}$ domain.** CRISPR/Cas9 editing with a homology-directed repair template was used to delete the nucleotides specifying the SurE domain within BifB. Briefly, oligonucleotides JEC225 and JEC226 were annealed and cloned into pCRISPomyces-2 using BbsI Golden Gate cloning as described.[56] A homology-directed repair template was PCR-amplified from genomic DNA using JEC221 and JEC222 (upstream arm) and JEC223 and JEC224 (downstream arm). The resulting PCR fragments were assembled with a XbaI-linearised pCRISPomyces-2 vector using NEB HiFi Assembly mix. The resulting plasmid was mobilized to *S. albidoflavus* S4 Δ5 Δ*bifO-bifD1* by conjugation as described and transconjugants were selected for apramycin resistance. This strain was passaged until the temperature-sensitive pCRISPomyces-2 plasmid was cured, which was identified phenotypically by apramycin sensitivity. The desired mutation was confirmed by PCR using oligonucleotides JEC245 and JEC246.

**Purification of hexahistidine tagged BifE, BifF and BifN.** Coding sequences for BifE, BifF and BifN were PCR amplified from *S. albidoflavus* S4 genomic DNA using RFS812 and RFS813 (BifE), DCLV179 and DCLV180 (BifF), and RFS814 and RFS815 (BifN) and cloned into pET28 at the NdeI and HindIII restriction sites using conventional cut and paste cloning with T4 DNA ligase. Buffers for lysis, washing and elution were as follows. Lysis buffer = 50 mM HEPES pH 8.0, 200 mM NaCl, 10 mM imidazole, cOmplete EDTA-free protease inhibitor, and DNAse I; wash buffer = 50 mM HEPES pH 8.0, 200 mM NaCl, 20 mM imidazole; elution buffer = 50 mM HEPES pH 8.0, 200 mM NaCl, 200 mM imidazole.

For BifE, an overnight culture of *E. coli* C41(DE3) harbouring pET28a-bifE was subcultured (1% v/v) into fresh LB medium (eight 2L flasks each harbouring 1L LB supplemented with kanamycin

(50 μg mL$^{-1}$)). Cultures were incubated at 37 ˚C with shaking. At $OD_{600nm}$ 0.2, cultures were supplemented with aminolevulinic acid in 1 M HEPES to a final concentration of 0.6 mM and incubation resumed. At $OD_{600nm}$ 0.6, production was induced with IPTG at a final concentration of 0.1 mM. Cultures were shifted to 16 ºC with shaking and incubated overnight. Cells were harvested by centrifugation at 5,000 x *g* for 15 min and resuspended in lysis buffer and subsequently lysed by sonication on ice (1 second on, 1 second off for 1.5 mins) twice. Lysate was clarified by centrifugation at 17,500 x *g* for 1 hour. Clarified lysate was subjected to gravity flow immobilisation metal affinity chromatography (IMAC) with 2 mL of Ni-NTA resin (Generon). The loaded resin was washed with wash buffer with regular sampling/analysis via Bradford reagent until no protein was detected in the flow-through. Bound protein was eluted from Ni-NTA resin with elution buffer and red coloured fractions (indicative of heme bound by $(His)_6$-BifE fractions were collected. Fractions were assessed by SDS-PAGE. Relevant elution fractions were pooled and concentrated with a 20 mL centrifugal filter with a 10 kDa molecular weight cut off and subject to size-exclusion chromatography on an ÄKTA Pure using a HiLoad Superdex S75 16/600 column. Red coloured fractions were assessed by SDS-PAGE and those containing $(His)_6$-BifE were pooled together, concentrated as above and snap-frozen with liquid nitrogen in 50 mM HEPES, 200 mM NaCl and 10% glycerol.

For BifF, an overnight culture of *E. coli* BL21(DE3) harbouring pET28a-bifF was subcultured (1% v/v) into fresh LB medium (four 2 L flasks each harbouring 0.5 L LB supplemented with kanamycin 50 μg mL$^{-1}$). $(His)_6$-BifF production was induced and purified as described above, but without gel filtration, concentrated as above and snap-frozen with liquid nitrogen in 50 mM HEPES, 200 mM NaCl and 10% glycerol.

For BifN, an overnight culture of *E. coli* BL21(DE3) harbouring pET28a-bifN was subcultured (1% v/v) into fresh LB medium (ten 2 L flasks each harbouring 1 L LB supplemented with kanamycin 50 μg mL$^{-1}$). $(His)_6$-BifN production was induced with 0.1 mM IPTG and cultivated at 16 ºC for 16 hrs whilst shaking at 180 rpm. Cells were harvested by centrifugation at 4,000 × *g* for 20 mins and resuspended in 50 mM HEPES pH 7.5, 200 mM NaCl and 20 mM imidazole supplemented with cOmplete protease inhibitor. Cells were lysed on ice by sonication at 40% power with 10 cycles of 10 s pulse followed by 20 s rest. The resulting lysate was clarified by centrifugation at 40,000 × *g* at 4 ºC for 30 mins. Cleared lysate was passed through a 0.45 μm syringe filter and loaded onto a 1 mL HisTrap column (GE Healthcare Life Sciences) using an ÄKTA Pure system. $(His)_6$-BifN was eluted using an imidazole gradient (20-200 mM). Fractions corresponding to high absorbance at 280 nm were pooled and analysed by SDS-PAGE. $(His)_6$-BifN was concentrated to 1 mL volume using a Vivaspin 20 centrifugal concentrator with a 10 kDa MWCO (Sartorius). $(His)_6$-BifN was then injected onto a HiLoad 16/600 superdex 75 pg column (GE Healthcare Life Sciences) equilibrated with 50 mM HEPES pH 7.5, 200 mM NaCl. Fractions were collected and those that eluted at the volume predicted for $(His)_6$-BifN (based on MW) were pooled and analysed by SDS-PAGE. The final $(His)_6$-BifN solution was supplemented with glycerol to 20% and stored at -80°C.

**BifE, BifEF and BifN activity assays.** BifE reactions consisted of a 50 µL volume composed of 20 mM ascorbate, 2 mM Trp or 5-Cl-Trp and 50 µM $(His)_6$-BifE in 100 mM HEPES. Reactions were incubated for 16h at 30 ºC without shaking. Negative control reactions consisted of 50 µM $(His)_6$-BifE heat-denatured at 95 ºC for 15 mins. Reactions were quenched with 150 µL methanol and incubated on ice for 10 mins. Any debris was removed by centrifugation at 16,000 × *g* for 10 mins. The clarified methanolic supernatant was diluted with an equal volume of water and 25 µL of sample was accessed by LC-MS (see below).

One-pot BifEF methoxylation assays consisted of a 50 µL volume composed of 20 mM ascorbate, 2 mM Trp or 5-Cl-Trp, 2 mM S-adenosylmethionine, 50 µM $(His)_6$-BifE and 0.1 µM $(His)_6$-BifF. Reactions were incubated as above and negative control reactions consisted of 0.1 µM $(His)_6$-BifF heat-denatured at 95 ºC for 15 mins. Reactions were quenched and analysed by LC-MS. For LC-MS, sample was injected into an Agilent Infinity II 1290 LC-MS system coupled with an Agilent InfinityLab Poroshell 120 EC-C18 (2.1 x 50 mm, 2.7 µm) column using the following gradient: solvent A, 0.1 % formic acid in water (v/v); solvent B, 0.1% formic acid in methanol (v/v); flow rate: 0.9 mL $min^{-1}$; T = 0 min, 2% B; T = 1.02 min, 2% B; T = 3.16 min, 98% B; T = 3.66 mins, 98% B; T = 4.26 min, 2% B; T = 7.00 min, 2% B.

The BifN hydroxylation assay consisted of 0.5 mM L-Lys, 0.5 mM ascorbic acid, 0.5 mM α-ketoglutaric acid and 0.5 mM $(NH_4)_2Fe(SO_4)_2 \cdot 6H_2O$ in 50 mM HEPES buffer pH 8.0 in a 3 mL volume. Reactions were started by addition of 1 µM $(His)_6$-BifN or heat-inactivated $(His)_6$-BifN and incubated at 30˚C for 2 h and quenched by freezing at -20 ºC. 0.5 mL of quenched reaction was derivatised by addition of 90 µL of fluorenylmethyloxycarbonyl (Fmoc) saturated with aqueous $NaHCO_3$, and then 1.25 mM *N*-9H-Fluoren-9-ylmethoxycarbonyloxy)succinimide (Fmoc-OSu) in 50% (v/v) acetonitrile, final concentration. Samples were dried under reduced pressure in a centrifugal evaporator and then resuspended in 0.5 mL deionised $H_2O$. Fmoc-derivatised sample was applied to a 1 mL / 30 mg Strata XL cartridge, washed with 3 mL of water and was sequentially eluted with 0.25 mL 50 % acetonitrile followed by 0.25 mL 100 % acetonitrile. Pooled eluent was diluted with 0.5 mL $H_2O$, and 8 µL of sample was injected into an Agilent Infinity II 1290 LC-MS system coupled with an Agilent InfinityLab Poroshell 120 EC-C18 (2.1 x 50 mm, 2.7 µm) column using the following gradient: solvent A, 0.1% formic acid in water (v/v); solvent B, 0.1% formic acid in methanol (v/v); flow rate: 0.9 mL $min^{-1}$; T = 0 min, 2% B; T = 0.22 min, 2% B; T = 2.36 min, 98% B; T = 2.86 mins, 98% B; T = 2.96 min, 2% B; T = 5.79 min, 2% B.

**AlphaFold analyses.** Structural models for BifN and the BifB-PCP-SurE-didomain were generated using the web implementation of AlphaFold3 available at https://alphafoldserver.com/. The first of five resulting structures was aligned with 6F6J (BifN)[27] or 6KSU ($BifB^{SurE}$)[43] using the align function in PyMol 2.5.8. To achieve a more informative structural comparison between $BifB^{SurE}$ and 6KSU, the single polypeptide chain was broken so that the β-lactamase domain and lipocalin domains comprised separate chains.

**Biffamycin A bioactivity assays. 1**, **3**, and **6** were dissolved in aqueous 10 mM HCl at 0.1 mg mL$^{-1}$, concentrated to apparent dryness under vacuum and freeze dried. **1** dissolved easily into water with no precipitate observed. The minimum inhibitory concentration (MIC) of **1** was determined for *S. aureus* USA300, *S. aureus* VRSA-5, *E. coli* IMHA 659045, *M. smegmatis* ATCC 101, *P. aeruginosa* PA14 and *S. pyogenes* ATCC 19615 according to CLSI standards.[59] MIC determination was carried out in triplicate and the mode MIC is reported. **1** was prepared to 1.28 mg mL$^{-1}$ in sterile water. A two-fold dilution series was generated from 1.28 mg mL$^{-1}$ to 0.025 mg mL$^{-1}$ in a 96-well plate by dispensing 10 µL per well. Unless otherwise stated, single colonies were used to inoculate Mueller-Hinton Broth 2 (MHB2, Millipore), which was incubated at 37 °C with shaking. For *M. smegmatis*, a single colony was spread plated on Mueller-Hinton Agar 2 (MHA2, MHB2 with 1.5 % (w/v) agar), grown at 37 °C for 24 h and cells harvested with MHB2 and a sterile cotton bud to create a cell suspension. For *S. pyogenes*, cultures were grown in MHB2-F (MHB2 with 5 % (v/v) lysed horse blood and 20 µg mL$^{-1}$ β-nicotinamide adenine dinucleotide). Cultures were diluted to $OD_{600nm}$ 0.1 in MHB2, and then diluted again to 5 × 10$^{5}$ CFU mL$^{-1}$ in MHB2 (MHB2-F for *S. pyogenes*). Appropriate dilutions were determined by calculating CFU mL$^{-1}$ at $OD_{600nm}$ 0.1 by the Miles and Misra method. In short, a 10-fold dilution series was generated from cultures at $OD_{600nm}$ 0.1 and 20 µL from 10$^{-2}$ to 10$^{-9}$ was spotted onto MHA2 and grown at 37°C till colonies appeared. CFU mL$^{-1}$ was calculated for spots containing 2-20 CFUs. 90 µL of inoculum was added to each well, thereby diluting compound concentrations 10-fold to a working concentration range from 128 µg mL$^{-1}$ to 0.25 µg mL$^{-1}$. A compound negative control, and an uninoculated media control were also included for each row. 96-well plates were sealed with Parafilm and incubated at 37 °C with agitation for 24 h. Compound MIC was determined at the compound concentration corresponding to no visible growth by eye. The mode MIC from three replicates was reported. For each 96-well plate, culture purity was determined by streaking for single colonies on MHA2 (MHB2-F for *S. pyogenes*) from the well corresponding to no compound. Plates were incubated at 37 °C overnight, aside from *M. smegmatis*, which was incubated for 36 h.

**Human embryonic kidney (HEK) 293 cell cytotoxicity assays.** HEK293 cells (26,000 cells per well) were seeded onto a 96-well microtiter plate and cultivated in Dulbecco's Modified Eagle's Medium (90 µL) overnight. After seeding, cells were challenged with either biffamycin A (128 µg mL$^{-1}$), puromycin (1 µg mL$^{-1}$) or empty vehicle (sterile deionised water) by applying 10 µL of a 10x solution for each compound. Four replicates were carried out for each treatment. After 48 h of treatment, culture medium was removed, and cells were washed with PBS prior to being fixed with 100 µL of 100% methanol for 10 min. After fixing, methanol was removed, cells were air-dried and treated with 100 µL of crystal violet solution (0.5% w/v in 20% methanol) and incubated for 20 min. Excess crystal violet stain was washed away with water and stained cells were air-dried. Finally, crystal violet retained by cells was solubilised in 100 µL of 10% acetic acid with agitation for 10 min and quantified spectrophotometrically at 590 nm.

**Fermentation of *S. albidoflavus* S4 Δ5Δ*bifO-bifD1*/pBiff and purification of biffamycin A and intermediates.** Seed-cultures were set up in spring-loaded 250 mL Erlenmeyer flasks with 100 mL of YEME-TSB (containing 25 μg $mL^{-1}$ nalidixic acid and 20 μg $mL^{-1}$ cycloheximide) and inoculated with 25 μL of *S. albidoflavus* S4 Δ5Δ*bifO-bifD1*/pBiff spore stock. Following 3 days of incubation at 30 ºC with shaking at 180 rpm, cells were harvested by centrifugation at 4,000 × *g* for 15 min and washed in 50 mL sterile PBS. Eight spring-loaded 2 L Erlenmeyer flasks containing 1 L of MM supplemented with 5 mM isoleucine were inoculated with ~6.25 g of wet cells per flask. Cultures were incubated for 3.5 days at 30 ºC with shaking at 180 rpm. The resultant cultures were pooled, and clarified by centrifugation at 10,500 × *g* for 20 min. The cell pellet was extracted with 2 volumes of methanol, and the resulting methanolic extract was filtered by gravity flow through grade 413 filter paper to remove large mycelial fragments prior to concentration under reduced pressure. The unextracted culture supernatant was first filtered through grade 413 filter paper to remove mycelial fragments and further clarified by filtration through sand (50-70 mesh white quartz) filtration. Following the addition of acetonitrile (3 % v/v), a total volume of 4 L of the filtered culture supernatant was passed through a Biotage Sfär C18 100 Å 30 μm 240 g cartridge pre-equilibrated in 5 % acetonitrile with 0.1 % formic acid. Following this, the cartridge was eluted using the following gradient: solvent A water with 0.1 % formic acid, solvent B acetonitrile with 0.1 % formic acid; flow rate 50 mL/min; elution started from 5 % B for 2 column volumes (CV); then gradient to 100 % B over 10 CV; and then holding at 100 % B for 2 CV. UV monitoring was at 226 nm and 275 nm. This resulted in two major peaks: the first contained predominantly **4** and the second containing **5**. LCMS analysis of fractions showed that **1** and **3** were spread across multiple fractions eluting between 5.5 CV – 8 CV, and these were pooled for purification by preparative HPLC (see below). The process was repeated on the remaining supernatant and fractions containing purified **1** were pooled.

For preparative HPLC, **1**- and **2**-**4**-containing Biotage chromatography fractions, as well as **6** from the cell pellet methanol extract, were purified by preparative HPLC on an Agilent 1260 Infinity II system fitted with a Phenomenex Kinetex XB-$C_{18}$ 100 Å 250 mm × 21.2 mm column over the following gradient: solvent A, 0.1% formic acid in water (v/v); solvent B, 0.1% formic acid in acetonitrile (v/v); flow rate, 20 mL $min^{-1}$; T = 0 min, 10 % B; T = 4 min, 10% B; T = 16 min, 75% B, T = 16.1 min, 98% B; T = 18.3 min, 98% B; T = 19 min, 10% B; T = 20 min, 10% B. The tetrapeptides **1**, **3** and **6** required an additional purification step by preparative HPLC on a Dionex Ultimate 3000 system fitted with a Phenomenex Kinetex XB-$C_{18}$ 100 Å 250 mm × 21.2 mm column over the following gradient: solvent A, 50 mM ammonium formate in water (w/v);, solvent B, acetonitrile, flow rate, 20 mL $min^{-1}$; T = 0 min, 5 % B; T = 3 min, 5% B; T = 3.5 min, 25% B, T = 25 min, 60% B; T = 25.5 min, 80% B; T = 29 min, 80% B; T = 30 min, 5% B; T = 32 min, 5% B; UV monitoring at 226 nm.

**Initial identification of biffamycins and their production under amino acid supplementation.** ~$10^6$ *S. albidoflavus* S4 Δ5Δ*bifO-bifD1*/pBiff spores were used to seed a lawn atop an MM agar plate supplemented with 5 mM of Ile, Val, or Leu or no supplement. Following 7 days of incubation at 30 ºC, agar was cut into 1 $cm^2$ blocks and extracted overnight at room temperature with 30 mL of 100% methanol. Mycelial fragments were removed from methanolic extract by filtration through grade 413 filter paper. The filtrate was concentrated to dryness under reduced pressure and resuspended in 5 mL of 100% methanol. Prior to analysis, particulate was sedimented by centrifugation at 13,500 × *g* for 10 mins and 2 µL of the supernatant was analysed by LCHRMS. For initial identification of biffamycins displayed in Fig. 2, LCHRMS was performed using a Bruker Maxis Impact II coupled with a Dionex Ultimate 3000 UHPLC with a Waters Acquity Premier VanGuard FIT CSH $C_{18}$ column (2.1mm × 1000 mm, 1.7 µm) using the following gradient: solvent A, 0.1% sodium formate in water (v/v); solvent B: 0.1% formic acid in acetonitrile (v/v); flow rate 0.7 mL $min^{-1}$; T = 0 min, 1% B; T = 1.5 min, 1%B; T = 2.5 min, 15% B; T = 3.0 min, 30% B; T = 5.0 min, 95% B; T = 6.0 min, 95% B; T = 6.5 min 1% B; T = 6.6 min, 1% B. The sample was ionised using a VIPHESI electrospray ionisation source and analysed in positive mode over the *m/z* range 50-2000. The instrument was calibrated with an infusion of 5 mM sodium formate solution and the resulting data was analysed using Bruker DataAnalysis 4.0. For analysis chemical extracts from supplemented cultures displayed in Figure S3, 2 µL of the supernatant was injected into a Thermo Scientific Orbitrap Exploris 250 mass spectrometer coupled with a Thermo Scientific Vanquish HPLC system. Metabolites were separated with a Hypersil GOLD $C_{18}$ column system (2.1 × 150 mm, 1.9 µm) using the following gradient: solvent A, 0.1% formic acid in water (v/v); solvent B, 0.1% formic acid in acetonitrile (v/v); flow rate 0.3 mL $min^{-1}$. T = 0 min, 1 % B; T = 1 min = 1 % B; T = 5 min, 40 % B; T = 6 min, 95 % B; T = 10 min, 95 % B; T = 11 min, 1 % B; T = 15 min, 1 % B. Mass measurement used full scan mode resolution of 60000, a *m/z* range of 80-800. The maximum injection time was set automatically by the software. MS2 selection was based on signal intensity of greater that 5E5 counts and charge state of 1-3. Dynamic exclusion of 10 s was used. Ions selected for MS2 were fragmented in the HCD cell using a relative collision energy of 30 % and measured with a resolution of 15000.

**LC-HRMS$^2$ analysis.** LCMS$^2$ analysis was performed using a Thermo QExactive LCMS instrument on a Kinetex $C_{18}$ column (50 × 2.1 mm, 1.7 µm). LCMS method, mobile phase A: water with 0.1% (v/v) formic acid; mobile phase B: acetonitrile with 0.1% (v/v) formic acid; flow rate 0.5 mL $min^{-1}$; injection volume 10 µL; elution gradient: *T* = 0 min, 5% B; *T* = 6 min, 95% B; *T* = 7.7 min, 95% B; *T* = 7.9 min, 5% B; *T* = 9 min, 5% B. The sample was analysed in positive mode over the *m*/*z* range 150–2000 with a resolution of 70 000. The spray voltage was set to 3000 V, and the capillary temperature was 350 °C. The sheaf gas was set to 35, and the auxiliary gas was set to 10. Data dependent MS$^2$ with 17 500 resolution and an isolation window of 4.0 *m*/*z* and an isolation offset of 1.0 *m*/*z* was employed with normalized collision energies of 20, 40, and 60 %. The instrument was calibrated according to the manufacturer's instructions, and the LCMS$^2$ data was analyzed using Thermo Scientific FreeStyle 1.7 software.

**Carbohydrate (HPAEC-PAD) Analysis.** Biffamycin (**1**, 50 µg) was sealed in a tube containing aqueous trifluoroacetic acid (TFA; 1.0 M, 1 mL) and heated to 105 °C overnight. The sample was then dried under vacuum using a GeneVac EZ-2 Elite HCl compatible evaporator. The residue was then dissolved in water/methanol (95:5, 1 mL) and passed through a C18 solid phase extraction cartridge (Waters, Sep-Pak Plus Short 360 mg). The cartridge was washed with water/methanol (95:5, 2 mL), and the eluted solvent was combined, dried and resuspended in water (150 µL). Carbohydrate analysis was performed by high performance anion exchange chromatography with pulsed amperometric detection (HPAEC-PAD) on a Dionex ICS-5000 system using a CarboPac PA20 (3 × 150 mm) analytical column coupled to a CarboPac PA20 (3 × 30 mm) guard column. For HPAEC-PAD analyses the following conditions were used: flow rate 0.25 mL $min^{-1}$; injection volume 5 µL; mobile phase A: 7.8 mM NaOH; mobile phase B: 156 mM NaOH with 100 mM AcONa; elution gradient: T = 0 min, 0% B; T = 30 min, 0% B; T = 33 min, 100% B; T = 55 min, 100% B; T = 58 min, 0% B; T = 72 min, 0% B. Peaks were identified by comparison with standards for hexoses of D-glucose, D-galactose, and D-mannose.

**Nuclear magnetic resonance (NMR) spectroscopy.** NMR spectra (1D and 2D) were recorded in DMSO-$d_6$ at 298 K on a Bruker Neo 600 MHz spectrometer equipped with 5 mm TCI CryoProbe. Two-dimensional $^1H$–$^1H$-COSY, $^1H$–$^{13}C$-HSQCed, HMBC, and ROESY experiments were performed using standard pulse sequences from the Bruker Topspin library. Data were processed using Topspin 4.1.4 and MestReNova 15.0.1 software, and spectra were calibrated to the TMS signals. ROESY spectra for compounds **5** and **6** were recorded on a Bruker Avance 400 MHz spectrometer equipped with a 5mm broadband probe.

**Semi-synthesis of *N*-acetyl biffamycin A (7).** The synthesis of **7** is summarised in **Figure S44.** Briefly, to synthesise **7,** $Ac_2O$ (10 µL, 10.8 mg, 0.11 mmol) was added to a solution of **1** (1.7 mg, 2.3 µmol) in anhydrous DMSO (1 mL) under argon and stirred at room temperature overnight. The solvent and all volatile components were removed *in vacuo*, and the resulting residue was stirred in a mixture of MeOH/$H_2O$/$Et_3N$ (5:2:1, 800 µL) at room temperature for 2 h. Next the reaction mixture was dried *in vacuo* and purified by preparative HPLC on a Dionex Ultimate 3000 system fitted with a Phenomenex Kinetex XB-$C_{18}$ 100 Å 250 mm × 10.0 mm column over the following gradient: solvent A, water + 0.1% (v/v) formic acid, solvent B, acetonitrile + 0.1% (v/v) formic acid, flow rate, 4 mL $min^{-1}$; T = 0 min, 10 % B; T = 5 min, 10% B; T = 7.5 min, 30% B, T = 47.5 min, 80% B; T = 50 min, 98% B; T = 60 min, 98% B; T = 62.5 min, 10% B; T = 72.5 min, 10% B; UV monitoring at 270 nm. The major UV active fraction was collected and dried *in vacuo* to give **7** (0.88 mg, 55%) as a white powder; $[\alpha]_D$ = -48° (*c* 0.1, DMSO); UV (DAD) $\lambda_{max}$ 226, 275 nm; $^1H$ and $^{13}C$ NMR data in DMSO-$d_6$, Table S6; HRESIMS *m/z* 786.3433 $[M+NH_4]^+$ (calc. for $C_{34}H_{53}ClN_7O_{12}^+$ 786.3435, Δ = -0.3 ppm).

**Compounds characterised in this study:**

**Biffamycin A (1)**

Brown powder; UV (DAD) $\lambda_{max}$ 226, 275 nm; $[\alpha]_D$ = -19° (*c* 0.1, 25 mM hydrochloric acid) HRESIMS *m/z* 737.3051 $[M+H]^+$ (calc. for $C_{32}H_{48}ClN_6O_{11}^+$ 727.3064, Δ = -1.8 ppm).

**Biffamycin B (2)**

Brown powder; HRESIMS *m/z* 713.2905 $[M+H]^+$ (calc. for $C_{31}H_{46}ClN_6O_{11}^+$ 713.2908, Δ = -0.4 ppm).

***Des*-hydroxy biffamycin A aglycon (3)**

Brown powder; UV (DAD) $\lambda_{max}$ 226, 275 nm; $^1H$ and $^{13}C$ NMR data in DMSO-$d_6$, **[Figures S13-19 and Table S7]**; HRESIMS *m/z* 549.2573 $[M+H]^+$ (calc. for $C_{26}H_{38}ClN_6O_5^+$ 549.2587, Δ = -2.5 ppm).

**5-Chloro-4-methoxy tryptophan (4)**

Brown powder; UV (DAD) $\lambda_{max}$ 226, 275 nm; $[\alpha]_D$ = -35° (*c* 0.1, DMSO); $^1H$ and $^{13}C$ NMR data in DMSO-$d_6$, **[Figures S20-26 and Table S7]**; HRESIMS *m/z* 269.0682 $[M+H]^+$ (calc. for $C_{12}H_{14}ClN_2O_3^+$ 269.0687, Δ = -1.9 ppm).

***N*-Acetyl-5-chloro-4-methoxy tryptophan (5)**

Brown powder; UV (DAD) $\lambda_{max}$ 226, 275 nm; $[\alpha]_D$ = -15° (*c* 0.1, DMSO); $^1H$ and $^{13}C$ NMR data in DMSO-$d_6$, **[Figures S27-33 and Table S7]**; HRESIMS *m/z* 311.0787 $[M+H]^+$ (calc. for $C_{14}H_{16}ClN_2O_4^+$ 311.0793, Δ = -1.9 ppm).

**Biffamycin A aglycon (6)**

Brown powder; UV (DAD) $\lambda_{max}$ 226, 275 nm; $^1H$ and $^{13}C$ NMR data in DMSO-$d_6$, **[Figures S34-40 and Table S7]**; HRESIMS *m/z* 565.2519 $[M+H]^+$ (calc. for $C_{26}H_{38}ClN_6O_6^+$ 565.2536, Δ = -3.0 ppm).

***N*-Acetyl biffamycin A (7)**

Brown powder; UV (DAD) $\lambda_{max}$ 226, 275 nm; $[\alpha]_D$ -48° (*c* 0.1, DMSO); $^1H$ and $^{13}C$ NMR data in DMSO-$d_6$, **[Figures S41-46 and Table S7]**; HRESIMS *m/z* 786.3433 $[M+NH_4]^+$ (calc. for $C_{34}H_{53}ClN_7O_{12}^+$ 786.3435, Δ = -0.3 ppm).

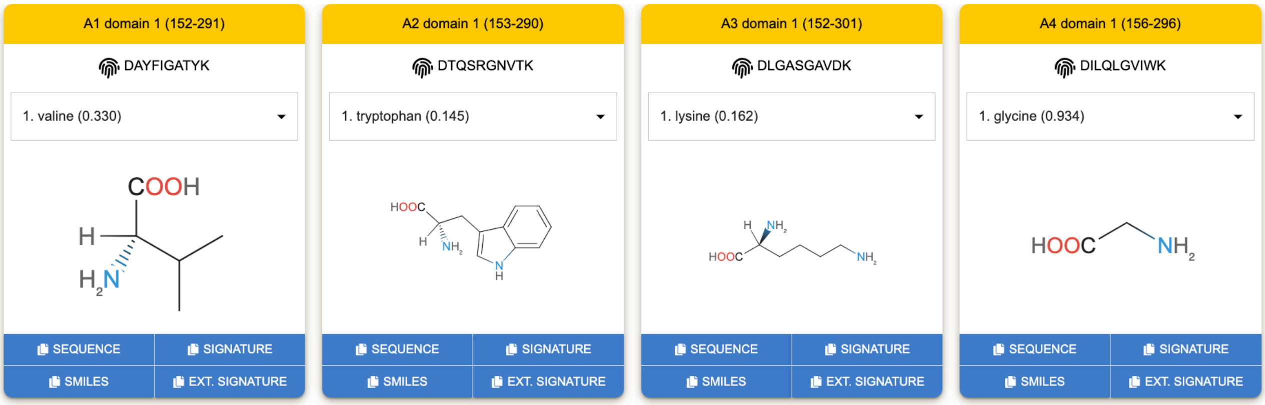


**Figure S1.** Analysis of adenylation domain active site residues with PARAS. The four adenylation domain sequences (BifC-A1, BifC-A2, BifB-A3, BifB-A4) were identified by antiSMASH v6.1.1[14] and analysed by PARAS[15] using the 'common substrates' setting.

```
BifB_PCP_SurE      TALAGVWAEVLGLPVESIGAHDDFFALGGDSVLCVQVAYRARRAGWWLVPRDLFSHPTVA      60
SurE_6KSU          ------------------------------------------------------------      0

BifB_PCP_SurE      LLAAVVRQEDTPVVPASAPSAPIASPTAVVPRPEVLADTGEPALGHTFLADRLAALARAH      120
SurE_6KSU          --------------------------------------MGAEGAERDAVGALFEELVREH      22
                                                          *  .  :  :.  :  *.* *

BifB_PCP_SurE      RVPGAQLAVRHAGGLTVVETGERVAGSGLPVTPDTAFPLASLTKPVTALAVRLLASDGRL      180
SurE_6KSU          RVTGAQLSVYRDGALSEYATGLASVRTGEPVTPRTGFPFGSVTKFLTAELVMQFVCDGDL      82
                   ** ****:* : *.*:   **   . :* **** *.**:.*:** :**  *  :..** *

BifB_PCP_SurE      DLDAPVGAYLPELSAA--SPAGRLTAYQLLTHTGGLAAAVDEREPSRDRRHWVERHCAQA      238
SurE_6KSU          DLDDPLAGLLPDLGRAAGPPLGTATVRQLLSHTAGVVDSIEYDEMRGPSYRRFAAACARQ      142
                   *** *:.. **:*. *   * *  *. ***:**.*:. :::  *      : .   **:

BifB_PCP_SurE      ALPHPPGTVFSYSDIGYVLLGRVVEVLTGTDWRTAVESALLRPLGITAAYQGQYG-----      293
SurE_6KSU          PALFPPGLAFSYSNTGYCLLGAVIEAASGMDWWTAMDSCLLRPLGIEPAFLHDPRPGQGG      202
                      .*** .****: ** *** *:*. :* ** **::*.*******  *:  :

BifB_PCP_SurE      -DRPAAHGHVVGD---RVLPVPEQSFSVLEEPATALAGNAADLVRLAGVHDRT-ATGGPL      348
SurE_6KSU          AARPVAEGHALRAGGERAEHV-DHMASLSLAAAGGLVGSATDLVTAARPHLADRKTFAQH      261
                     **.*.**.:     *.  * ::  *:    * .*.*.*:***  *  *     * .

BifB_PCP_SurE      DRAAADALLVDRTGDLRVGPFGLADGWGLGWAVHRGAGGDWNGHDGSGDGTWAHLRFDPA      408
SurE_6KSU          DLLPEDAVLAMRTCVPDAEPFGLADGWGLGLMRHGTGDGAWYGHDGAVGGASCNLRIHPD      321
                   *    **:*. **    . ***********   *  ..* * ****: .*: .:**:.*

BifB_PCP_SurE      TGTAVALVTNASTGAALWEALLAEIATAGLDVPHHSPQSPAGPLAPAADQAACAGSYANG      468
SurE_6KSU          RSLALALTANSTAGPKLWEALVARLPEAGLDVGHYALPVPD--SAPLAPDAGHLGTYANG      379
                    . *:**.:*:::*  *****:*.:  ***** *::   *    ** * :*.  *:****

BifB_PCP_SurE      DWVCLVEEGPEGIALSVGGG-PRSALTYYPDLRF---TSVGHGVPPTGRFLRDPDTGQVT      524
SurE_6KSU          DLELMVTHDAAGDLFLTRESYSDYRLSLHEDDLFVARSGEPGALPITGRFVREHPAGPVA      439
                   *   :* ..  *  : .  .     *: : *  *   :.   .:* ****:*:  :*  *:

BifB_PCP_SurE      LLQLAGRLLRRTGSPAR   541
SurE_6KSU          LLQYGGRAMHRL-----   451
                   *** .** ::*
```

**Figure S2.** Clustal O alignment of the BifB-PCP-SurE di-domain and the SurE (PDB ID 6KSU). Amino acids comprising the PCP domain are coloured yellow, while amino acids comprising the β-lactamase and lipocalin domains are coloured salmon and blue, respectively. Asterisks indicate amino acid conservation.

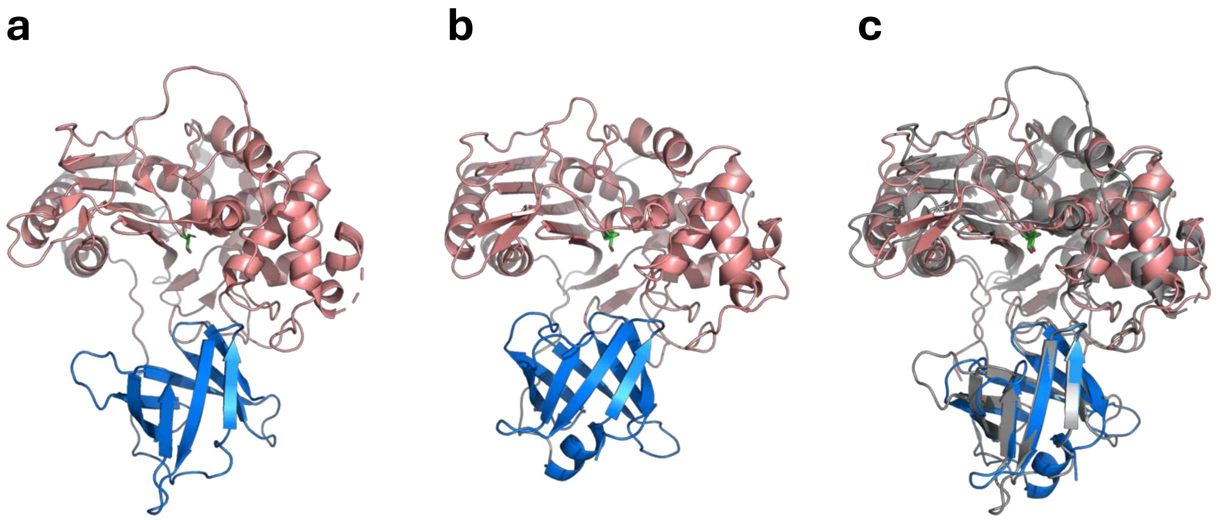


**Figure S3.** Structural analysis of BifB$^{SurE}$. (a) SurE crystal structure (PDB ID 6KSU) (b) AlphaFold3 structural model of BifB$^{SurE}$ domain. For both (a) and (b) salmon colour denotes the β-lactamase domain and blue colour denotes the lipocalin domain and the active site serine residue is coloured green. (c) Superimposition of SurE (grey) and BifB$^{SurE}$ coloured in salmon and blue as in (b). To achieve a more accurate structural comparison, protein chains were broken between the β-lactamase and lipocalin domains prior to alignment in PyMol 2.5.4.

| | Alignment position | 149 ▼ 153 | 289 ▼ 293 |
|---|---|---|---|
| This study | BifB_Eohlys | AHHYV | ELEGH |
| Characterised *in vitro* | FenD_Ethr | IHHLV | SMEGH |
| | TycB_Ephe | IHHLV | NLEGH |
| | TycA_Ephe | IHHLV | NLEGH |
| | GrsA_Ephe | IHHLV | NLEGH |

**Figure S4.** Multisequence alignment of nonribosomal peptide synthetase epimerase (E) domains. Displayed is an excerpt from a Muscle alignment of the E-domain from BifB$^{M3}$ in comparison to characterised E domains. Black triangles indicate proposed active site residues. The E-domain substrate is indicated within the sequence name. Numbers represent position within the alignment. Protein sequences are from[52]; which are available along with the alignment at 10.6084/m9.figshare.27958413.

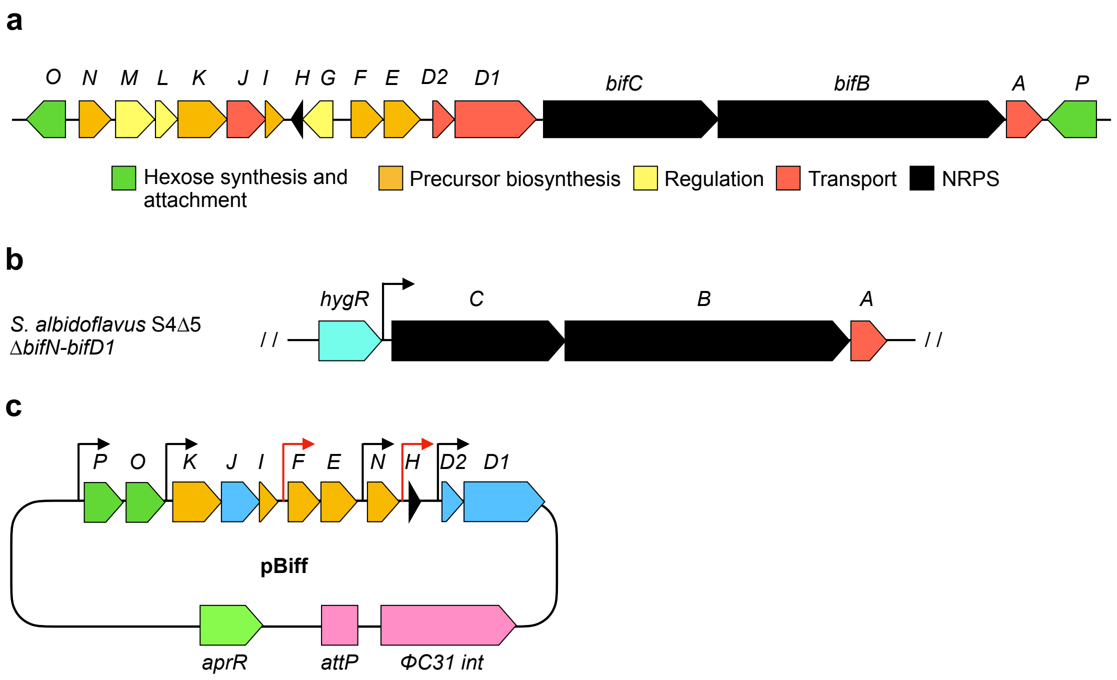


**Figure S5.** De-regulation of the biffamycin *(bif)* biosynthetic gene cluster (BGC). (a) Gene organisation of the *bif* BGC colour-coded by deduced function as indicated. (b) Schematic representation for transcriptional de-regulation of the *bifCBA* in the *S. albidoflavus* S4Δ5 chromosome. *hygR,* resistance gene. (c) Schematic representation of pBiff. *aprR,* apramycin resistance gene; *attP,* phage attachment site; int, integrase; narrow black arrows, *ermE** promoter; narrow red arrows, *rpsL(XC)* promoter.

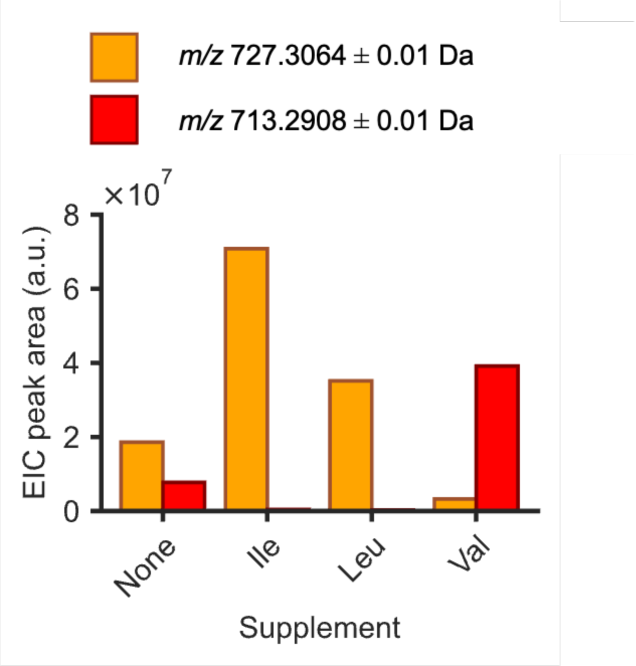


**Figure S6.** Analysis of **1** and **2** production by *S. albidoflavus* S4Δ5Δ*bifO-bifD1*/pBiff when cultivated under amino acid supplementation. Minimal Media supplemented (MM) was supplemented with 5 mM of the indicated amino acid. The histogram was constructed based on the peak area for extracted ion chromatograms (EICs) for **1** (*m/z* 727.2974 ± 0.01 Da) and **2** (*m/z* 713.2908 ± 0.01 Da); a.u., arbitrary units. MS2 fragmentation mapping of 1 and 2 displayed below in Figure S4 and 5 and Supplementary Tables 1 and 2, respectively.

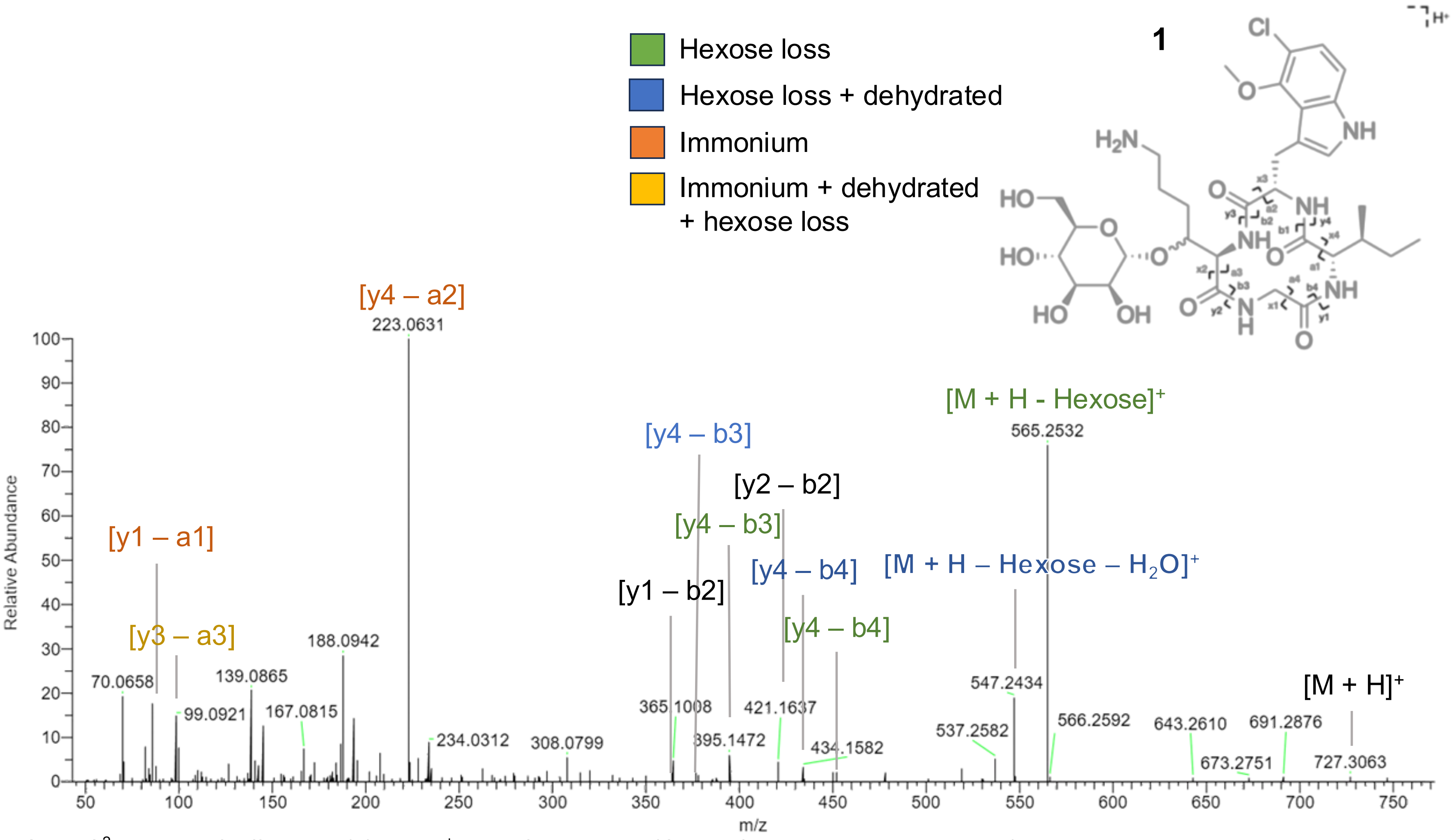


**Figure S7.** $MS^2$ spectra of biffamycin (**1**) $[M+H]^+$ ion *m/z* 727.3051 (Calc *m/z* 727.3064; Δ = -1.8 ppm).

**Table S2.** Mapping of $MS^2$ spectra for biffamycin A (**1**).

| Fragment | | Expected *m/z* | Observed *m/z* | Error (*m/z*) | Error (ppm) |
|---|---|---|---|---|---|
| Biffamycin A (**1**) | $[M+H]^+$ | 727.3064 | 727.3063 | -0.0001 | -0.1 |
| Hexose loss | $[M+H-Hexose]^+$ | 565.2536 | 565.2532 | -0.0004 | -0.7 |
| Hexose loss + dehydration | $[M+H-Hexose - H_2O]^+$ | 547.2430 | 547.2434 | 0.0004 | 0.7 |
| Hexose loss + isoleucine loss | [y4 – b4] | 452.1695 | 452.1684 | -0.0009 | -2.4 |
| Hexose loss + isoleucine loss + dehydration | [y4 – b4] | 434.1590 | 434.1583 | -0.0007 | -1.6 |
| Hexosyl-oxy-lysine /hydroxylysine loss | [y2 – b2] | 421.1637 | 421.1637 | 0.0000 | 0.0 |
| Hexose loss + isoleucine and glycine loss | [y4 – b3] | 395.1481 | 395.1472 | -0.0009 | -2.3 |
| Isoleucine and glycine loss + dehydration | [y4 – b3] | 377.1375 | 377.1360 | -0.0015 | -4.0 |
| Hexosyl-oxy-lysine /hydroxylysine and glycine loss | [y1 – b4] | 364.1422 | 364.1425 | 0.0003 | 0.8 |
| Chloro-methoxy-tryptophan immonium | [y4 – a2] | 223.0633 | 223.0631 | -0.0002 | -0.9 |
| Dehydrated hydroxy-lysine immonium | [y3 – a3] | 99.0917 | 99.0921 | 0.0003 | 4.0 |
| Isoleucine immonium | [y1 – a1] | 86.0964 | 86.0969 | 0.0005 | 5.8 |

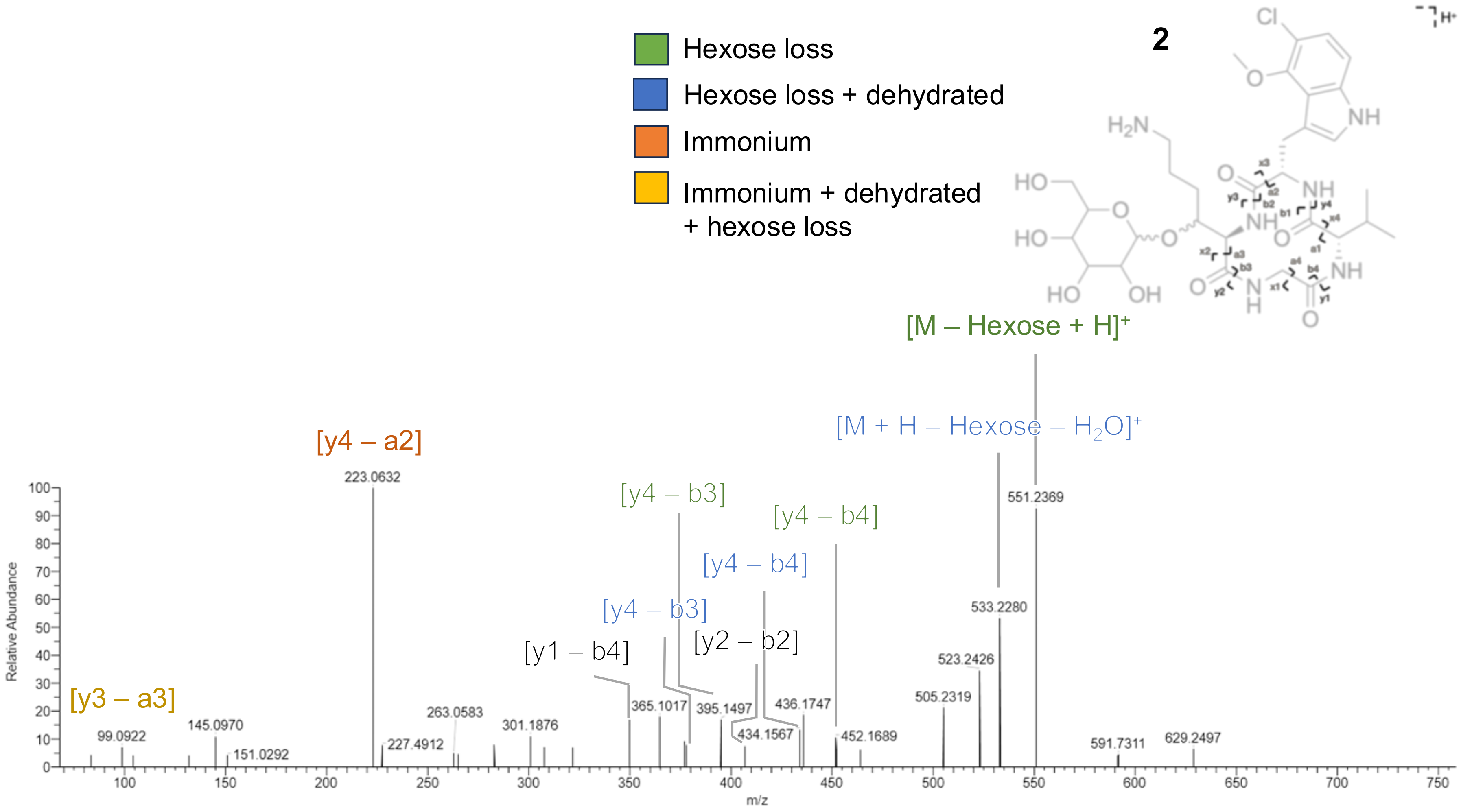


**Figure S8.** MS$^2$ spectra of biffamycin (**2**) $[M+H]^+$ ion *m/z* 713.2905 (Calc *m/z* 713.2908; Δ = -0.4 ppm).

**Table S3.** Mapping of $MS^2$ spectra for biffamycin B (**2**).

| **Fragment** | | **Expected *m/z*** | **Observed *m/z*** | **Error (*m/z*)** | **Error (ppm)** |
|---|---|---|---|---|---|
| Hexose loss | $[M+H\text{-Hexose}]^+$ | 551.2379 | 551.2369 | -0.0010 | -1.8 |
| Hexose loss + dehydration | $[M+H\text{-Hexose -H2O}]^+$ | 533.2274 | 533.2280 | 0.0006 | 1.1 |
| Hexose loss + valine loss | [y4 – b4] | 452.1695 | 452.1695 | 0.0000 | 0 |
| Hexose loss + valine loss + dehydration | [y4 – b4] | 434.1590 | 434.1567 | -0.0023 | -5.3 |
| Hexosyl-oxy-lysine /hydroxylysine loss | [y2 – b2] | 407.1481 | 407.1458 | -0.0023 | -5.6 |
| Hexose loss + valine and glycine loss | [y4 – b3] | 395.1481 | 395.1497 | 0.0016 | 4.0 |
| Hexose loss + valine and glycine loss + dehydration | [y4 – b3] | 377.1375 | 377.1394 | 0.0019 | 5 |
| Hexosyl-oxy-lysine /hydroxylysine and glycine loss | [y1 – b4] | 350.1266 | 350.1285 | 0.0019 | 5.4 |
| Chloro-methoxy-tryptophan immonium | [y4 – a2] | 223.0633 | 223.0632 | -0.0001 | -0.4 |

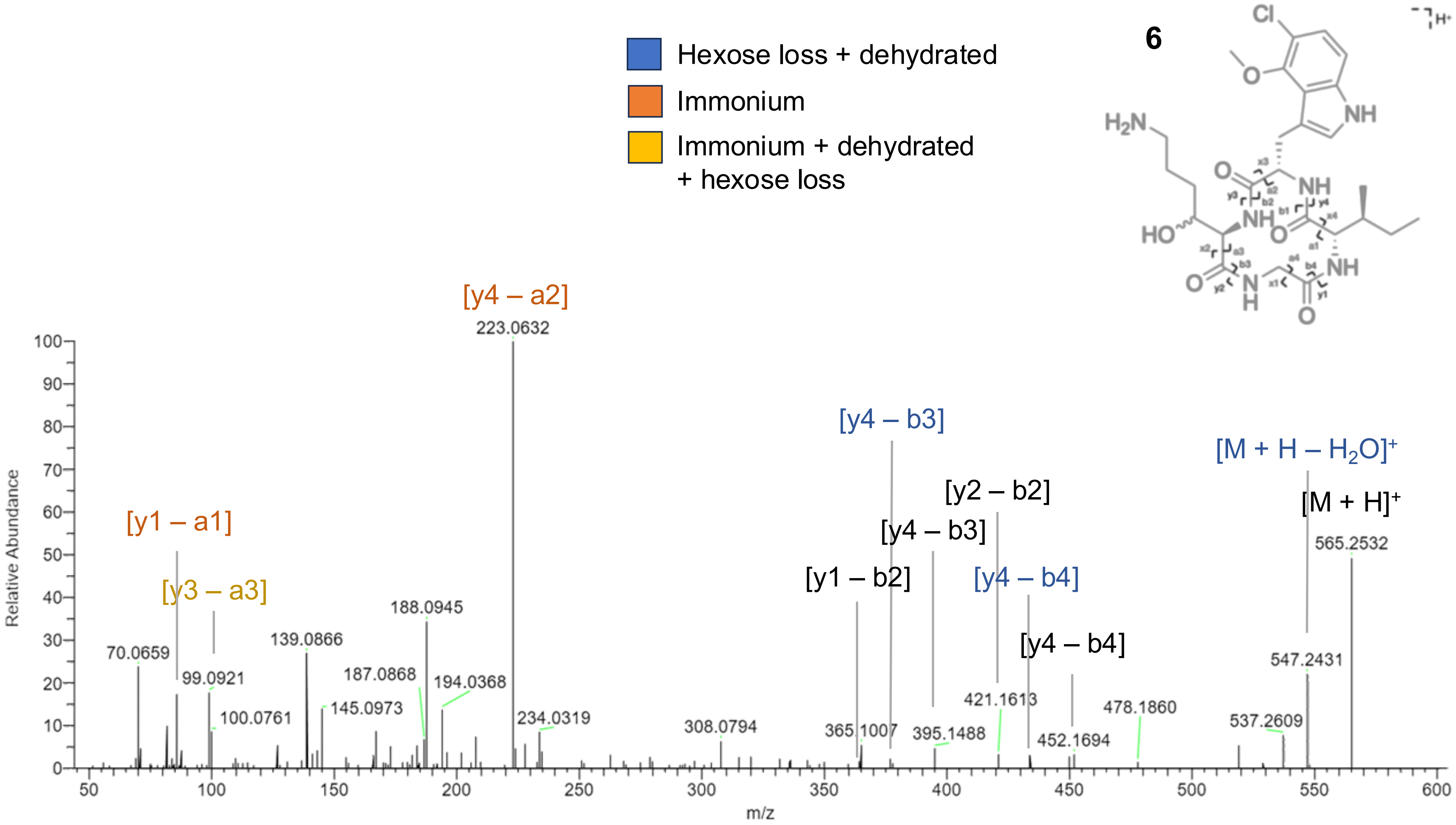


**Figure S9.** MS[2] spectra of biffamycin A aglycone (**6**) [M+H]$^+$ ion *m/z* 565.2535 (Calc *m/z* 565.2536; Δ = -0.2 ppm).

**Table S4.** Mapping of $MS^2$ spectra for biffamycin A aglycone (**6**).

| Fragment | | Expected *m/z* | Observed *m/z* | Error (*m/z*) | Error (ppm) |
|---|---|---|---|---|---|
| Biffamycin A aglycone (**6**) | $[M+H]^+$ | 565.2536 | 565.2532 | -0.0004 | -0.7 |
| Dehydration | $[M+H-H_2O]^+$ | 547.2430 | 547.2431 | 0.0001 | 0.2 |
| Isoleucine loss | [y4 – b4] | 452.1695 | 452.1694 | -0.0001 | -0.2 |
| Isoleucine loss + dehydration | [y4 – b4] | 434.1590 | 434.1596 | 0.0006 | 1.4 |
| Hydroxylysine loss | [y2 – b2] | 421.1637 | 421.1613 | 0.0027 | -5.7 |
| Isoleucine and glycine loss | [y4 – b3] | 395.1481 | 395.1488 | 0.0007 | 1.8 |
| Isoleucine and glycine loss + dehydration | [y4 – b3] | 377.1375 | 377.1367 | -0.0008 | -2.1 |
| Hydroxylysine and glycine loss | [y1 – b2] | 364.1422 | 364.1413 | -0.0009 | -2.5 |
| Chloro-methoxy-tryptophan immonium | [y4 – b2] | 223.0633 | 223.0632 | -0.0001 | -0.4 |
| Dehydrated hydroxy-lysine immonium | [y3 – b3] | 99.0917 | 99.0921 | 0.0003 | 4.0 |
| Isoleucine immonium | [y1 – b1] | 86.0964 | 86.0970 | 0.0006 | 7.0 |

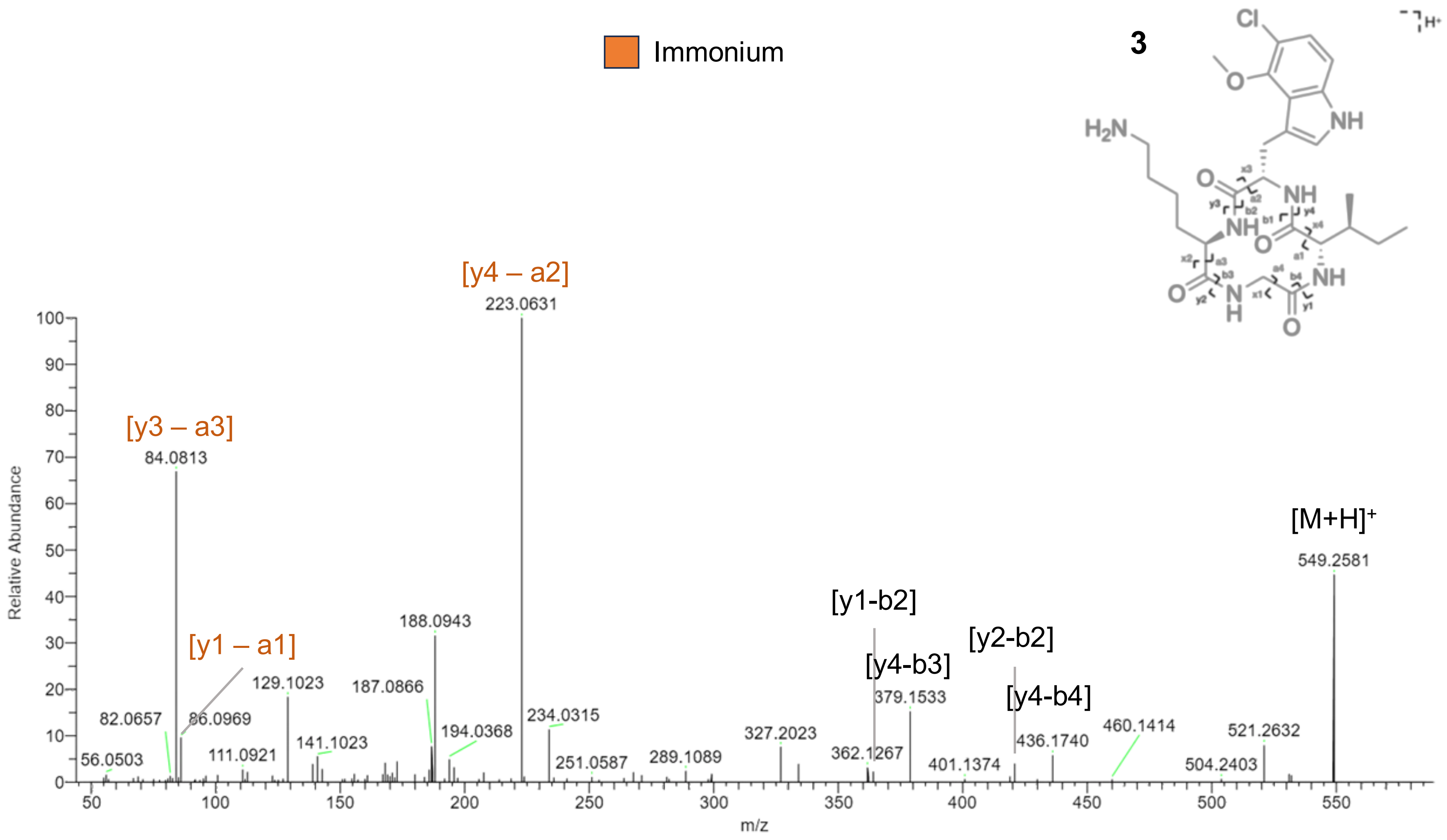


**Figure S10.** $MS^2$ spectra of *des*-hydroxy biffamycin A aglycone (**3**) $[M+H]^+$ ion *m/z* 549.2585 (Calc *m/z* 549.2587; Δ = -0.4 ppm).

**Table S5.** Mapping of MS$^2$ spectra for biffamycin *des*-hydroxy-aglycone (**3**).

| Fragment | | Expected *m/z* | Observed *m/z* | Error (*m/z*) | Error (ppm) |
|---|---|---|---|---|---|
| *Des*-hydroxy-aglycone (**3**) | $[M+H]^+$ | 549.2587 | 549.2581 | -0.0006 | -1.1 |
| Isoleucine loss | [y4 – b4] | 436.1746 | 436.1740 | -0.0006 | -1.4 |
| Lysine loss | [y2 – b2] | 421.1637 | 421.1626 | -0.0011 | -2.6 |
| Isoleucine and glycine loss | [y4 – b3] | 379.1531 | 379.1533 | 0.0002 | 0.5 |
| Lysine and glycine loss | [y1 – b2] | 364.1422 | 364.1401 | -0.0019 | -5.8 |
| Chloro-methoxy-tryptophan immonium | [y4 – a2] | 223.0633 | 223.0631 | -0.0002 | -0.9 |
| Isoleucine immonium | [y1 – a1] | 86.0964 | 86.0969 | 0.0005 | 5.8 |
| Lysine derived immonium | [y3 – a3] | 84.0808 | 84.0813 | 0.0005 | 5.9 |

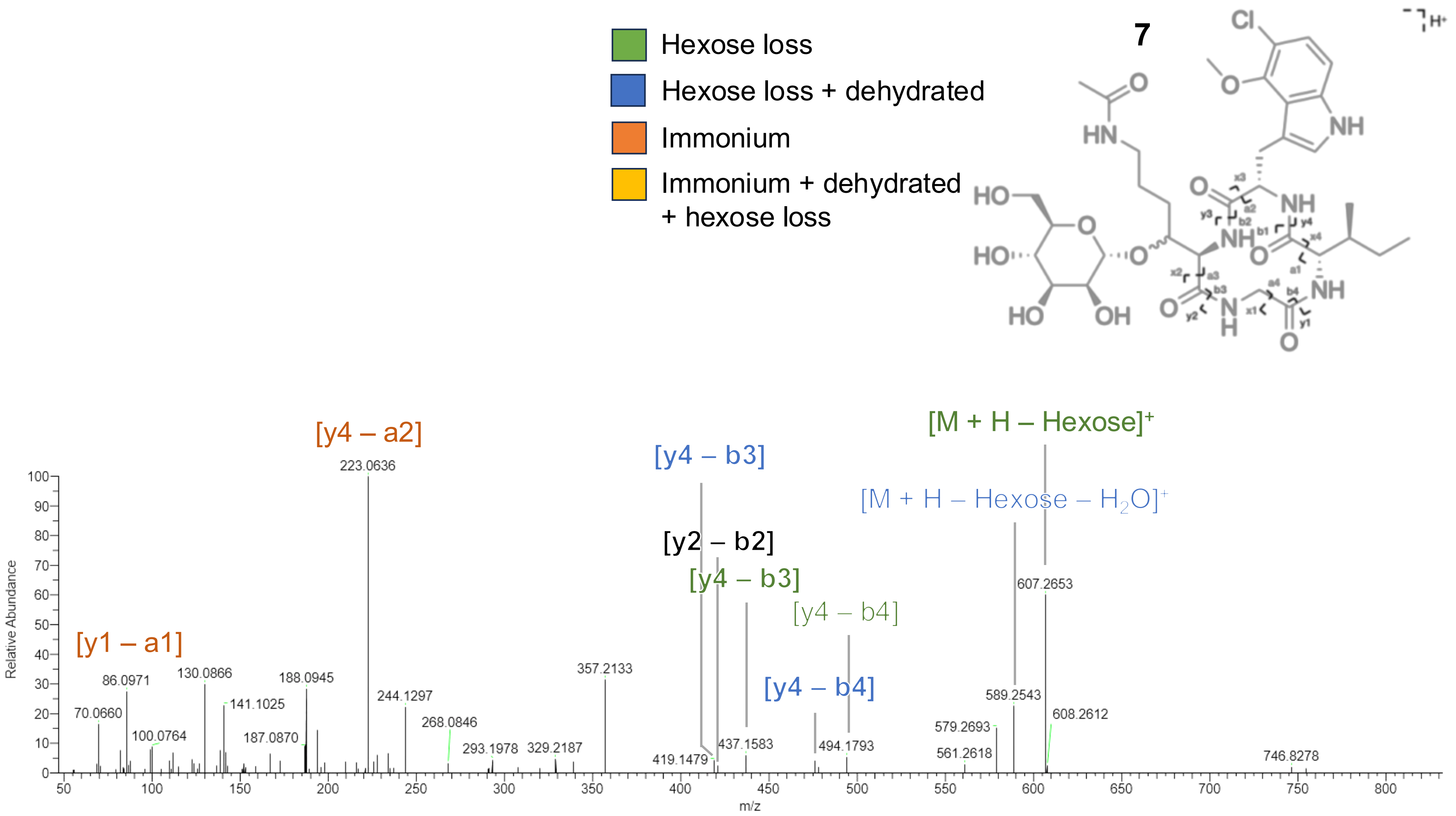


**Figure S11.** $MS^2$ spectra of *N*-acetyl biffamycin A (**7**) $[M+NH_4]^+$ ion *m/z* 786.3433 (Calc *m/z* 786.34353; Δ = -0.3 ppm).

**Table S6.** Mapping of $MS^2$ spectra for *N*-acetyl biffamycin A (**7**).

| Fragment | | Expected *m/z* | Observed *m/z* | Error (*m/z*) | Error (ppm) |
|---|---|---|---|---|---|
| Hexose loss | $[M+H\text{-}Hexose]^+$ | 607.2642 | 607.2653 | 0.0011 | 1.8 |
| Hexose loss + dehydration | $[M+H\text{-}Hexose\text{-}H_2O]^+$ | 589.2536 | 589.2543 | 0.0007 | 1.2 |
| Hexose loss + isoleucine loss | [y4 – b4] | 494.1801 | 494.1793 | 0.0008 | -1.6 |
| Hexose loss + isoleucine loss + dehydration | [y4 – b4] | 476.1696 | 476.1685 | -0.0011 | -2.3 |
| Hexose loss + *N*-acetyl-hydroxylysine loss | [y2 – b2] | 421.1637 | 421.1647 | 0.0010 | 2.4 |
| Hexose loss + isoleucine and glycine loss | [y4 – b3] | 437.1587 | 437.1583 | -0.0004 | -0.9 |
| Isoleucine and glycine loss + dehydration | [y4 – b3] | 419.1481 | 419.1479 | -0.0002 | -0.5 |
| Chloro-methoxy-tryptophan immonium | [y4 – a2] | 223.0633 | 223.0636 | 0.0003 | 1.3 |
| Isoleucine immonium | [y1 – a1] | 86.0964 | 86.0971 | 0.0007 | 8.1 |

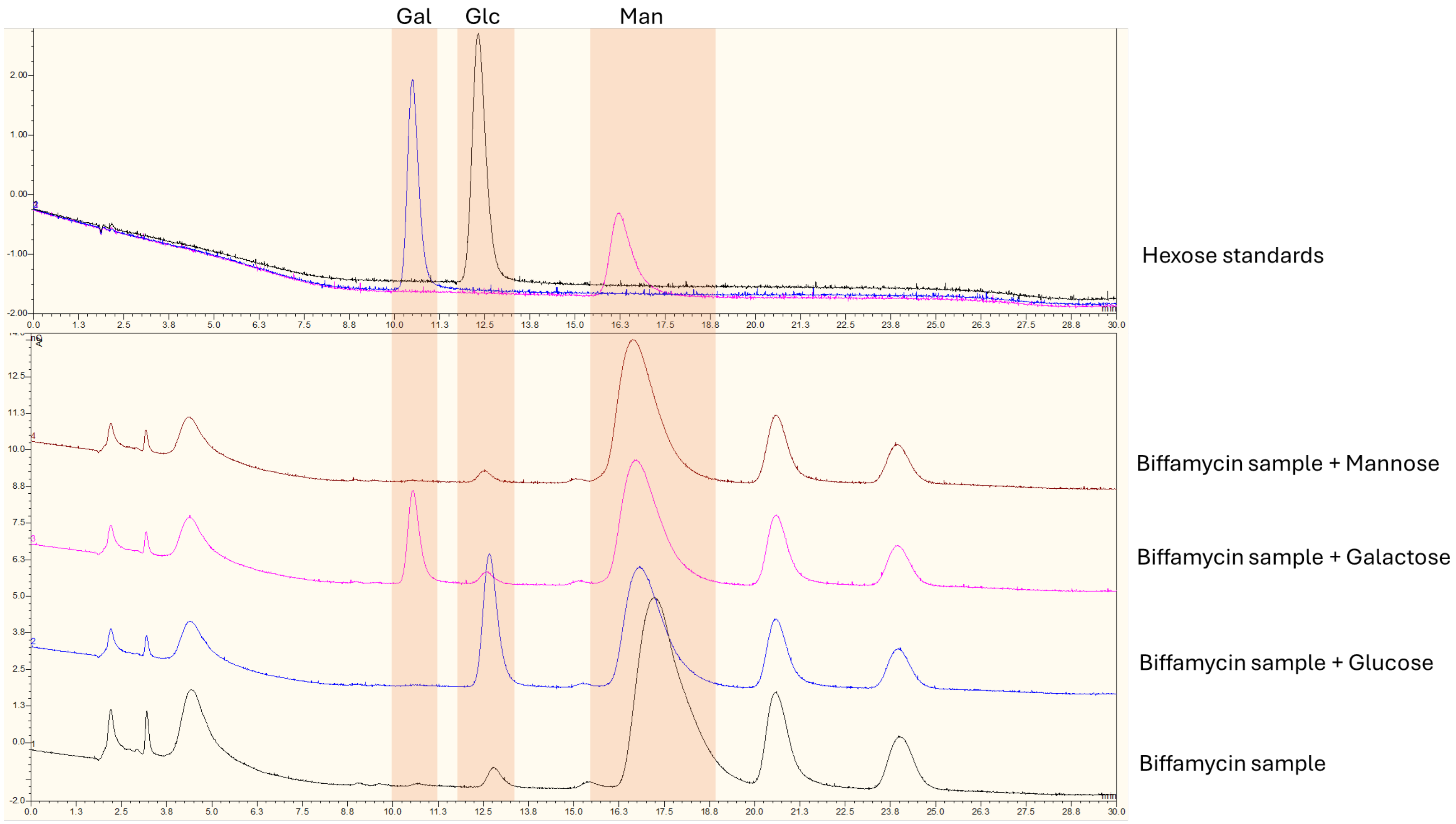


**Figure S12.** Carbohydrate analysis of products of acid hydrolysis of **1** analysed by high performance anion exchange chromatography with pulsed amperometric detection (HPAEC-PAD). Gal, galactose; Glc, glucose; Man, mannose.

### 1D and 2D NMR Spectra and $^{1}H$ and $^{13}C$ NMR spectra assignment tables for compounds 3-7

**Table S7.** Resonance assignments in the $^{1}H$ and $^{13}C$ NMR spectra.

| | Compound **3** | | Compound **4** | | Compound **5** | | Compound **6** | | Compound **7** | |
|---|---|---|---|---|---|---|---|---|---|---|
| No | $\delta_H$ (Multiplicity, *J* ) | $\delta_C$ | $\delta_H$ (Multiplicity, *J* ) | $\delta_C$ | $\delta_H$ (Multiplicity, *J* ) | $\delta_C$ | $\delta_H$ (Multiplicity, *J* ) | $\delta_C$ | $\delta_H$ (Multiplicity, *J* ) | $\delta_C$ |
| G1 | - | 170.6 | | | | | - | 170.7 | - | 170.4 |
| G2' | 3.80 -3.70 (m) | 44.5 | | | | | 3.59 (dd, 14.1,6.5 Hz) | 44.7 | 3.61*[a] | 44.8 |
| G2'' | 3.44 (br s) | 44.5 | | | | | 3.54 (m) | 44.7 | 3.52*[b] | 44.8 |
| GNH | 8.45 (m) | - | | | | | 8.64 (br s) | - | 8.03 (s) | - |
| I1 | - | 173.3 | | | | | - | 173.3 | - | 173 |
| I2 | 3.80 -3.70 (m) | 61.4 | | | | | 3.74 (m) | 61.5 | 3.74 (t, 10.6 Hz) | 61.5 |
| I3 | 1.79 (m) | 34.7 | | | | | 1.79 (m) | 34.7 | 1.66 | 34.7 |
| I4' | 1.46 - 1.38 (m) | 25.5 | | | | | 1.48 - 1.39 (m) | 25.4 | 1.42 (m) | 25.3 |
| I4'' | 1.00 (dt, 14.2,7.6 Hz) | 25.5 | | | | | 0.98 (m) | 25.4 | 0.97 (m) | 25.3 |
| I5 | 0.79 (t, 7.4 Hz) | 10.7 | | | | | 0.78 (t, 7.4 Hz) | 10.7 | 0.79 (t, 7.4 Hz) | 10.7 |
| I6 | 0.61 (d, 6.7 Hz) | 15.6 | | | | | 0.57 (d, 6.6 Hz) | 15.5 | 0.52 (d, 6.7 Hz) | 15.5 |
| INH | 8.58 (m) | - | | | | | 8.49 (m) | - | 7.53 | - |
| K1 | - | 172.5 | | | | | - | 172.7 | - | 171.5 |
| K2 | 4.19 (m) | 53.1 | | | | | 4.14 (t, 7.6 Hz) | 58.3 | 4.47 (m) | 56.8 |
| K3 | 1.63 (m); 1.50 (m) | 29.4 | | | | | 3.75 (m) | 68.9 | 3.83 (m) | 75.2 |
| K4 | 1.19 (s) | 23.1 | | | | | 1.38 - 1.23 (m) | 31.3 | 1.49 (m) | 27.5 |
| K5 | 1.46 - 1.38 (m) | 25.5 | | | | | 1.48 -1.39 (m) | 25.4 | 1.42 (m) | 25 |
| K6 | 2.6 (m) | 40.3 | | | | | 2.61 (s) | 40.9 | 2.98 (q, 6.6 Hz) | 39.2 |
| KNH | 7.57 (br s) | - | | | | | 7.17 (d, 8.5 Hz) | - | 7.53 | - |
| W1 | - | 172.6 | - | 170.6 | - | 174.6 | - | 172.7 | - | 172.8 |
| W2 | 4.60 (q, 8.2 Hz) | 53.8 | 3.59 (dd, 10.1,3.8 Hz) | 55.6 | 4.48 (ddd, 9.8,8.0,4.5 Hz) | 53.9 | 4.63 (m) | 53.7 | 4.65 (q, 8.3 Hz) | 53.7 |
| W3' | 3.23 (dd, 15.1,7.1 Hz) | 26.7 | 2.93 (ddd, 15.5,10.0,2.5 Hz) | 28.8 | 3.32 (dd, 14.8,4.5 Hz) | 28.7 | 3.21 (m) | 26.4 | 3.15 (m) | 26.4 |
| W3'' | 3.03 (dd, 15.0,8.3 Hz) | 26.7 | 3.49 (dd, 15.3,3.7 Hz) | 28.8 | 2.99 (dd, 14.8,9.7 Hz) | 28.7 | 3.07 (dd, 15.0,8.9 Hz) | 26.4 | 3.15 (m) | 26.4 |
| W4 | - | 110.4 | - | 110.0 | - | 110.6 | - | 110.5 | - | 110.4 |
| W5 | 7.08 (s) | 124.8 | 7.21 (s) | 125.9 | 7.14 (d, 2.4 Hz) | 125.3 | 7.09 (br s) | 125.0 | 7.08 (s) | 124.6 |
| W6 | 11.17 (s) | - | 11.21 (m) | - | 11.09 (br s, Hz) | - | 11.14 (br s) | - | 10.97 | - |
| W7 | - | 137.5 | - | 138.0 | - | 137.5 | - | 137.5 | - | 137.4 |
| W8 | 7.10 (d, 8.6 Hz) | 109.3 | 7.15 (d, 8.5 Hz) | 109.4 | 7.12 (d, 8.6 Hz) | 109.4 | 7.11 (d, 8.7 Hz) | 109.3 | 7.11 (d, 8.6 Hz) | 109.4 |
| W9 | 7.04 (d, 8.6 Hz) | 122.7 | 7.07 (d, 8.5 Hz) | 122.8 | 7.04 (d, 8.5 Hz) | 122.7 | 7.04 (d, 8.6 Hz) | 122.7 | 7.04 (d, 8.6 Hz) | 122.6 |
| W10 | - | 115.7 | - | 115.8 | - | 115.7 | - | 115.7 | - | 115.7 |
| W11 | - | 149.3 | - | 149.3 | - | 149.3 | - | 149.3 | - | 149.3 |
| W12 | - | 122 | - | 122.0 | - | 121.9 | - | 122.0 | - | 122.2 |
| W13 | 3.86 (s) | 61.8 | 3.88 (s) | 62.0 | 3.86 (s) | 61.8 | 3.86 (s) | 61.8 | 3.87 (s) | 61.4 |
| WNH | 8.26 (m) | - | | | 8.03 (d, 8.0 Hz) | - | 8.38 (d, 9.2 Hz) | - | 7.60 (d, 9.0 Hz) | - |
| W14 | | | | | - | 169.4 | | | | |
| W15 | | | | | 1.77 (s) | 23.0 | | | | |
| K6NH | | | | | | | | | 7.85 | - |
| 1 | | | | | | | | | - | 169.4 |
| 2 | | | | | | | | | 1.78 (s) | 23.1 |
| M1 | | | | | | | | | 4.70 (s) | 99.3 |
| M2 | | | | | | | | | 3.57 (s) | 71.1 |
| M3 | | | | | | | | | 3.40*[c] | 71.1 |
| M4 | | | | | | | | | 3.43*[c] | 67.2 |
| M5 | | | | | | | | | 3.43*[c] | 74.6 |
| M6' | | | | | | | | | 3.64*[a] | 61.8 |
| M6'' | | | | | | | | | 3.49*[b] | 61.8 |

*$\delta_H$ was defined from 2D HSQCed. The signal was a part of multiplet: [a]3.68-3.59; b [b]3.54-3.46; [c]3.45-37.

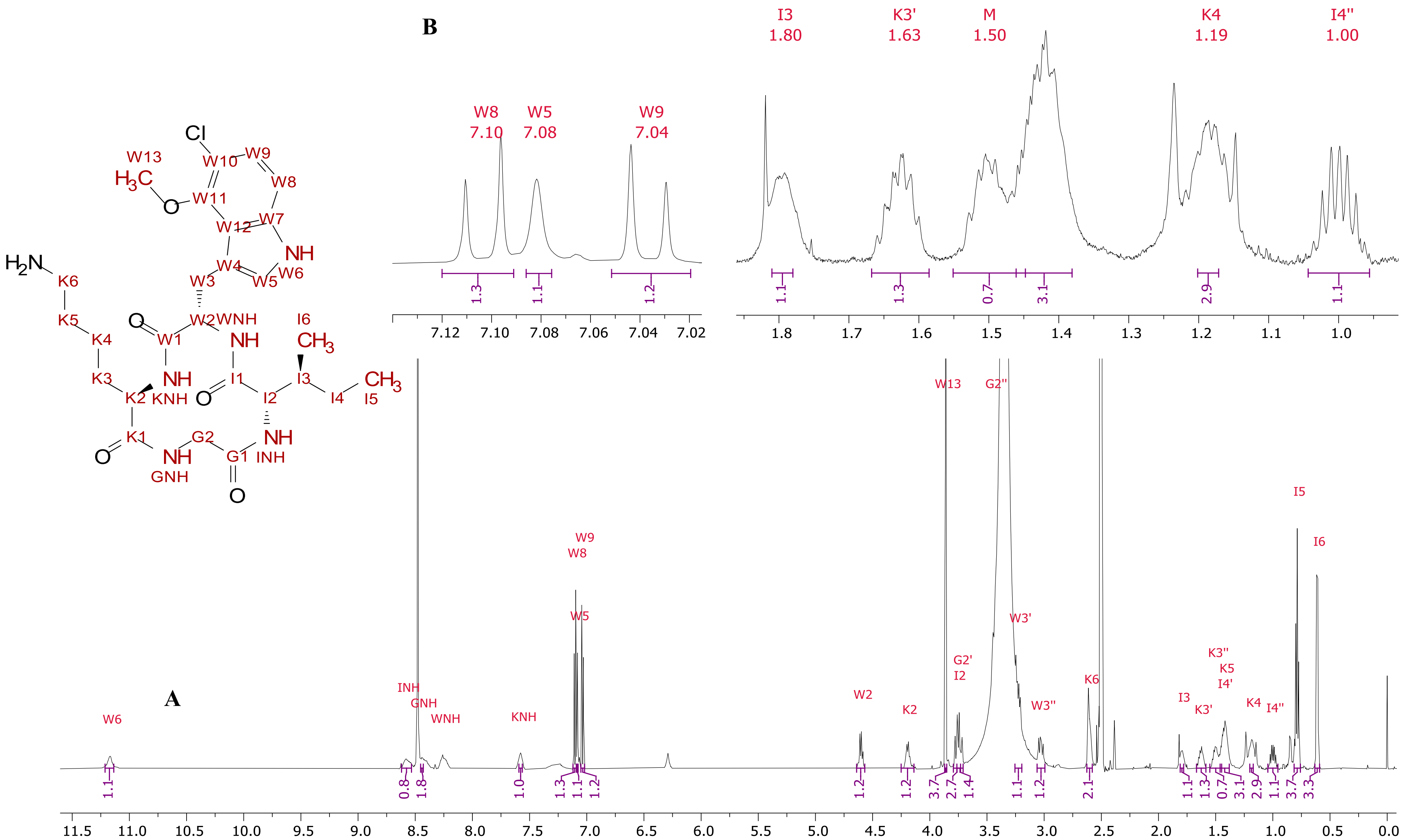


**Figure S13.** [1]H NMR of **3** (DMSO-d6, 600 MHz, 298 K). (a) Whole spectrum overview and (b) an expansion of assigned signals. Multiplets of H-G2" and H-W3' are hidden under the overlapping signal of water.

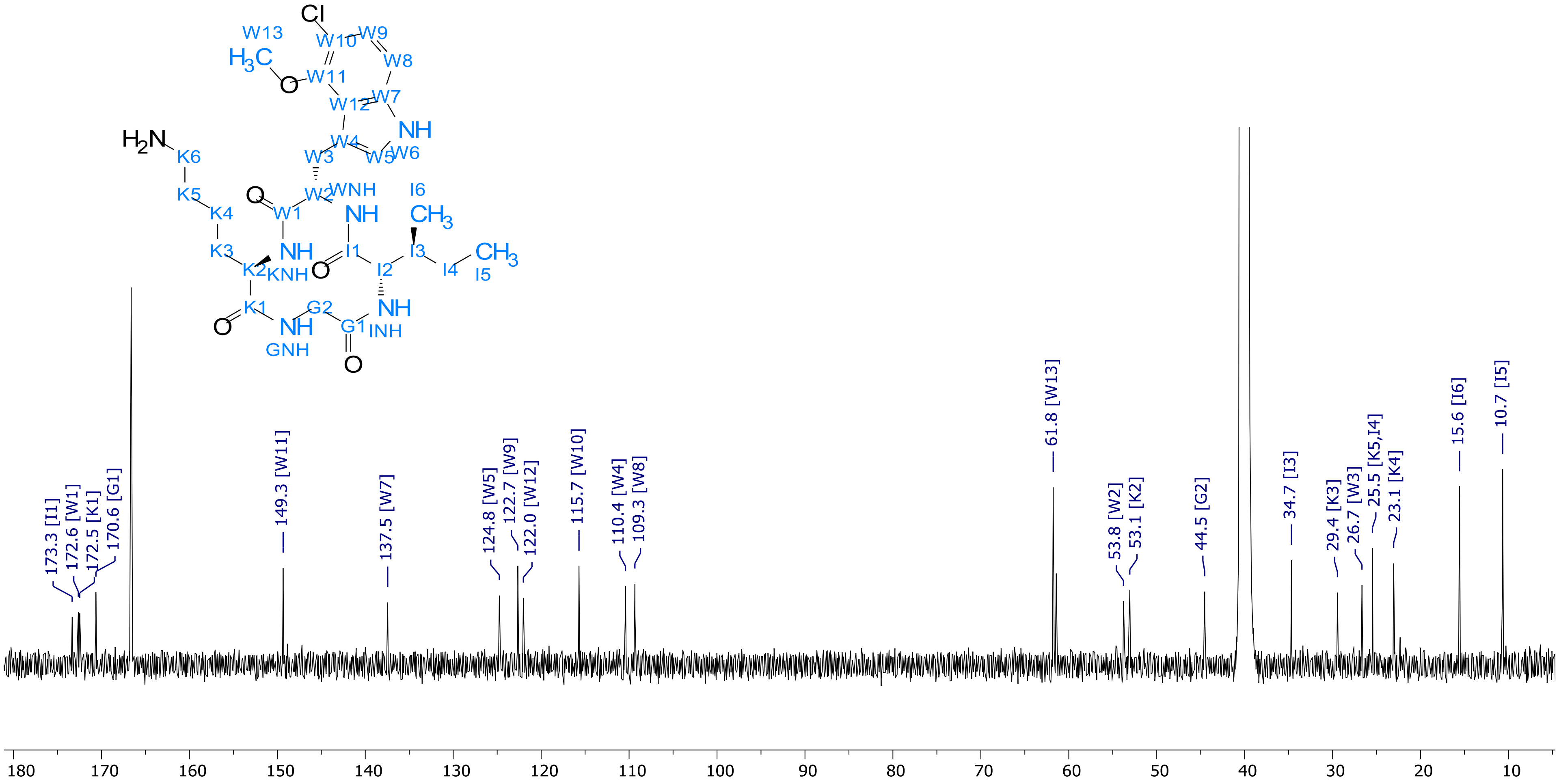


**Figure S14.** $^{13}$C NMR of compound **3** (DMSO-$d_6$, 151 MHz, 298 K). Unambiguous assignment of K5 was complicated since HSQC and HMBC data analysis for corresponding peak was inconclusive. Resonance of C-K5 was assigned to 25.5 ppm coinciding with resonance of I4. An alternative assignment of K5 overlapping with signal of K3 at 29.4 ppm is also possible.

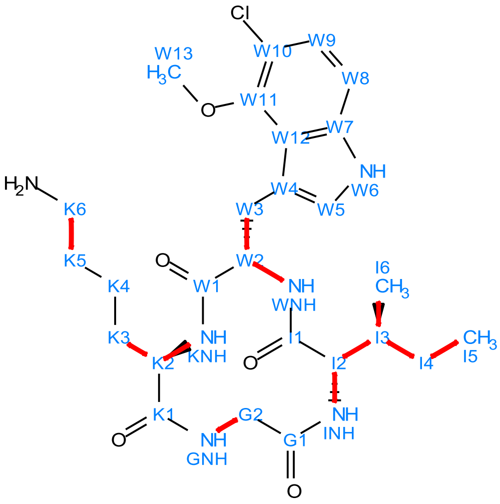

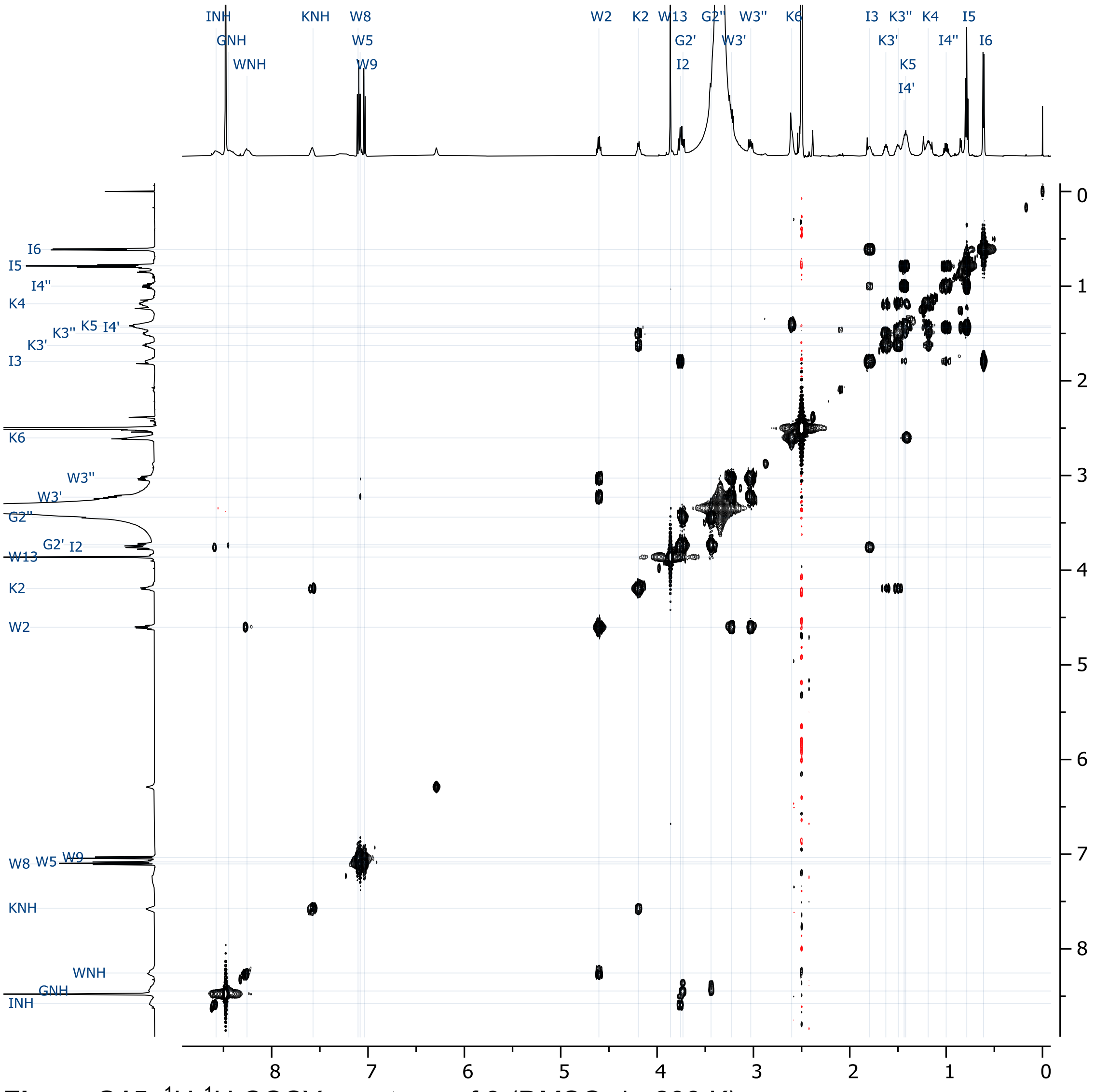

**Figure S15.** $^{1}H$-$^{1}H$ COSY spectrum of **3** (DMSO-$d_6$, 298 K).

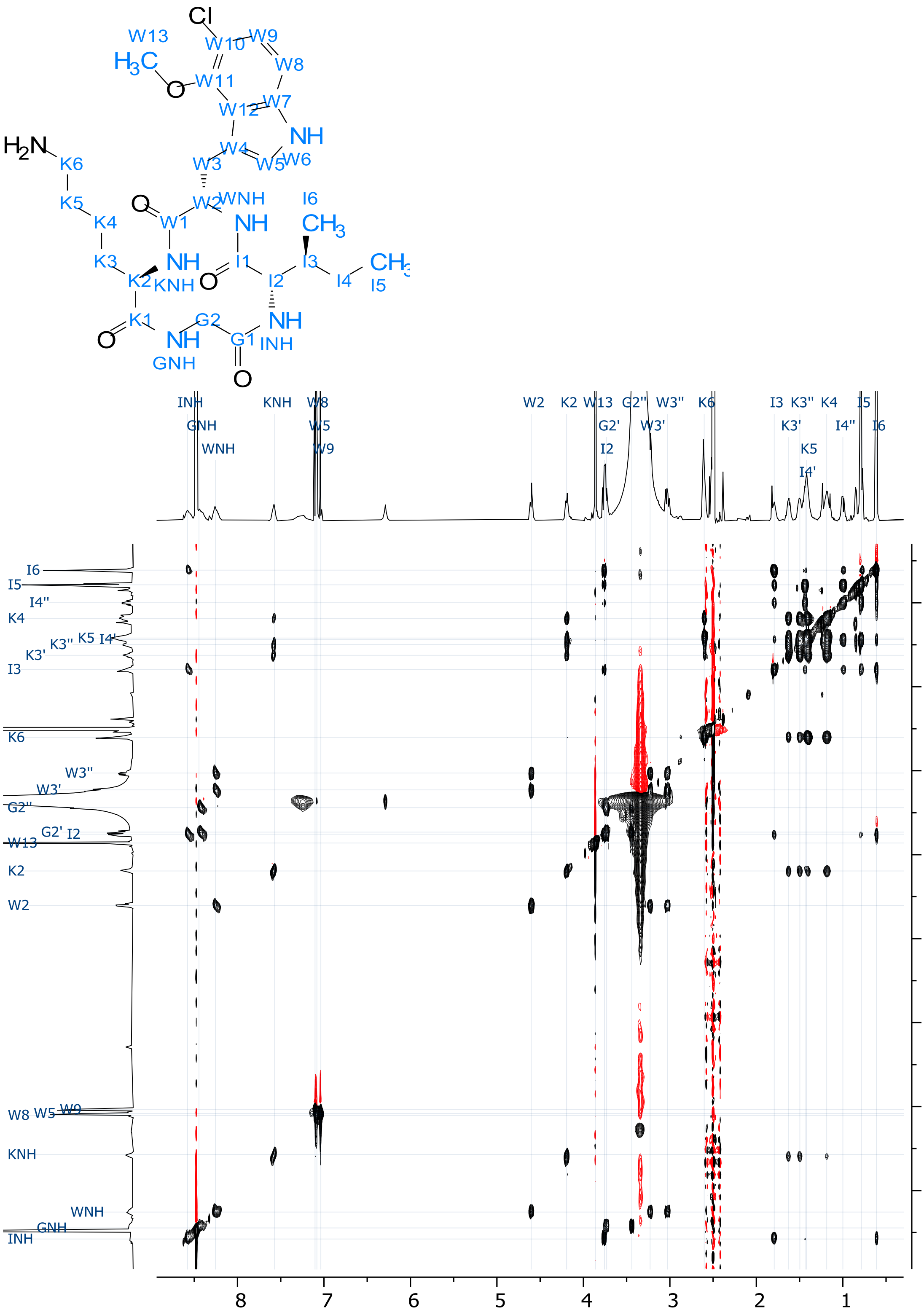

**Figure S16.** 2D TOCSY spectrum of **3** (DMSO-d$_6$, 298 K).

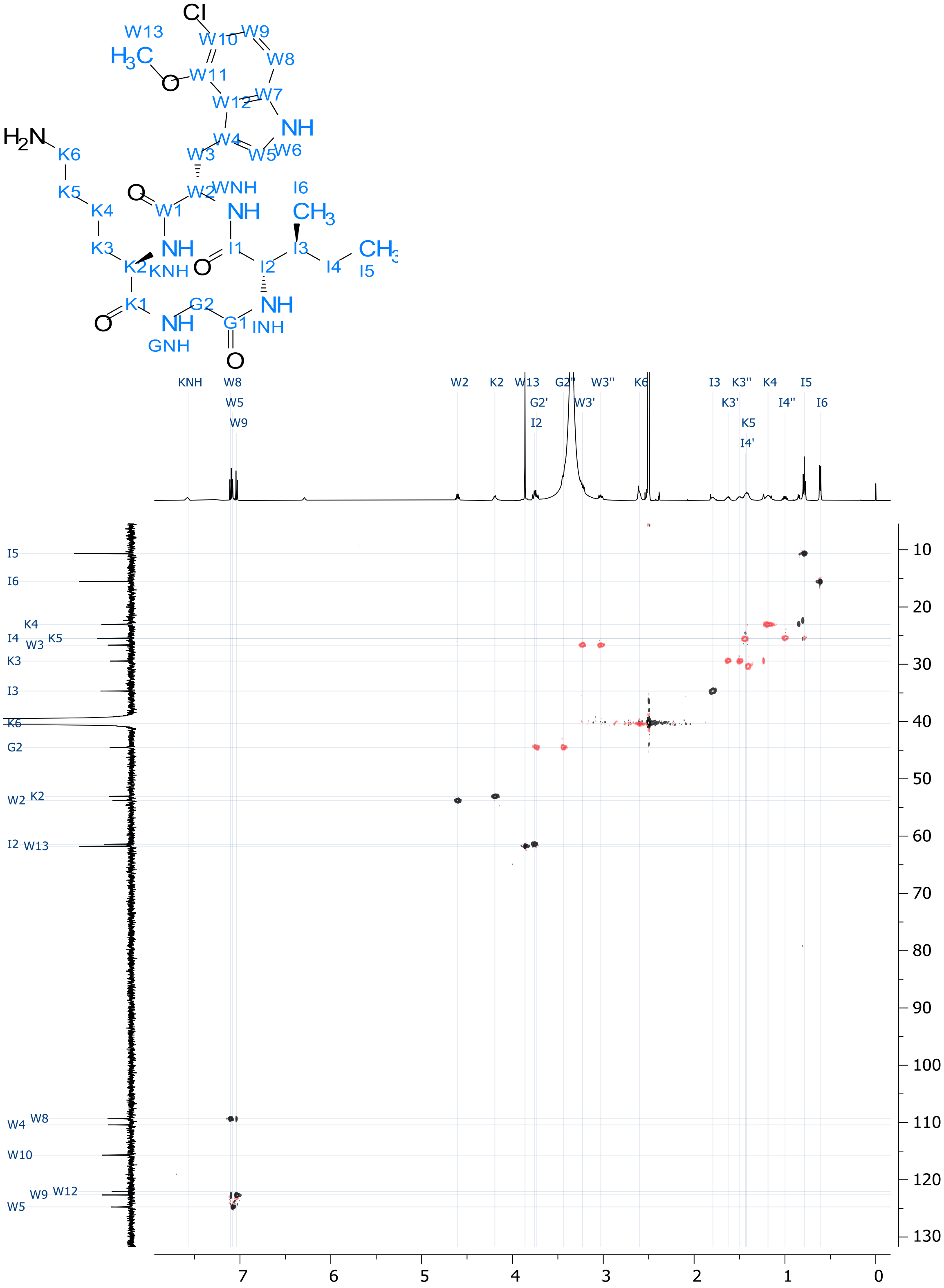


**Figure S17.** $^{1}$H-$^{13}$C HSQC-edited spectrum of **3** (DMSO-d$_6$, 298 K).

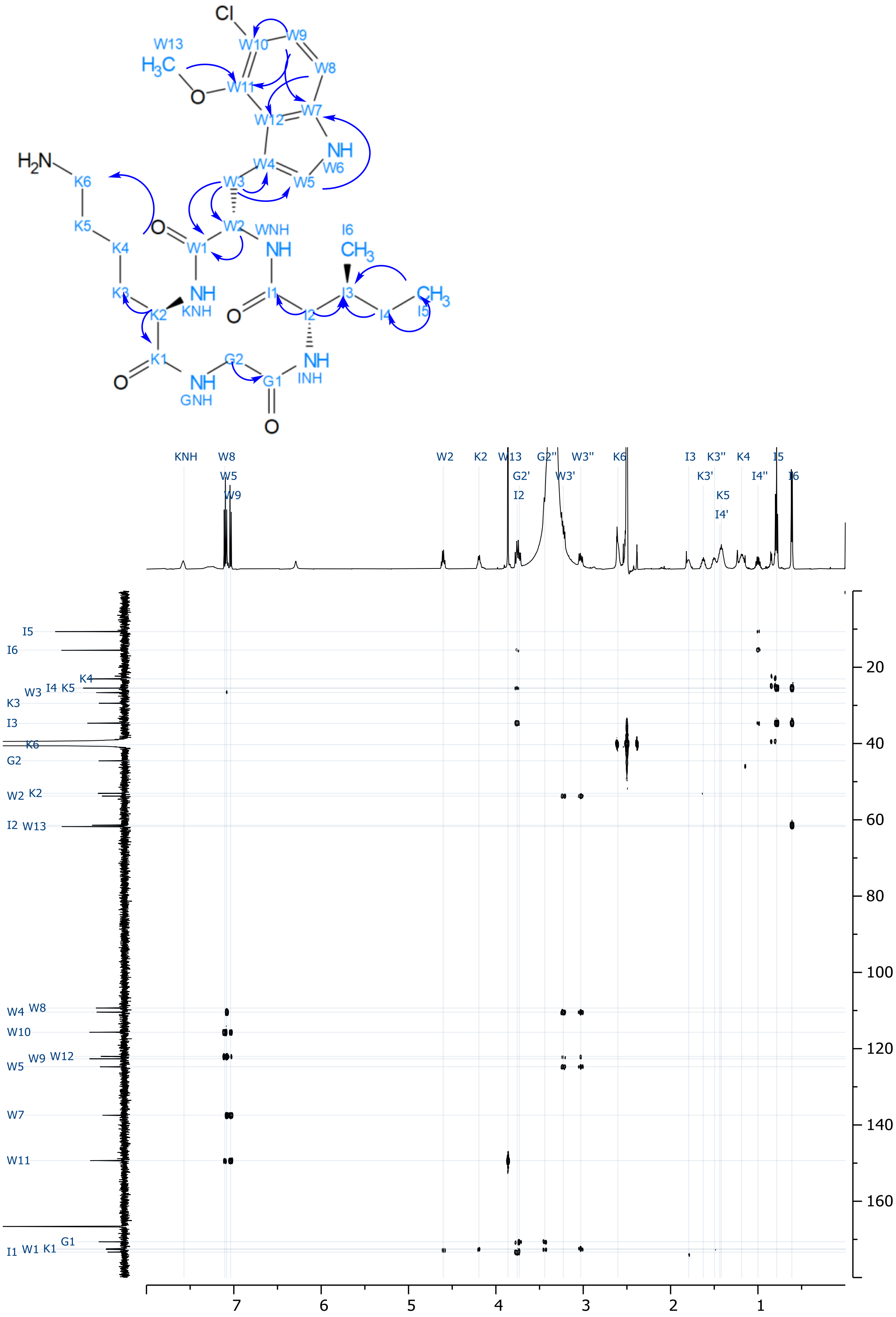


**Figure S18.** $^{1}H$-$^{13}C$ HMBC spectrum of **3** (DMSO-d6, 298 K).

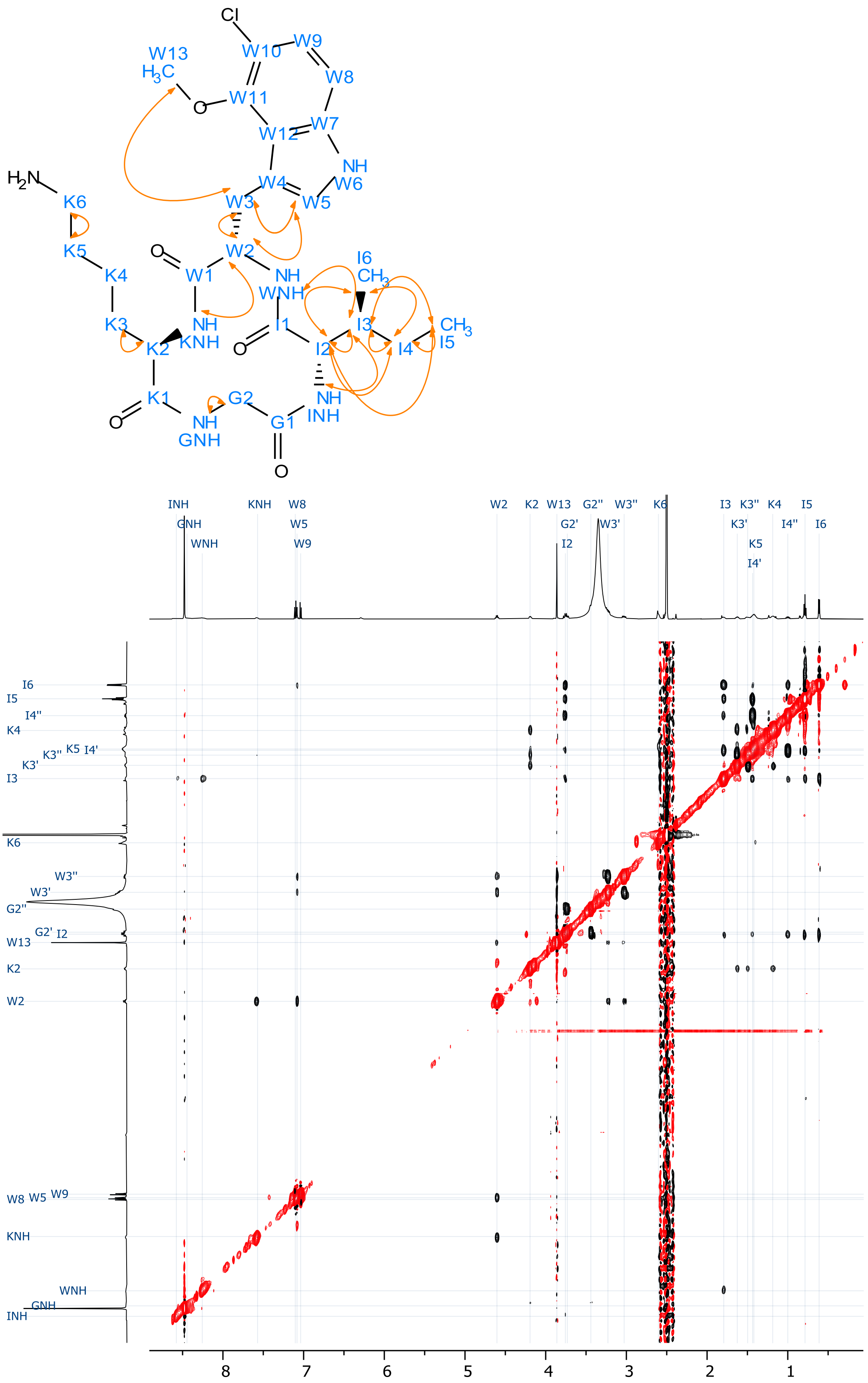


**Figure S19.** $^1H$-$^1H$ ROESY spectrum of **3** (DMSO-$d_6$, 298 K).

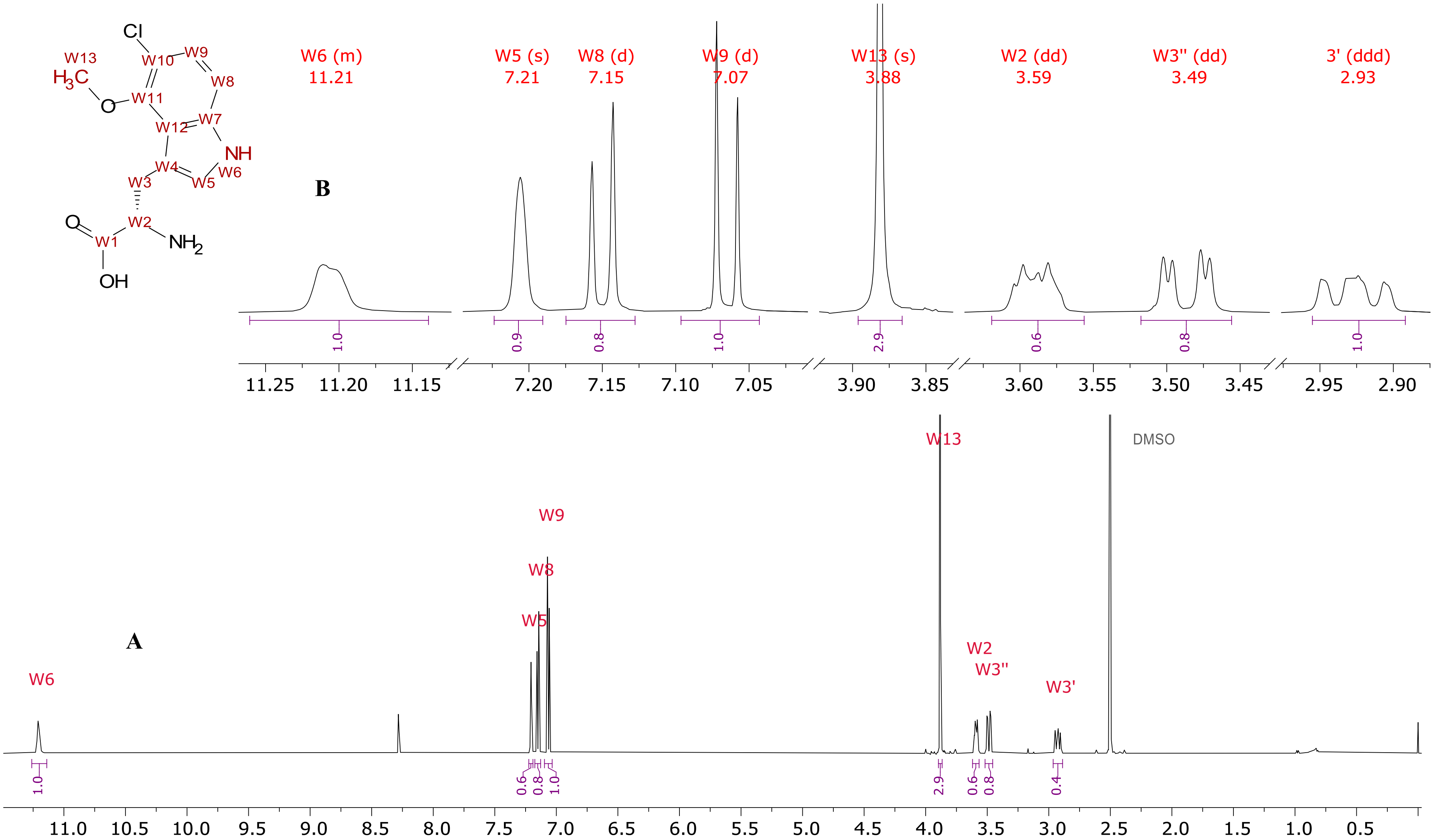


**Figure S20.** $^{1}$H NMR spectrum of **4** (DMSO-d$_6$, 600 MHz, 298 K). (a) Whole spectrum overview and (b) an expansion of assigned signals.

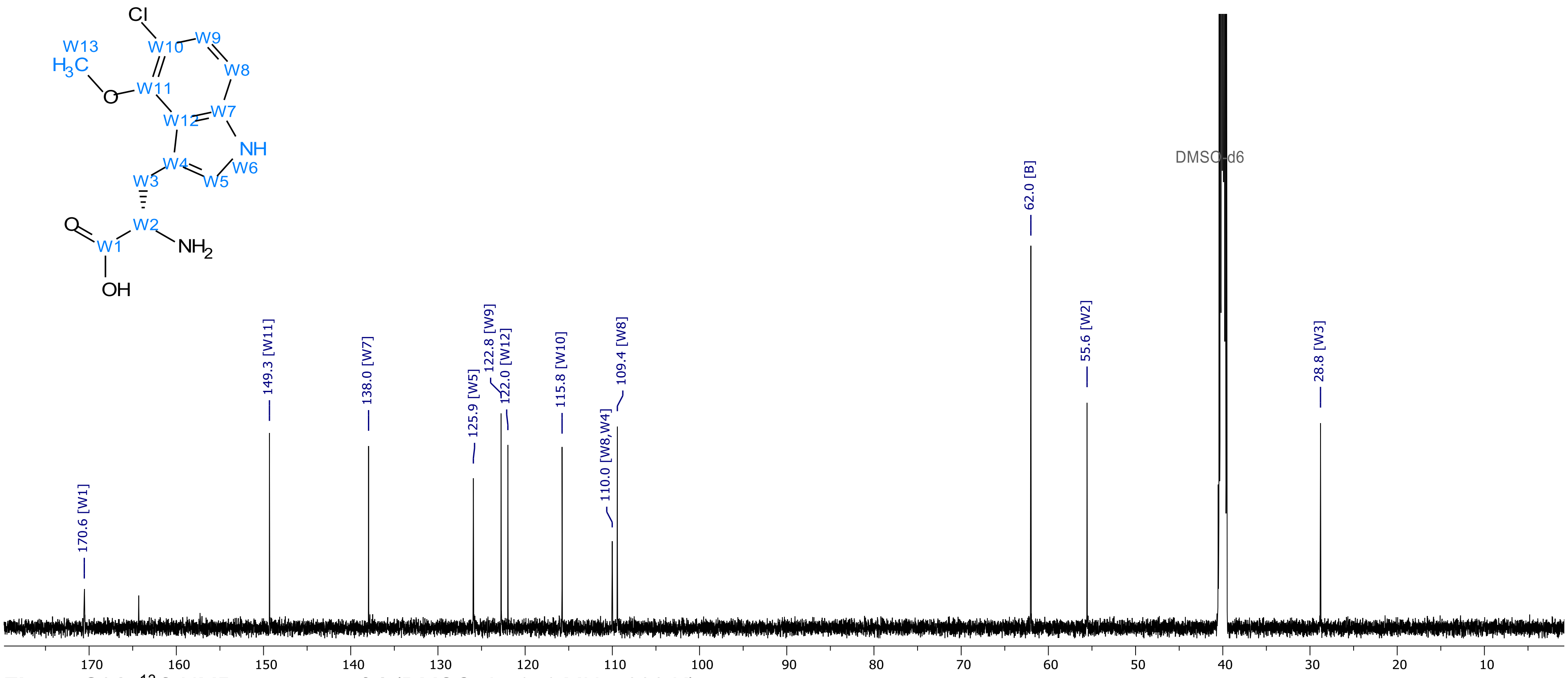


**Figure S21.** $^{13}$C NMR spectrum of **4** (DMSO-$d_6$, 151 MHz, 298 K).

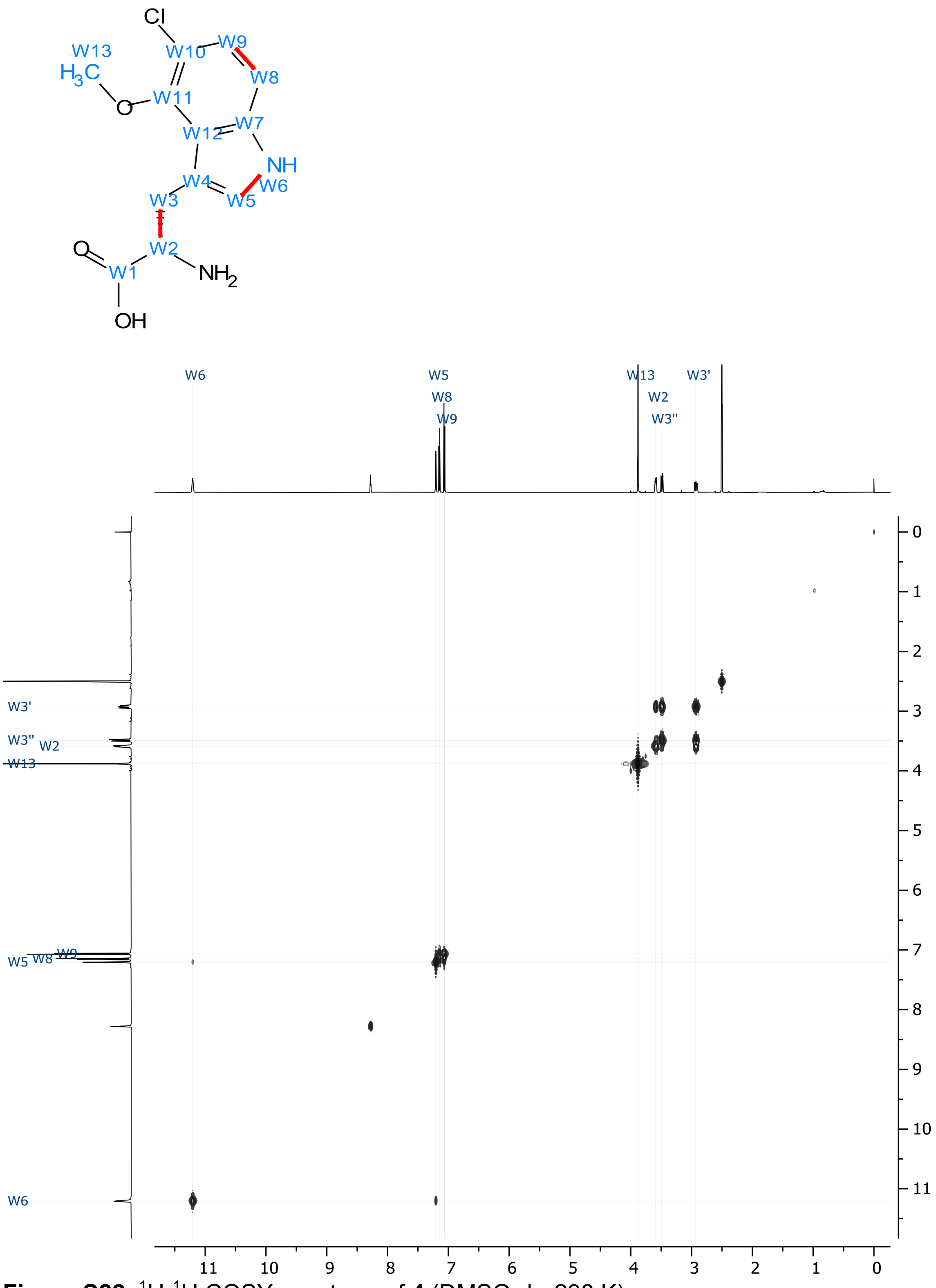


**Figure S22.** $^1$H-$^1$H COSY spectrum of **4** (DMSO-d$_6$, 298 K).

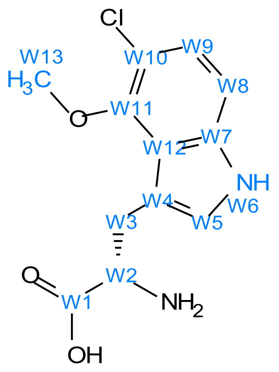


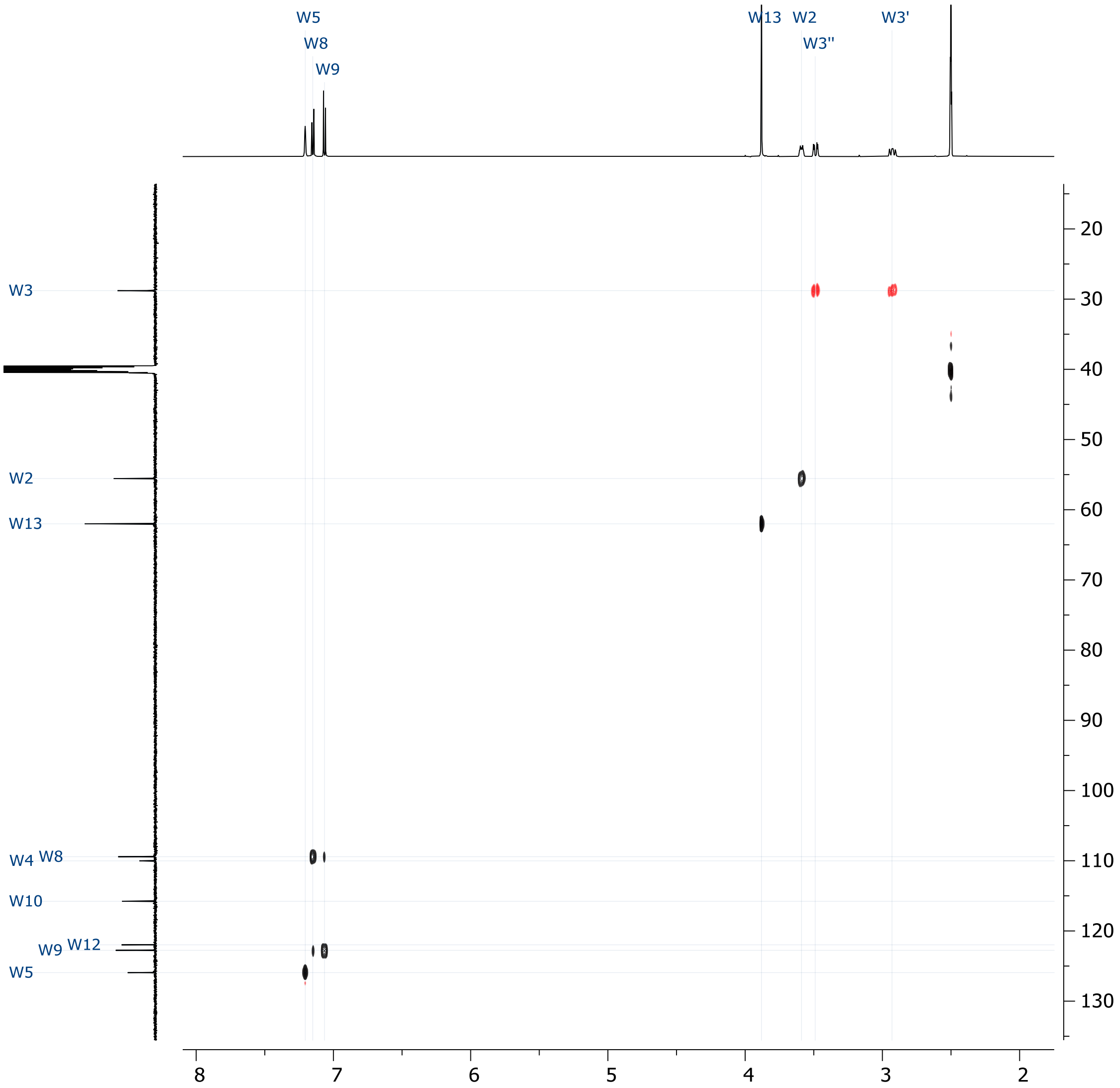


**Figure S23.** $^{1}H$-$^{13}C$ HSQC-edited spectrum of **4** (DMSO-$d_6$, 298 K).

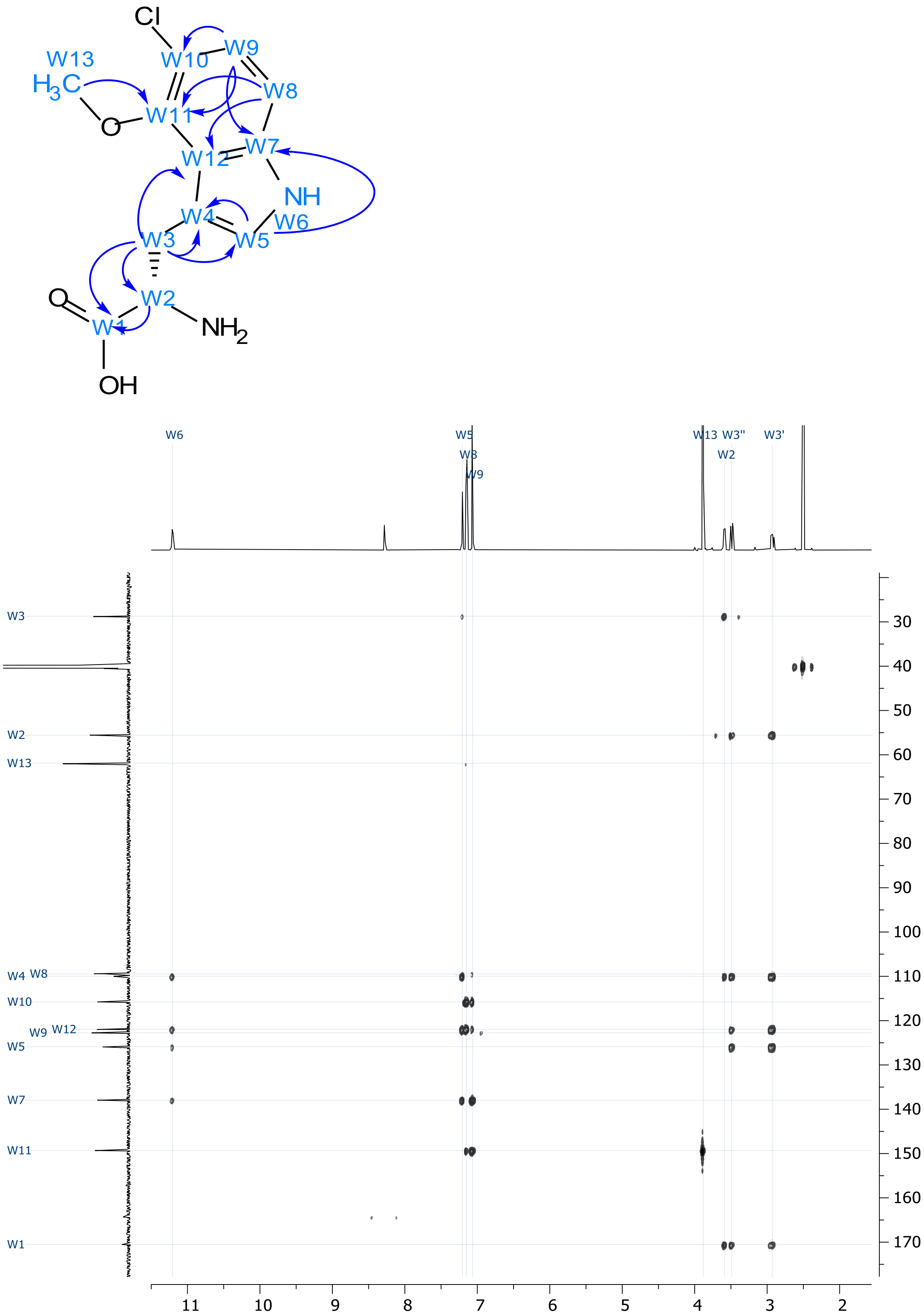


**Figure S24.** $^1$H-$^{13}$C HMBC spectrum of **4** (DMSO-d$_6$, 298 K).

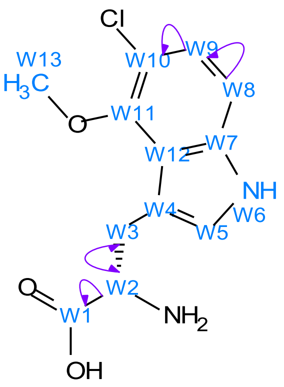

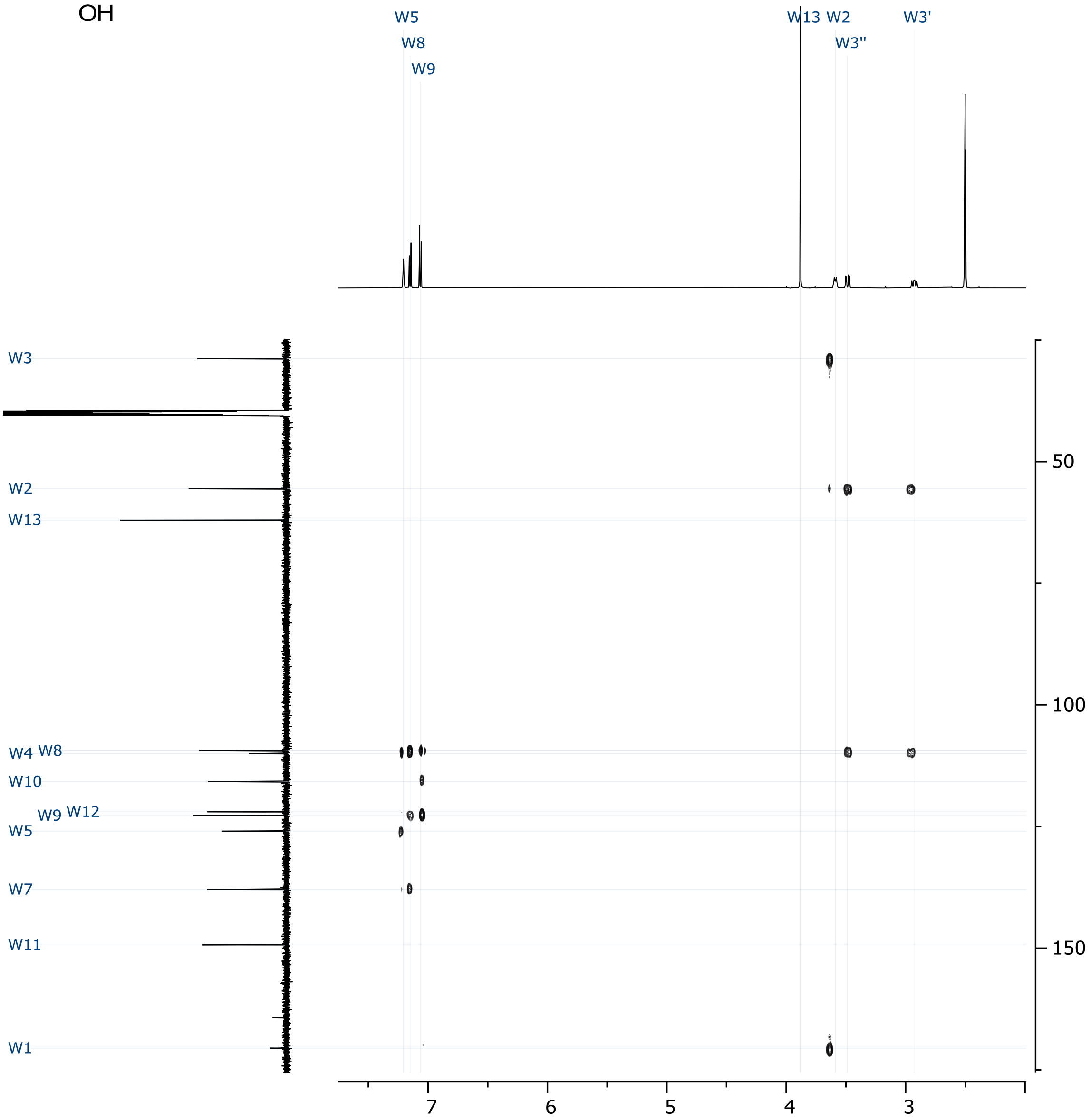


**Figure S25.** 1,1-ADEQUATE spectrum of **4** (DMSO-$d_6$, 298 K).

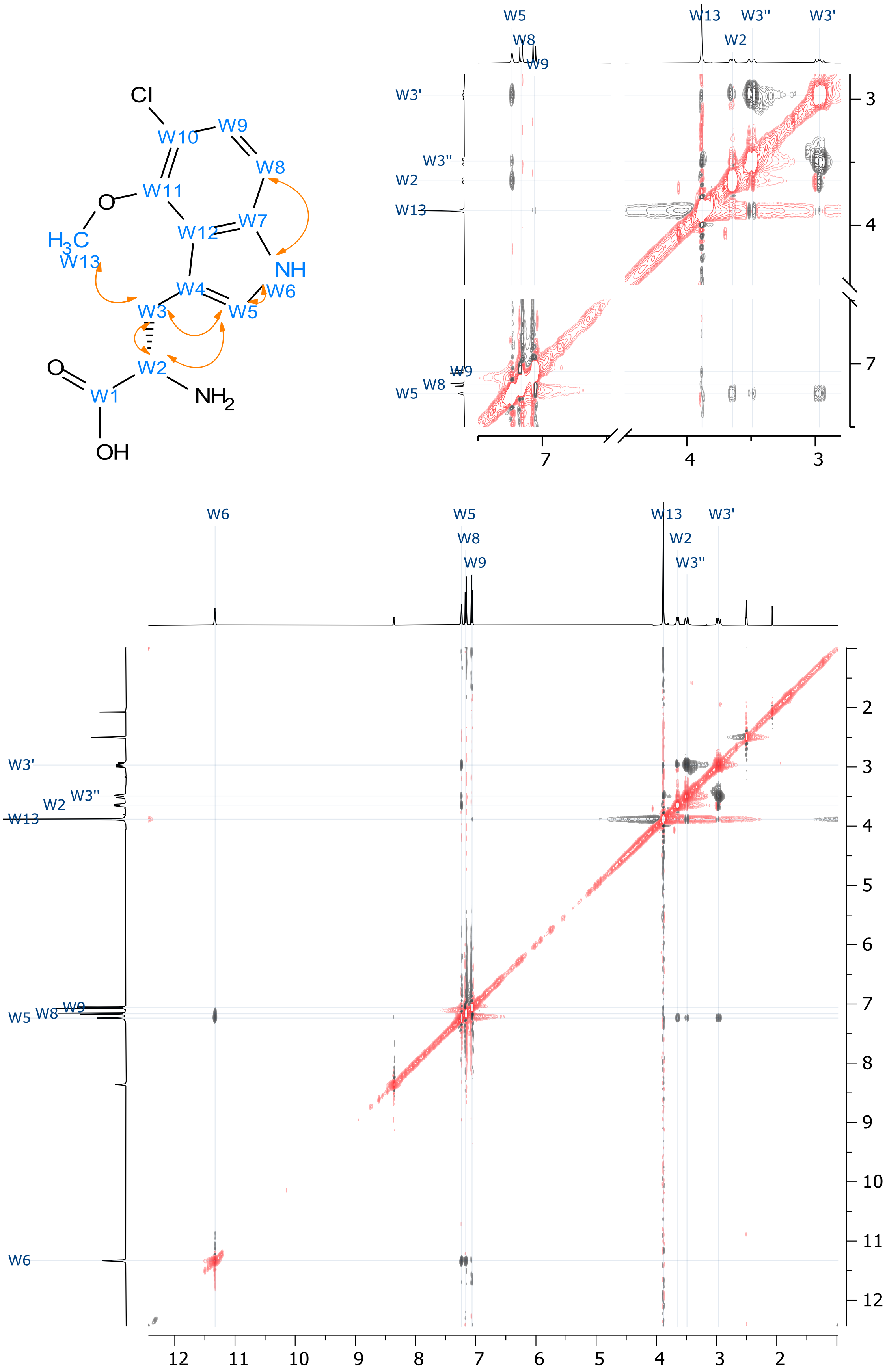


**Figure S26.** $^1H$-$^1H$ ROESY spectrum of **4** (DMSO-$d_6$, 298 K.

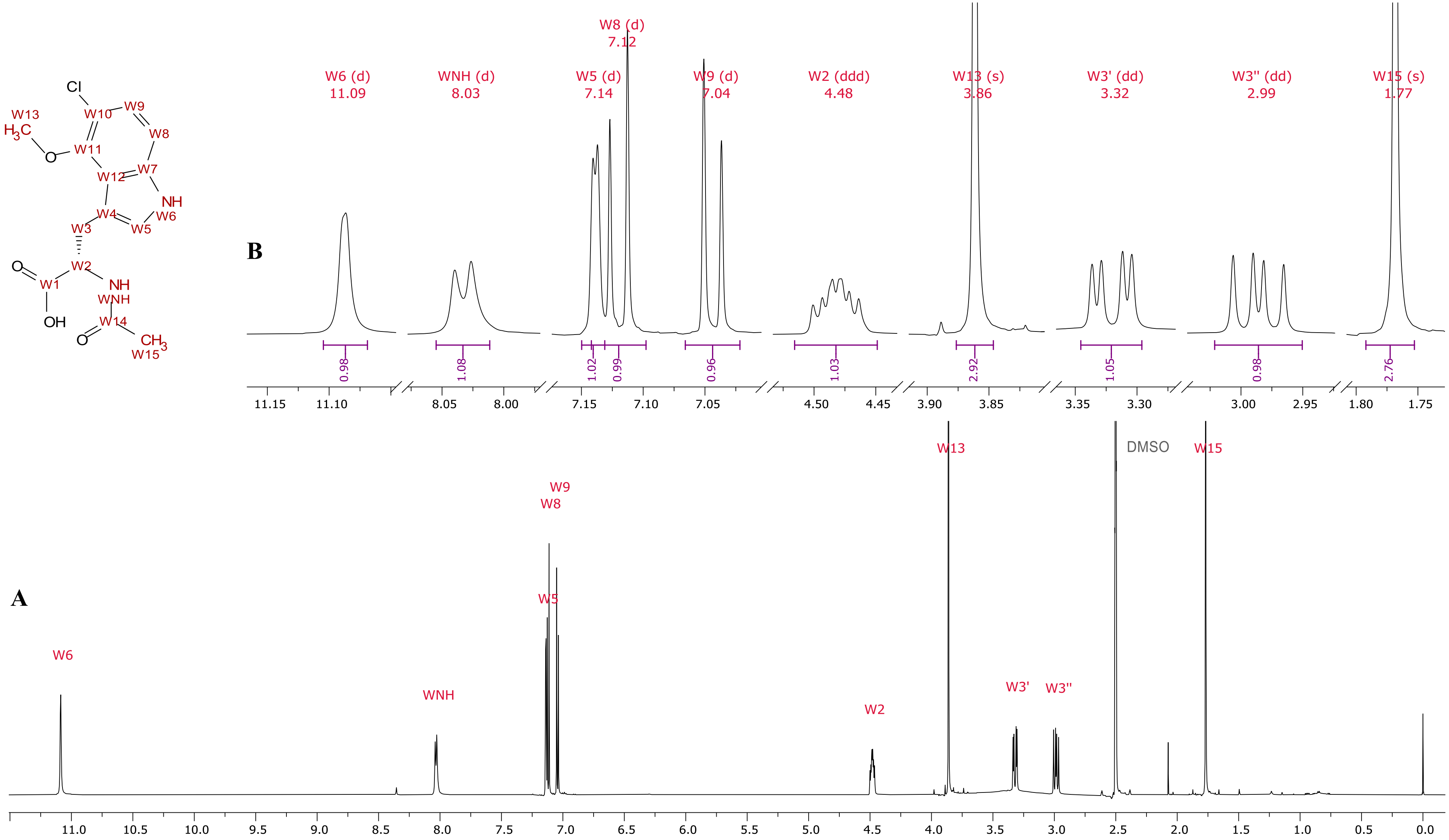


**Figure S27.** $^1$H NMR spectrum of compound **5** (DMSO-$d_6$, 600 MHz, 298 K), (a) spectrum overview and (b) expanded areas of assigned signals.

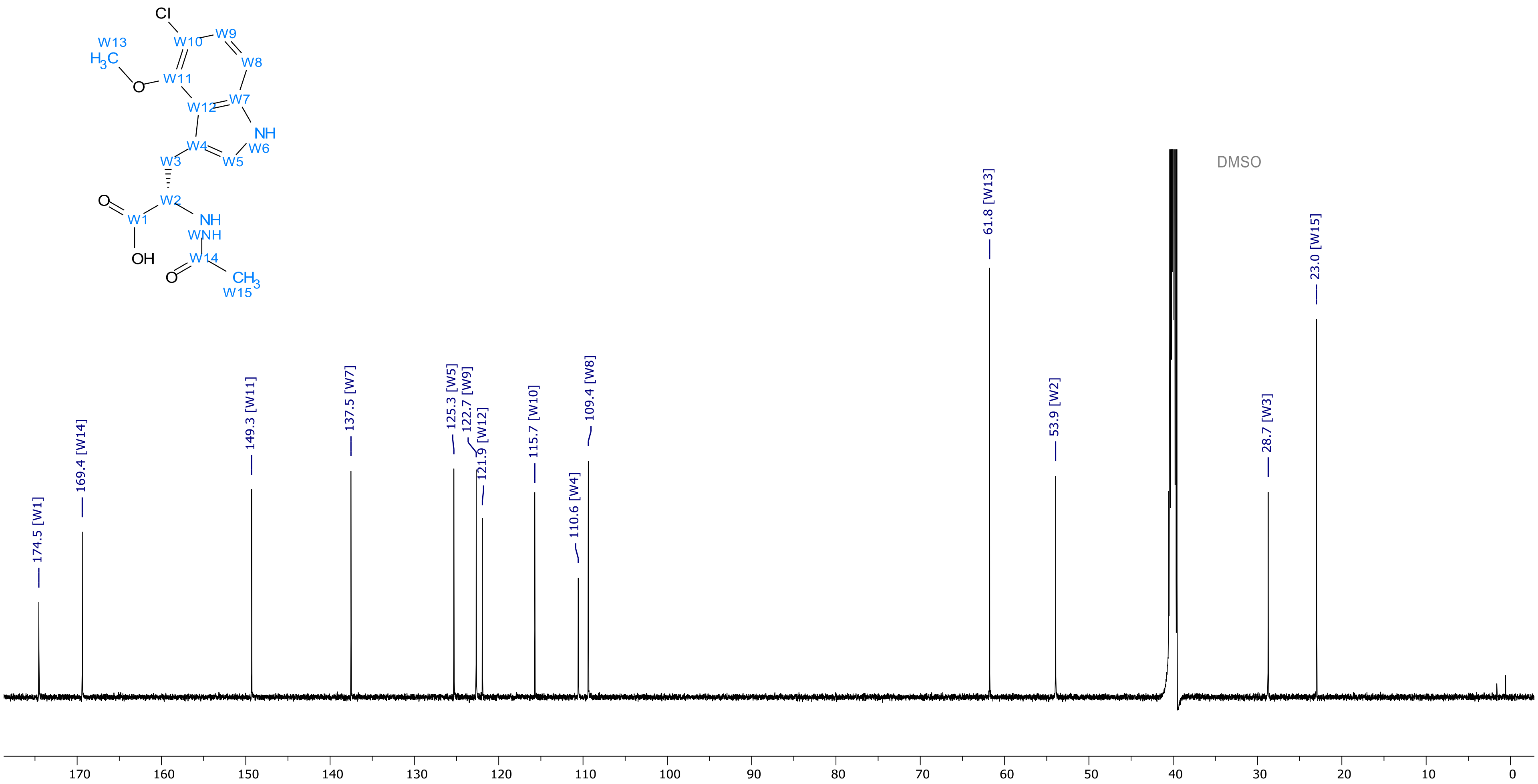


**Figure S28.** $^{13}C$ NMR of compound **5** (DMSO-$d_6$, 151 MHz, 298 K).

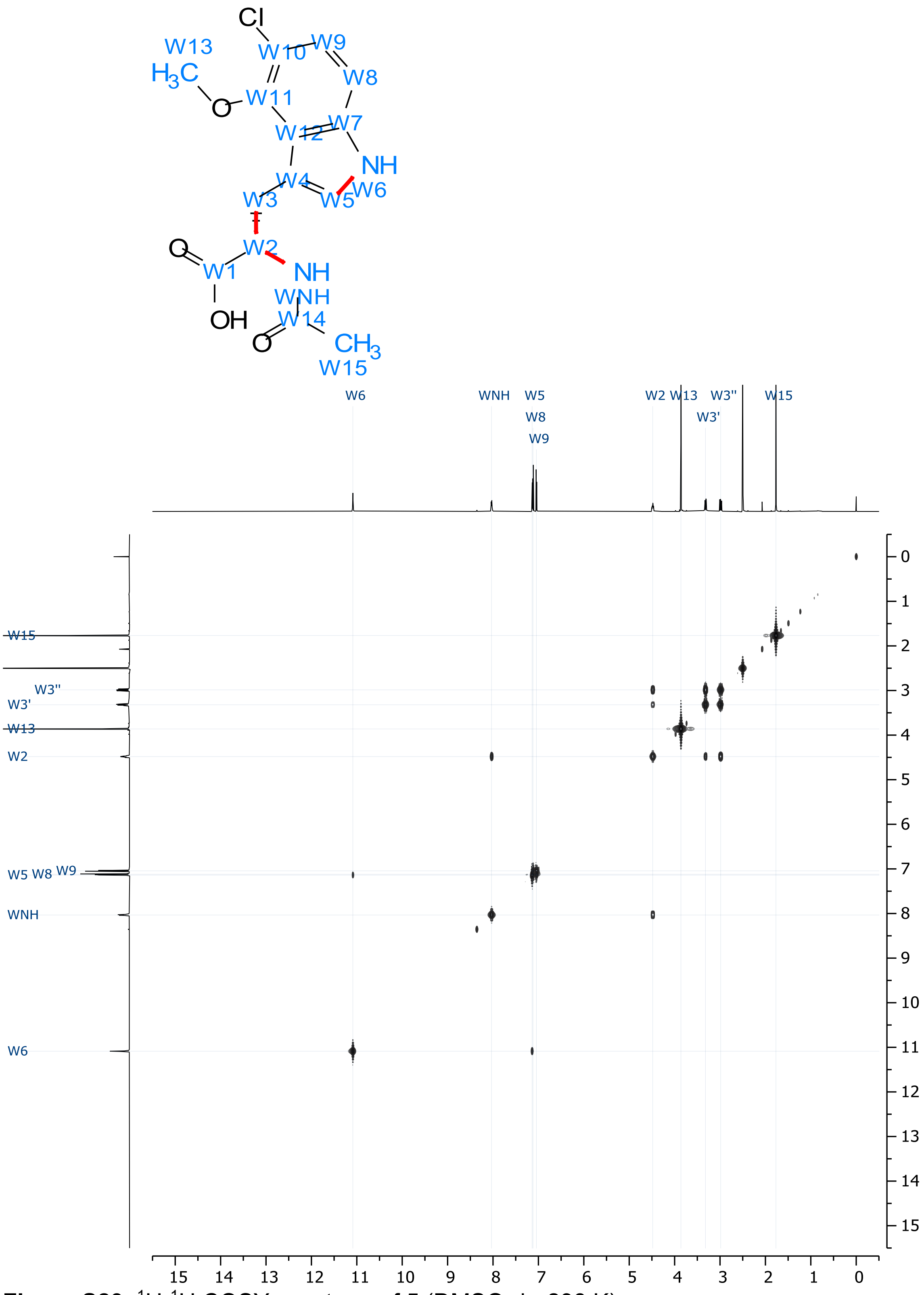


**Figure S29.** ¹H-¹H COSY spectrum of **5** (DMSO-d$_6$, 298 K).

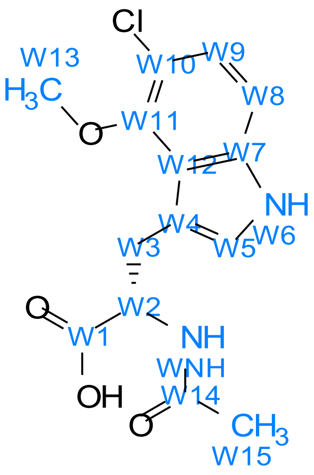


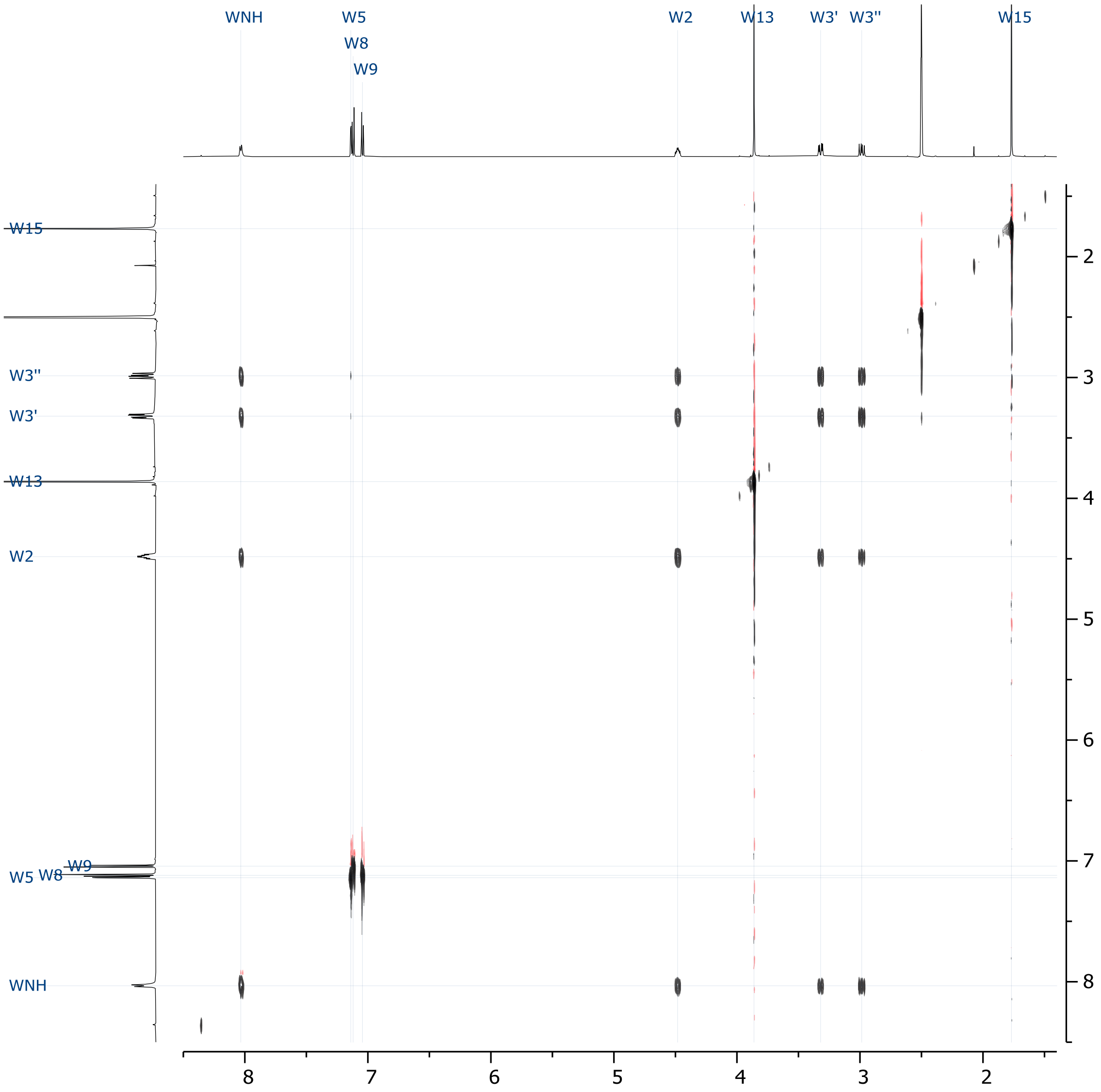


**Figure S30.** 2D TOCSY spectrum of **5** (DMSO-$d_6$, 298 K).

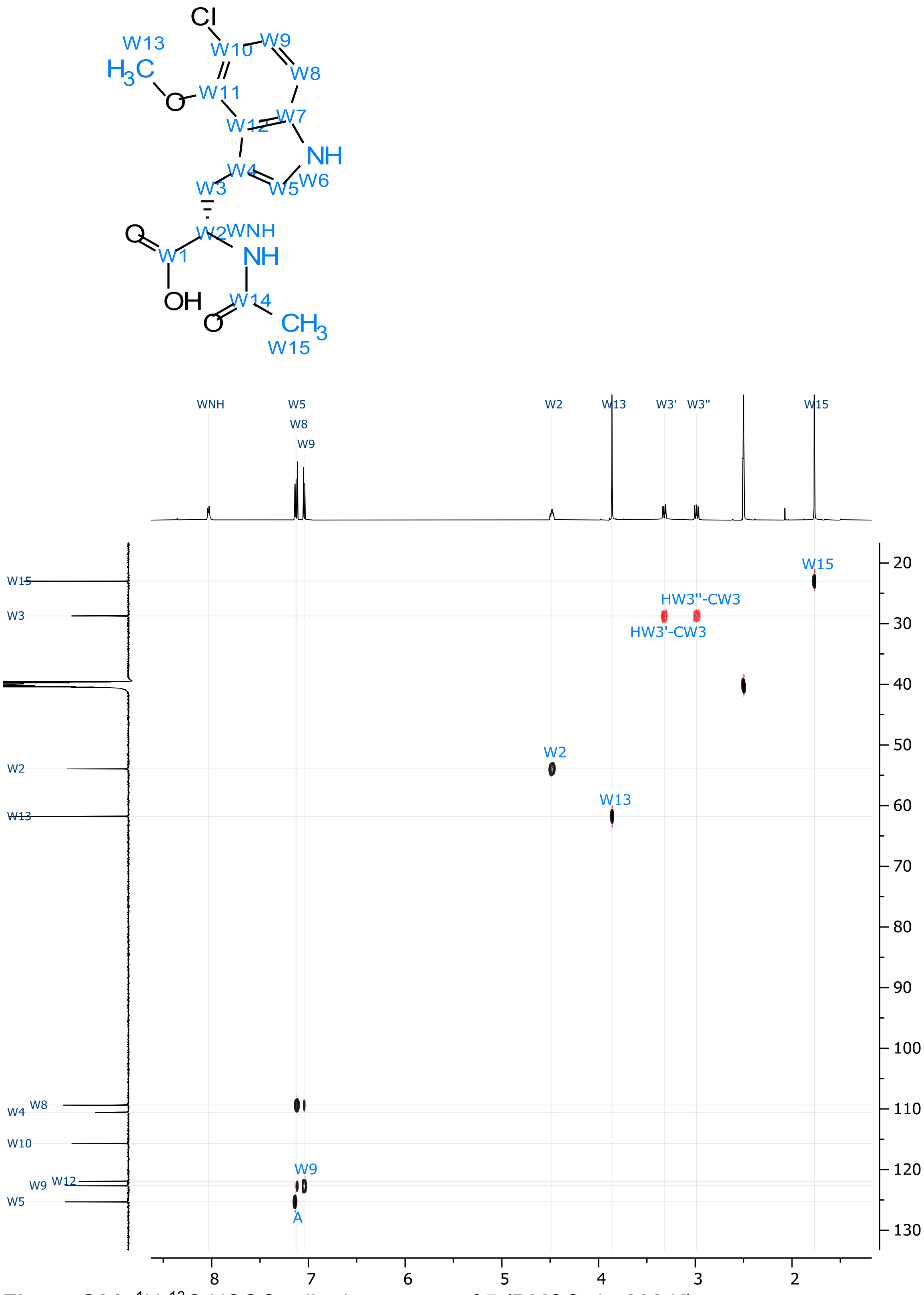


**Figure S31.** $^1H$-$^{13}C$ HSQC-edited spectrum of **5** (DMSO-$d_6$, 298 K).

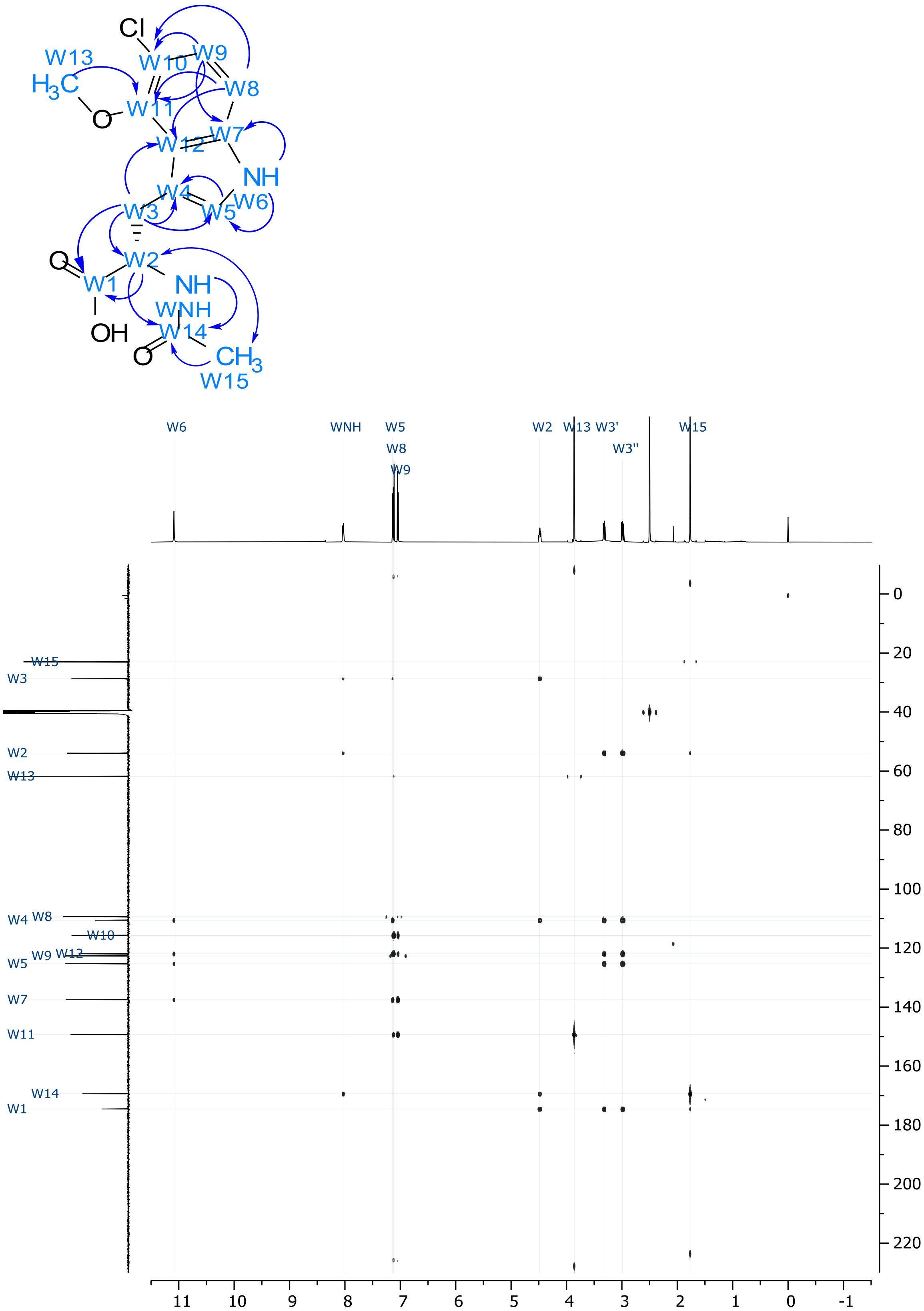


**Figure S32.** $^{1}H$-$^{13}C$ HMBC spectrum of **5** (DMSO-$d_6$, 298 K).

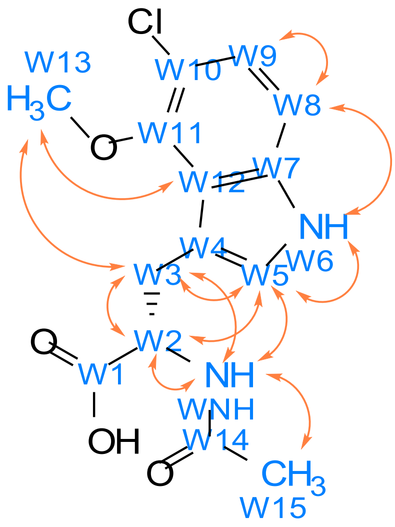


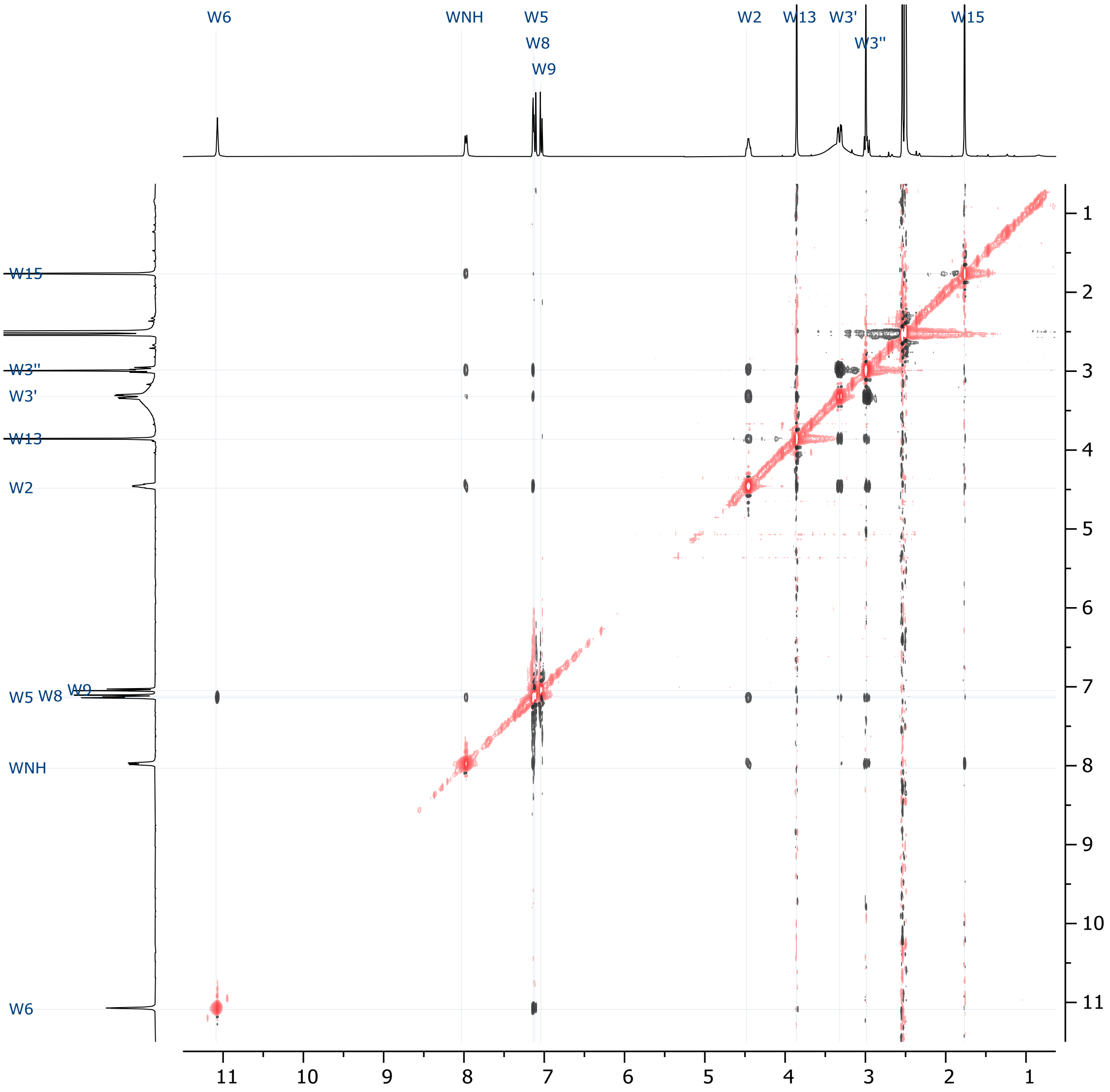


**Figure S33.** $^{1}$H-$^{1}$H ROSEY spectrum of **5** (DMSO-d$_6$, 298 K).

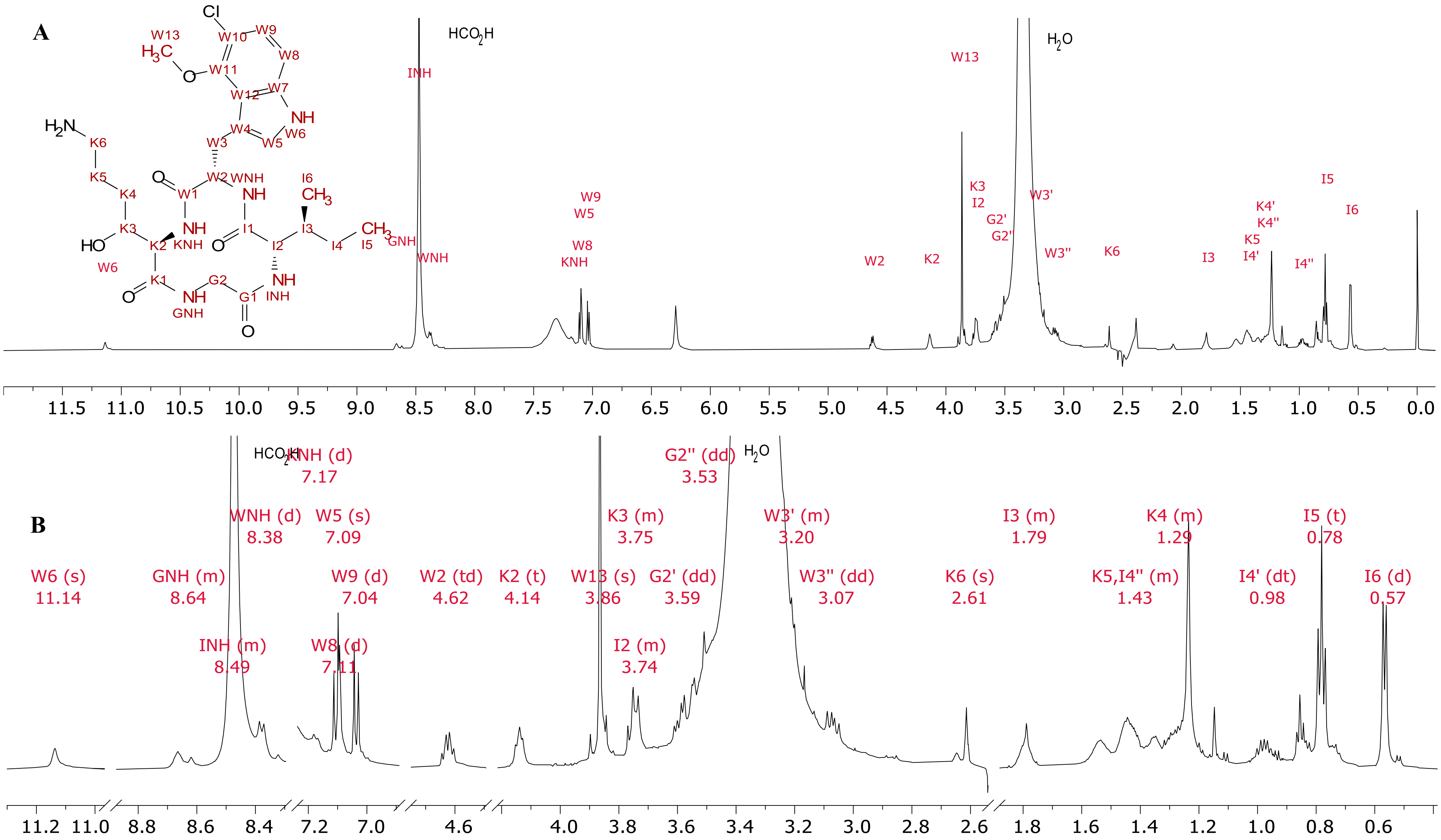


**Figure S34.** [1]H NMR of **6** (DMSO-$d_6$, 600 MHz, 298 K) recorded with suppression (Bruker noesypresat experiment) of the residual solvent (DMSO-d6). (b) The spectrum overview and (b) an expansion of assigned signals.

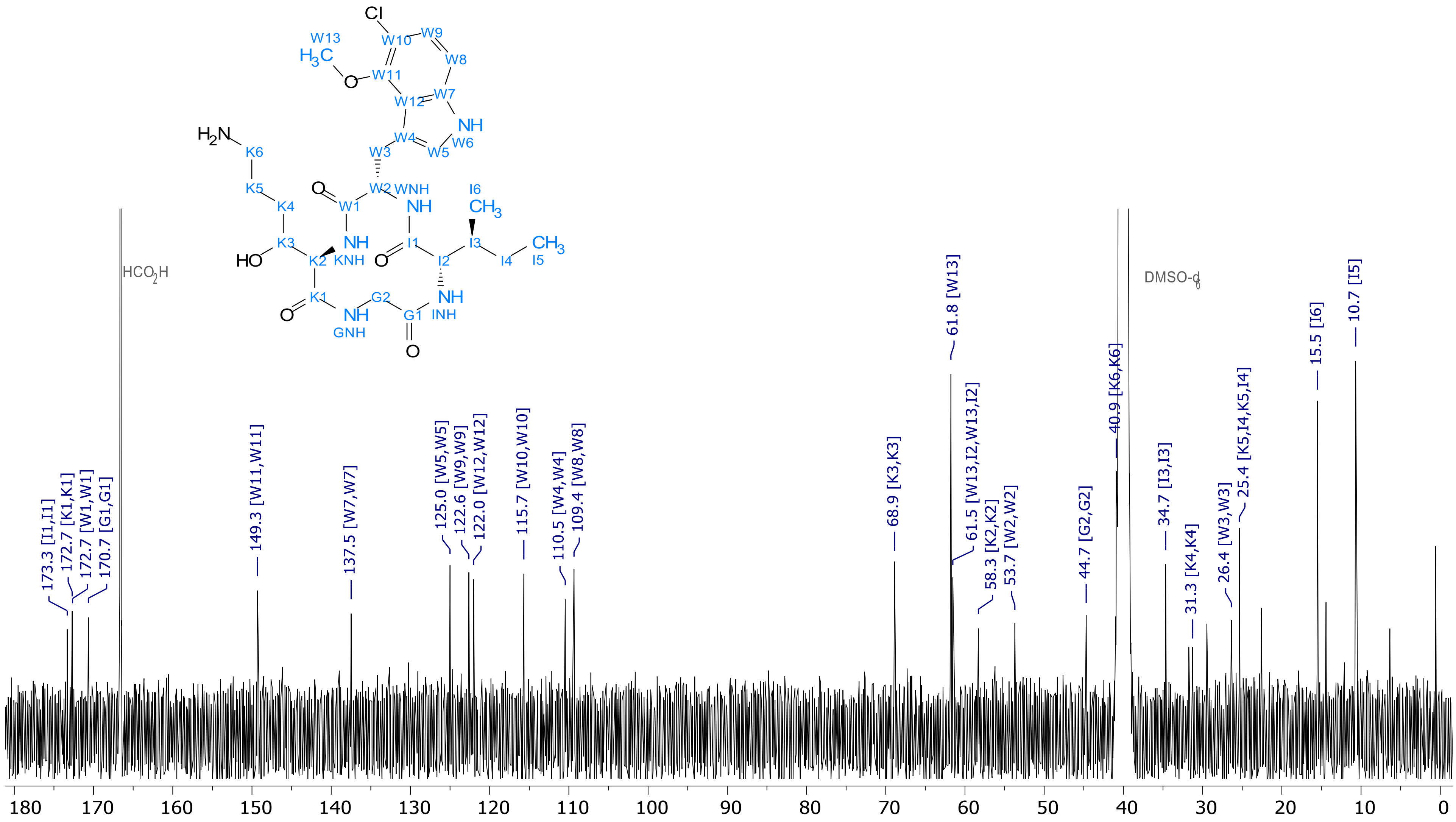


**Figure S35.** $^{13}$C NMR of **6** (DMSO-d$_6$, 151 MHz, 298 K).

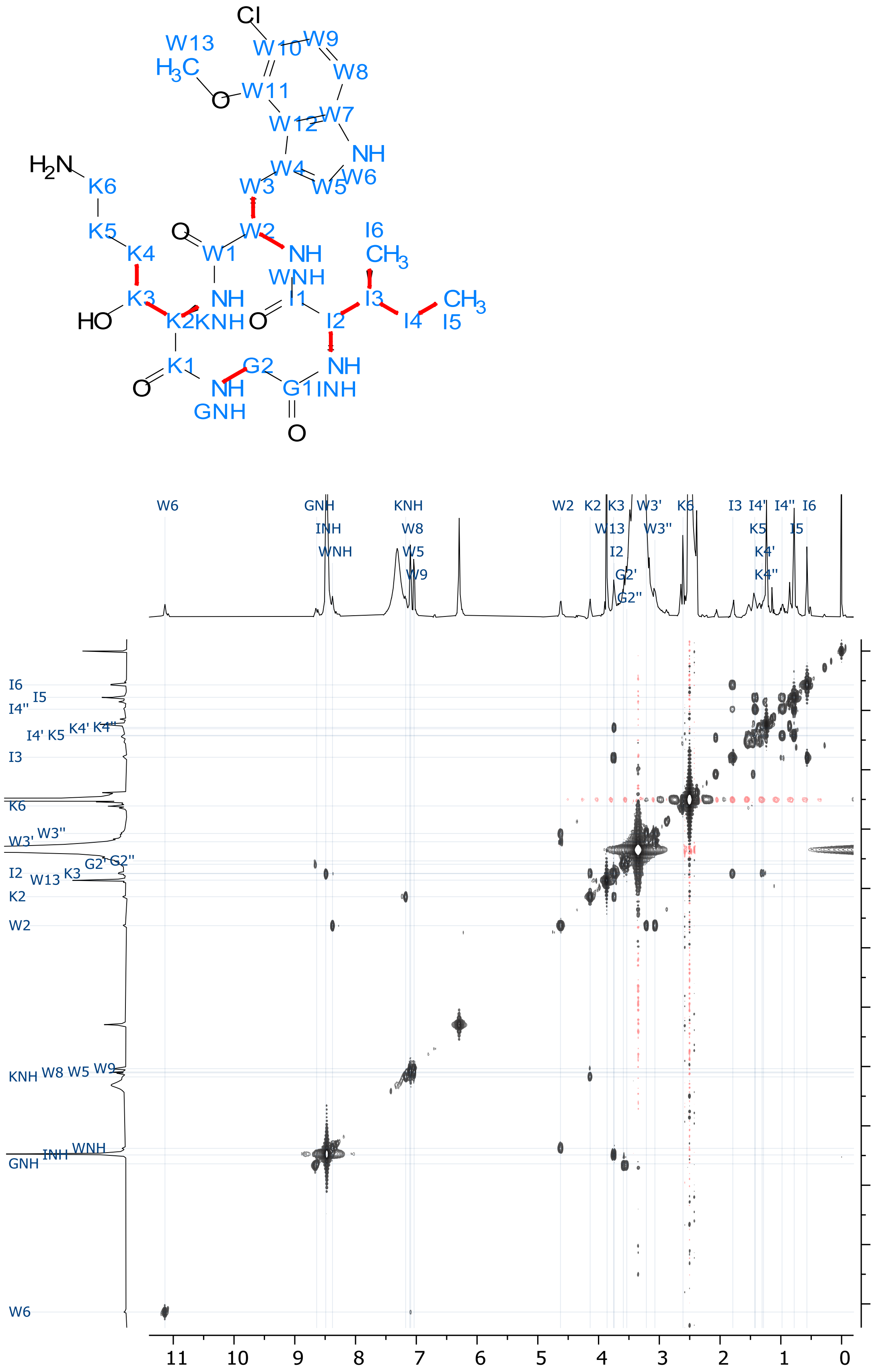


**Figure S36.** $^{1}H$-$^{1}H$ COSY spectrum of **6** (DMSO-$d_6$, 298 K).

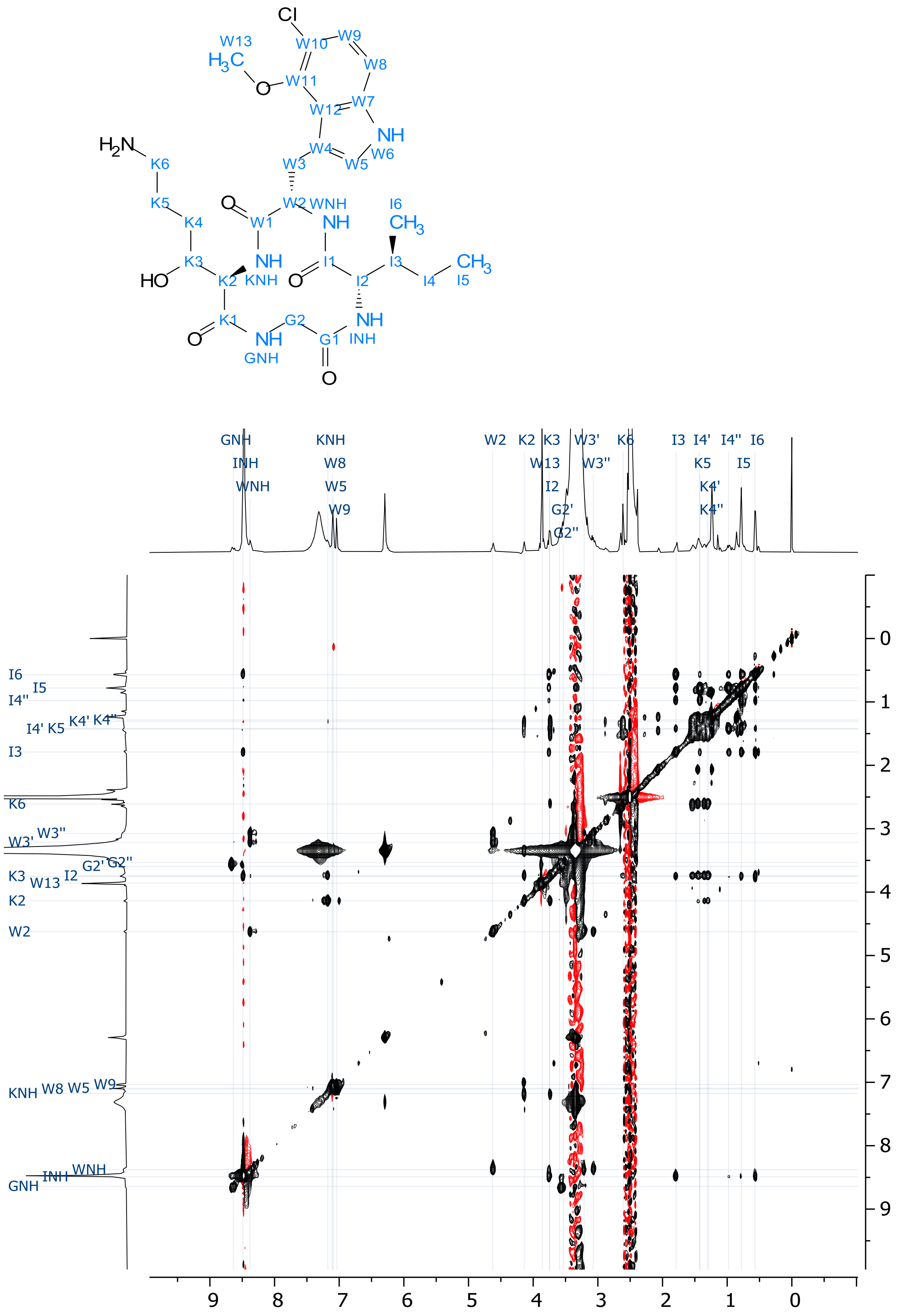


**Figure S37.** 2D TOCSY NMR spectrum of **6** (DMSO-d₆, 298 K).

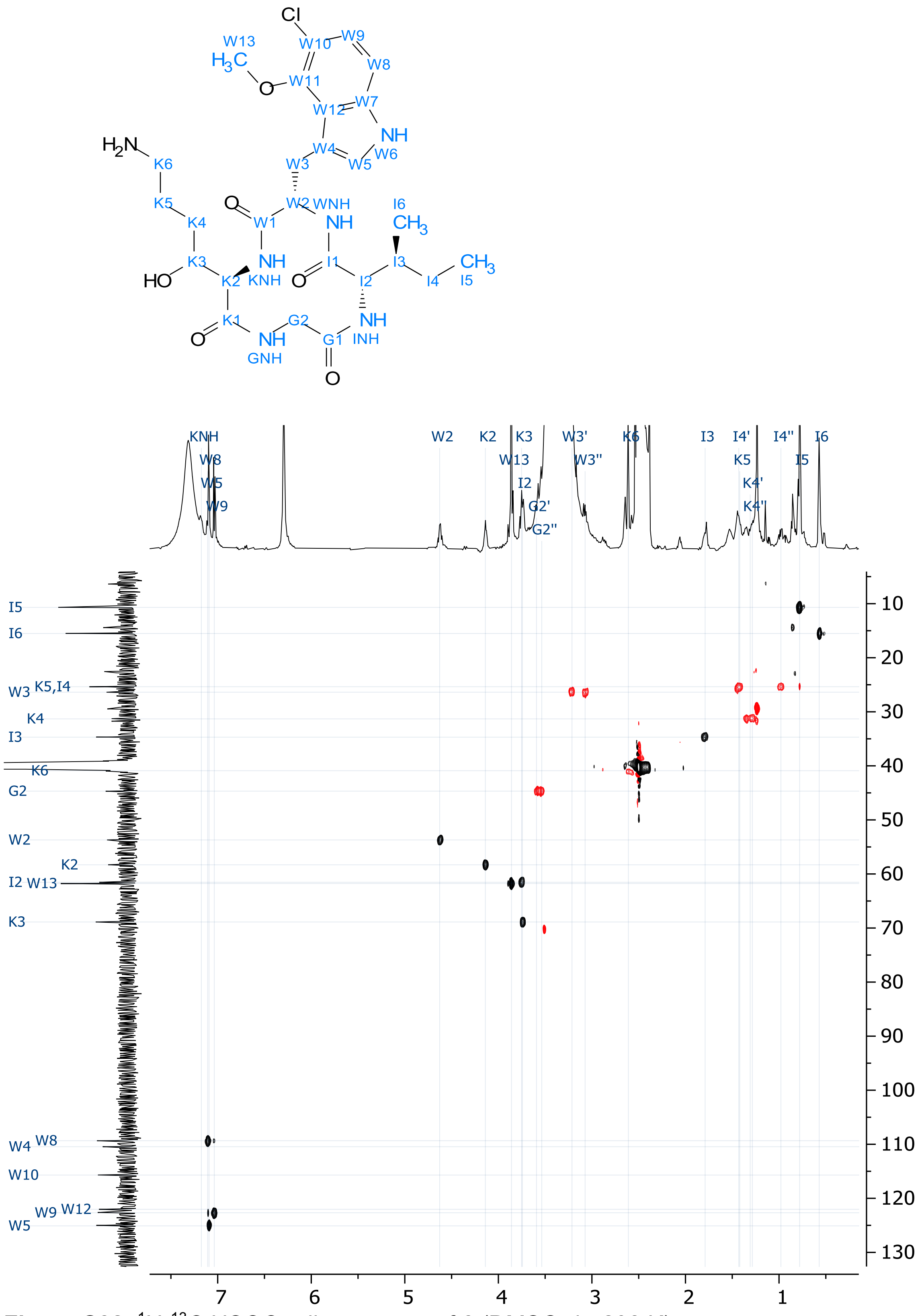


**Figure S38.** $^{1}H$-$^{13}C$ HSQC-edit spectrum of **6** (DMSO-$d_6$, 298 K).

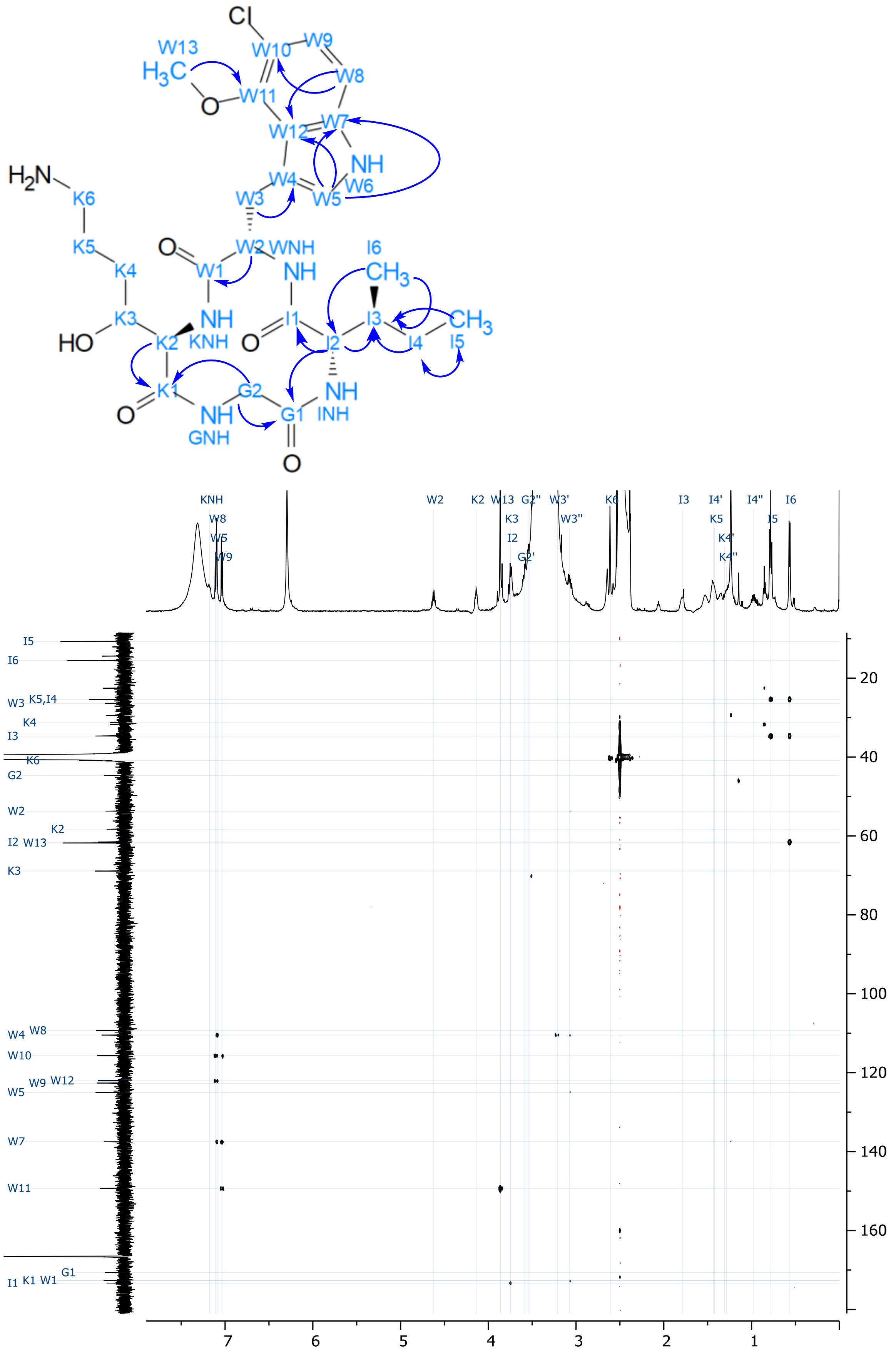

**Figure S39.** $^{1}H$-$^{13}C$ HMBC spectrum of **6** (DMSO-$d_6$, 298 K).

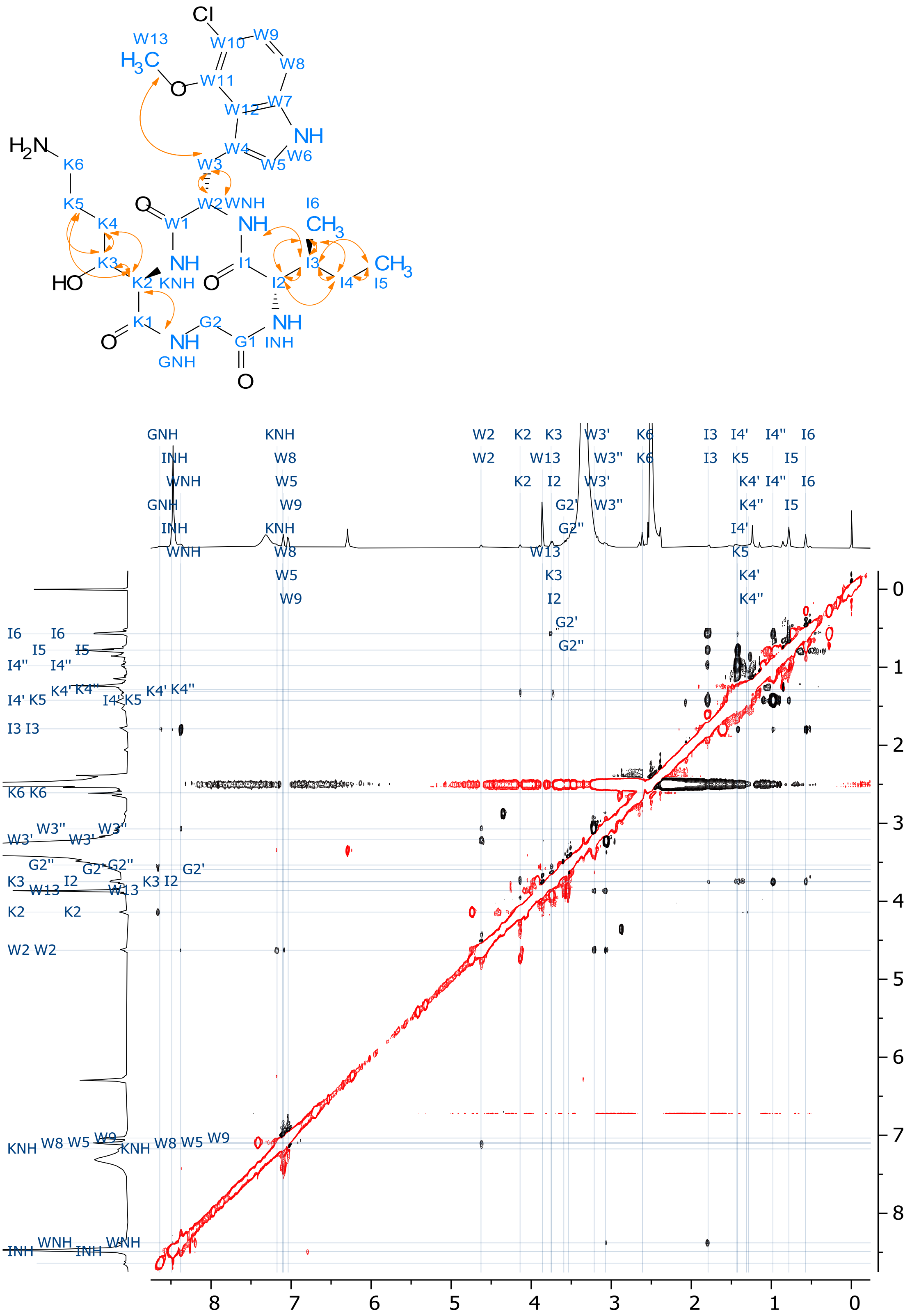


**Figure S40.** $^{1}H$-$^{1}H$ ROESY spectrum of **6** (DMSO-$d_6$, 298 K).

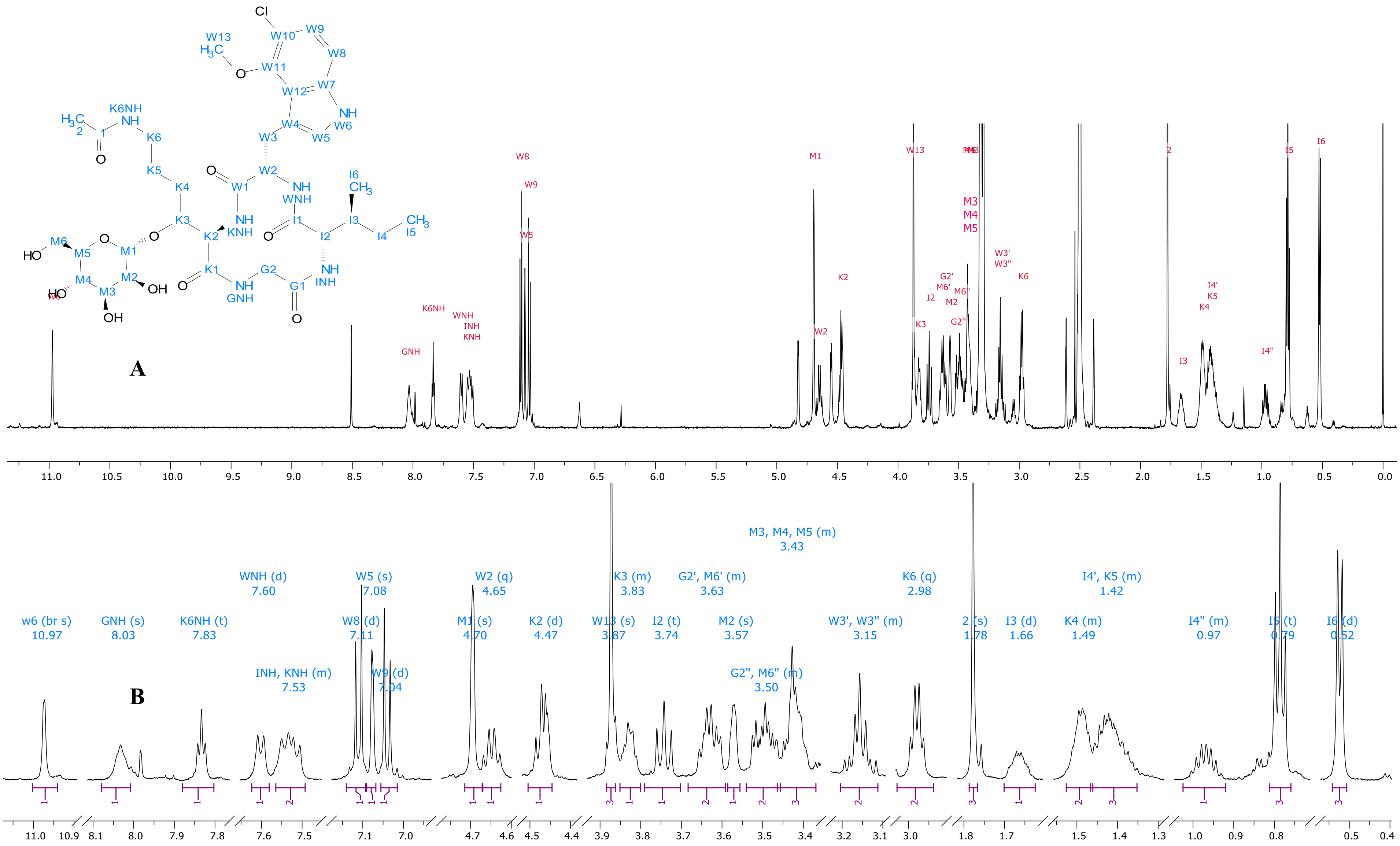

**Figure S41.** $^1H$ NMR spectra of compound **7** (DMSO-$d_6$, 600 MHz, 298 K). (a) Spectrum overview and (b) expansions of signal multiplets. Unlabelled resonances correspond to an unidentified impurity.

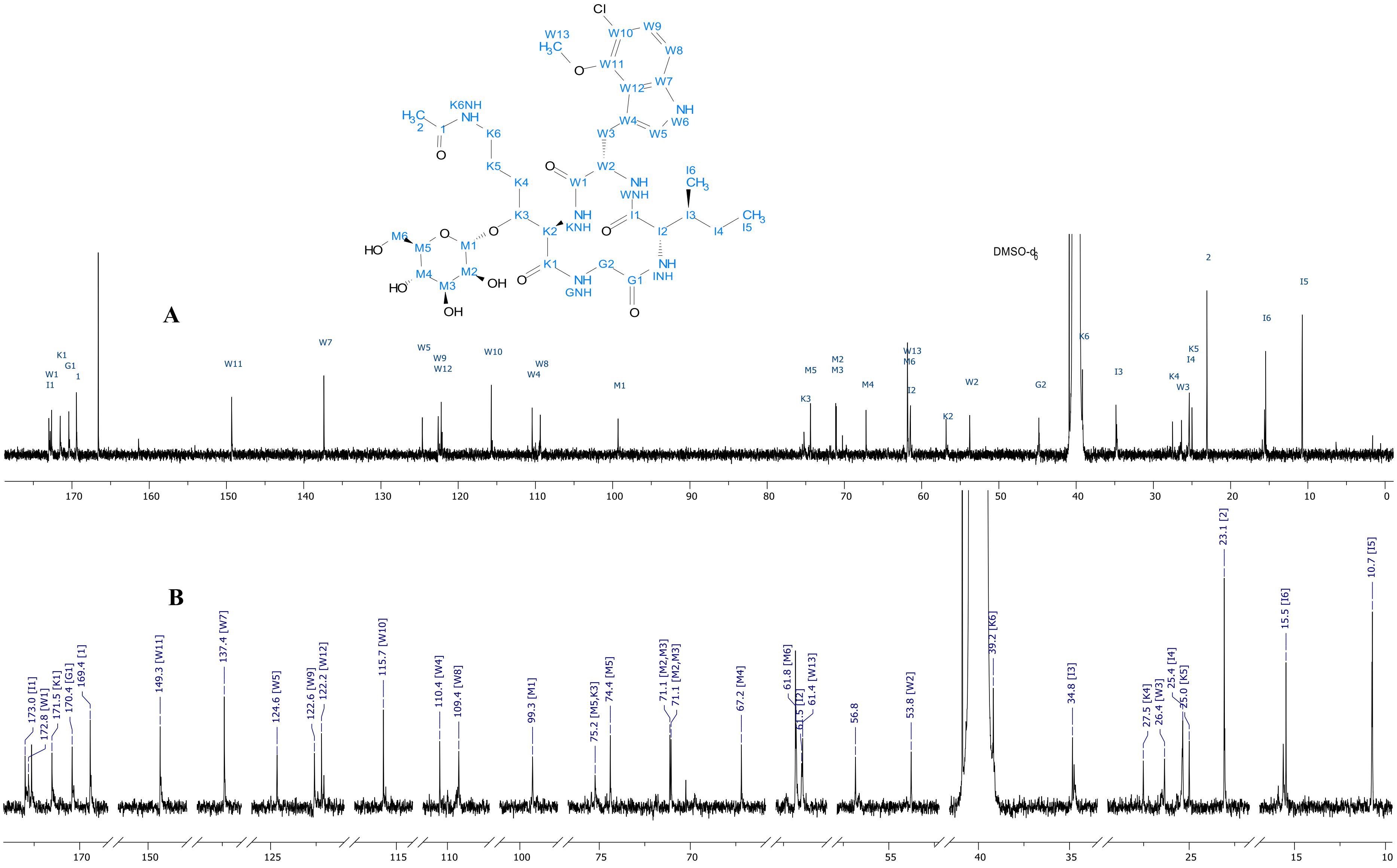


**Figure S42**. $^{13}$C NMR spectra of Compound **7** (DMSO-d6, 151 MHz, 298 K). (a) Spectrum overview and (b) expansion of spectrum fragments with labelled signals.

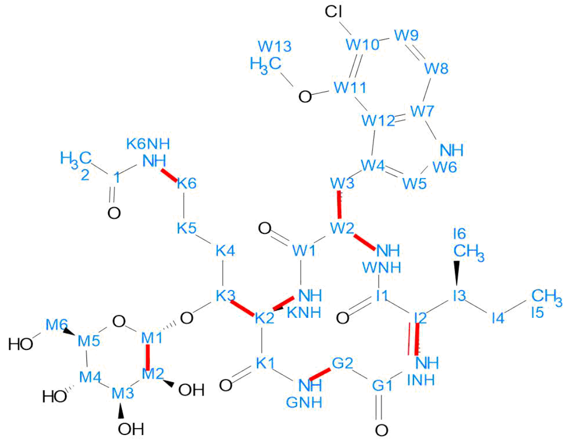


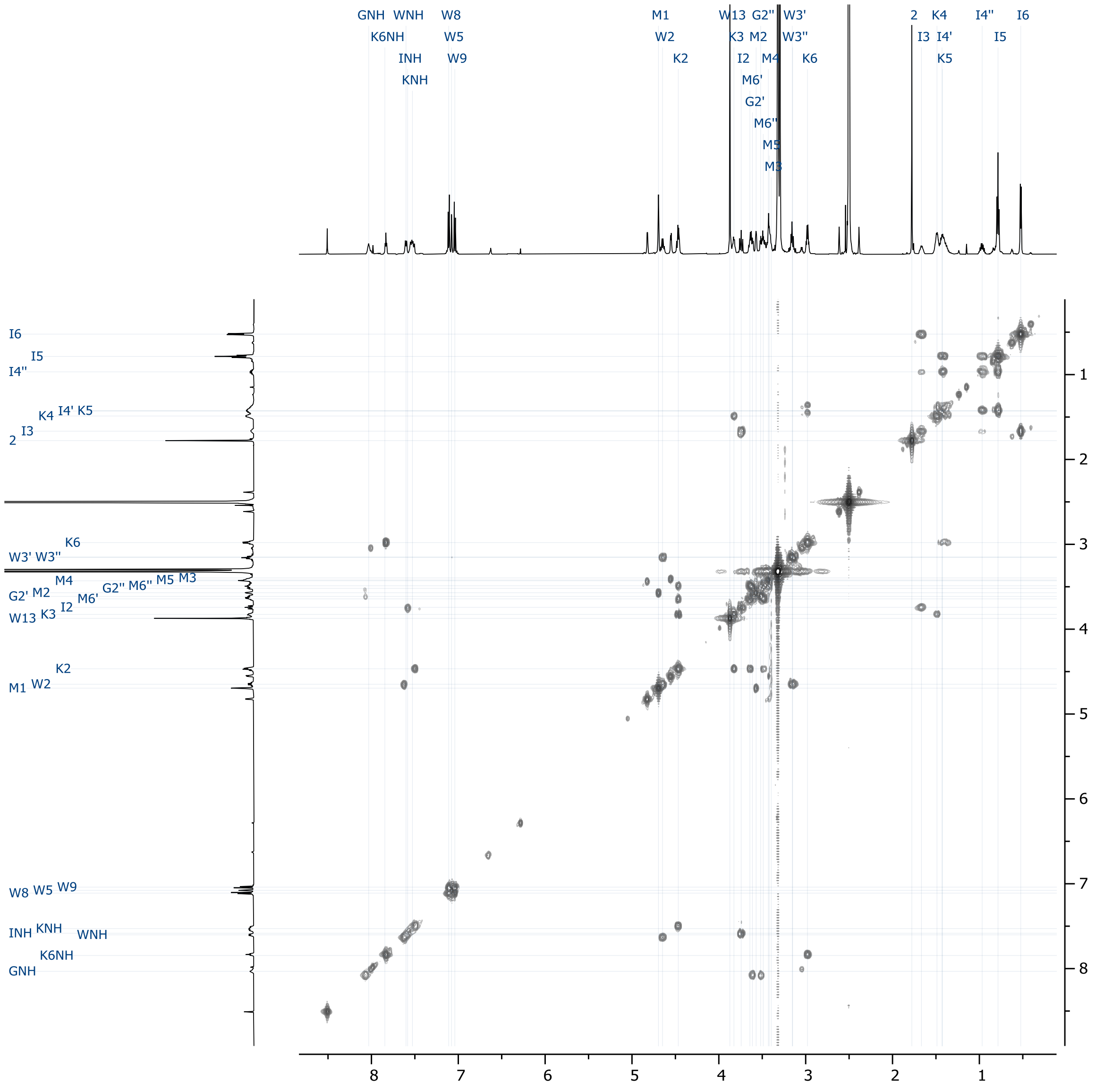


**Figure S43**. 2D COSY spectrum of compound **7** (DMSO-$d_6$ at 298 K).

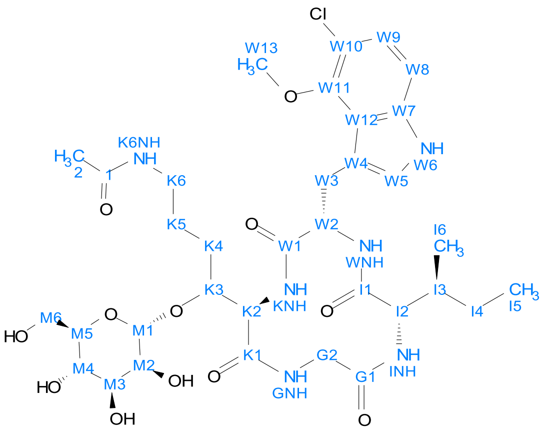

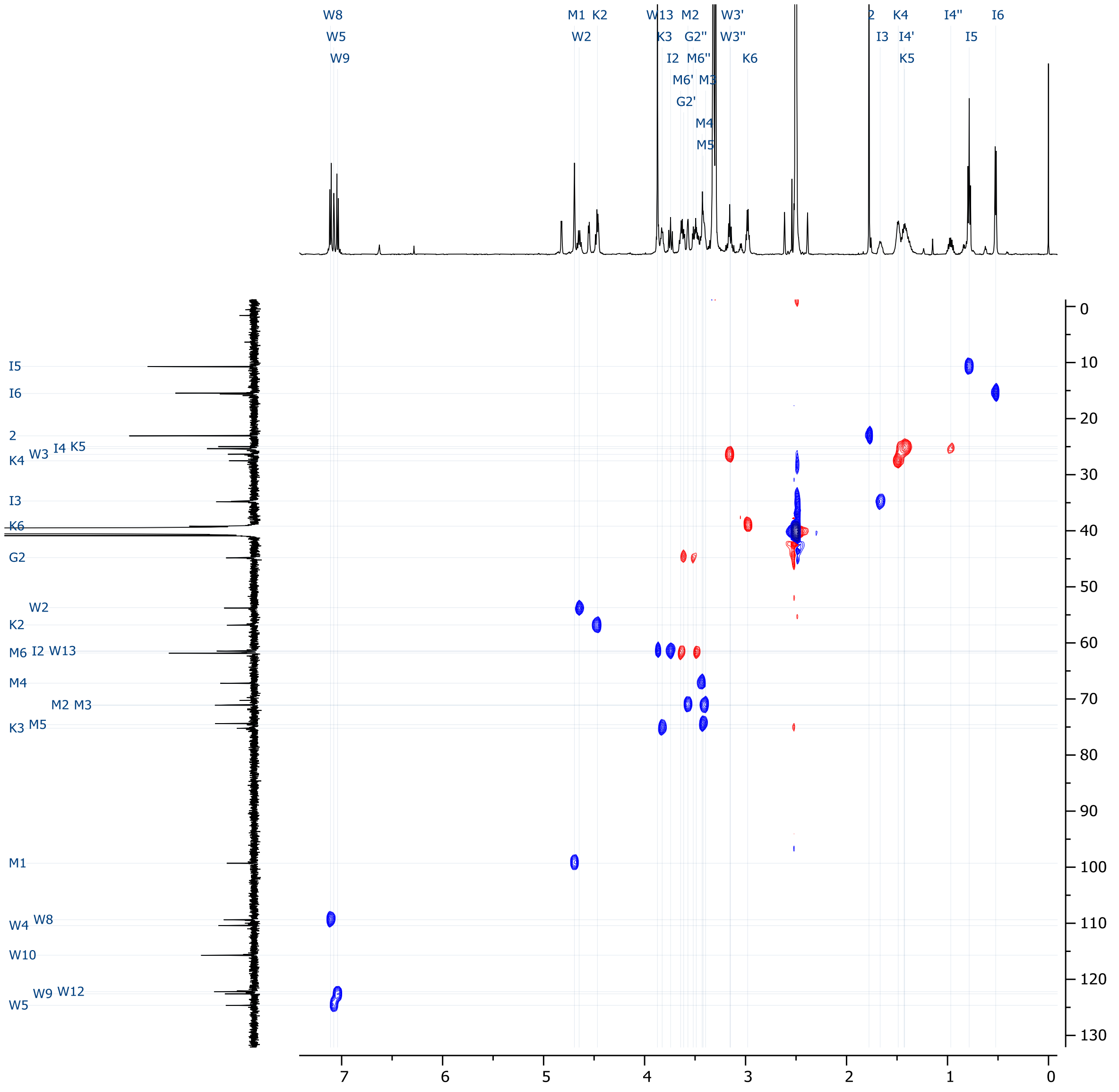

**Figure S44**. 2D HSQC-EDITED spectrum of compound **7** (DMSO-$d_6$ at 298 K).

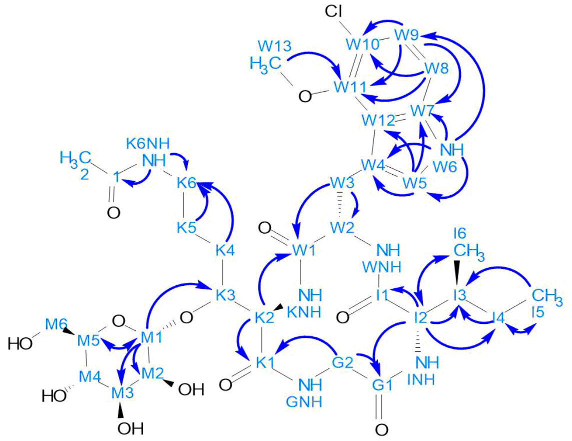

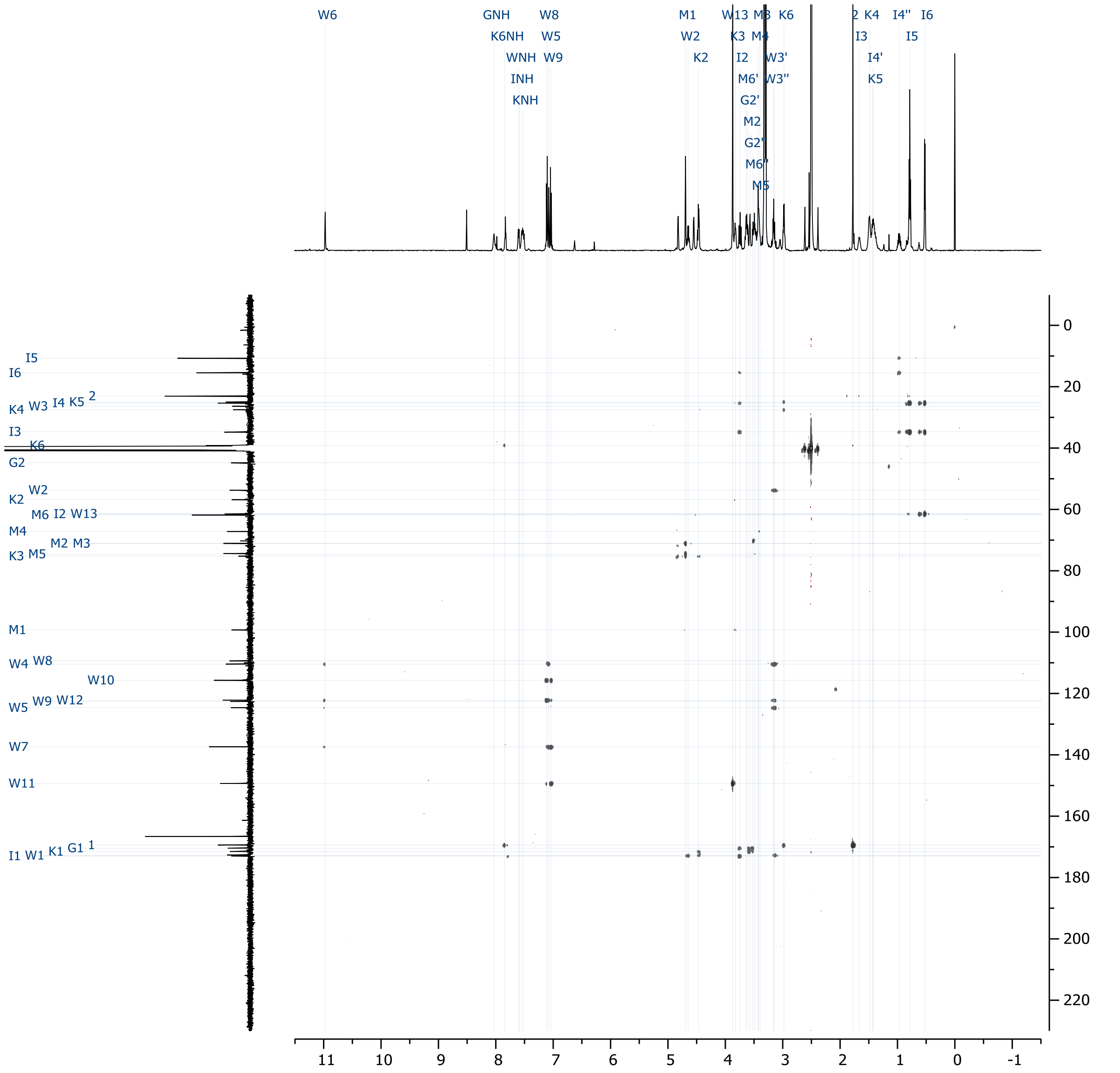

**Figure S45**. 2D HMBC spectrum of compound **7** (DMSO-$d_6$ at 298 K).

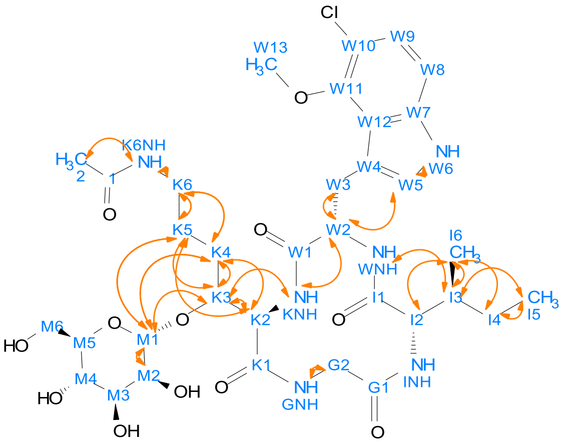

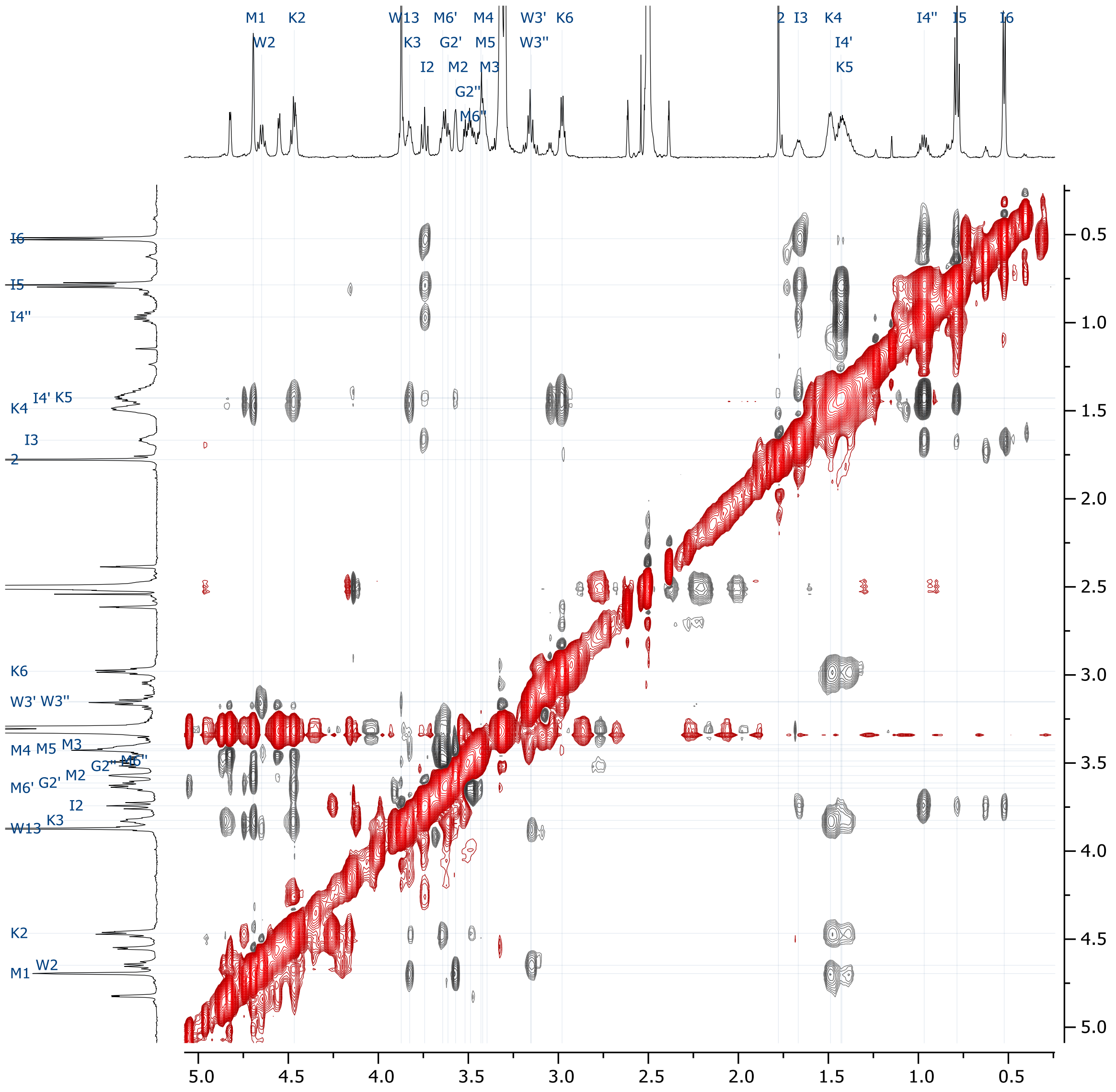


**Figure S46**. 2D ROESY spectrum of compound **7** (DMSO-$d_6$ at 298 K).

**A**

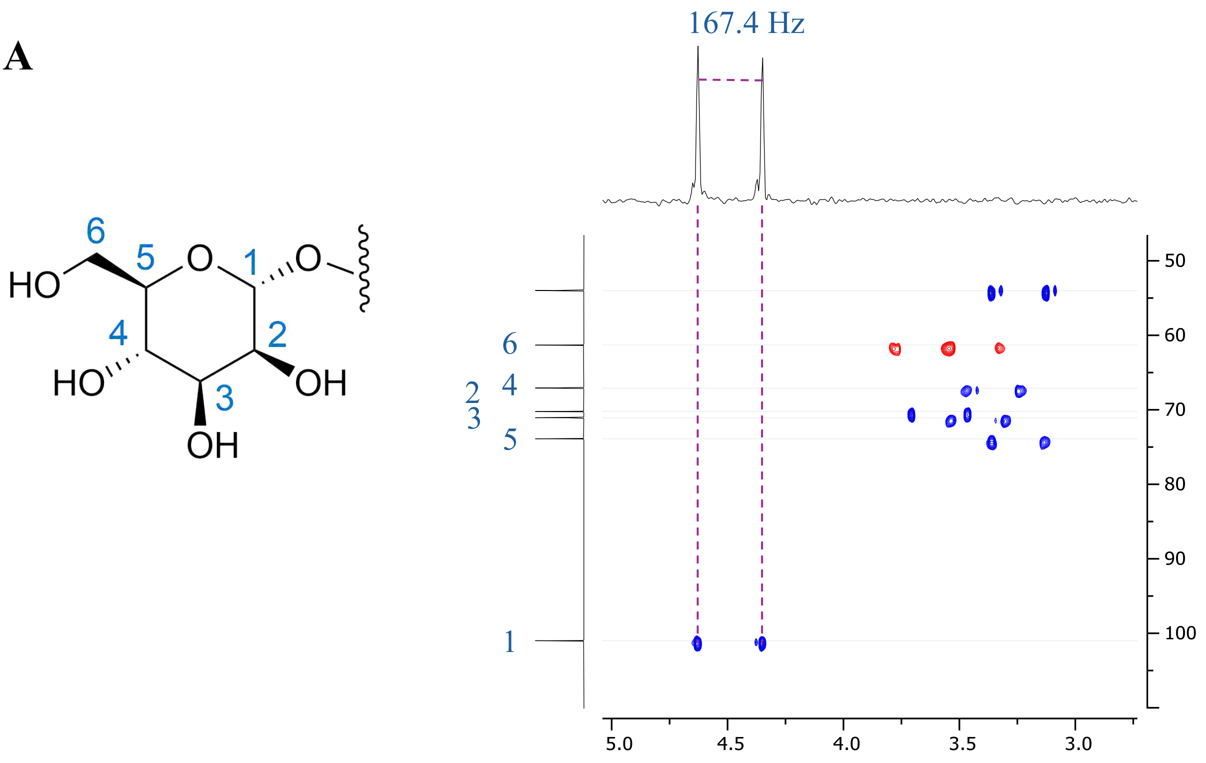


**B**

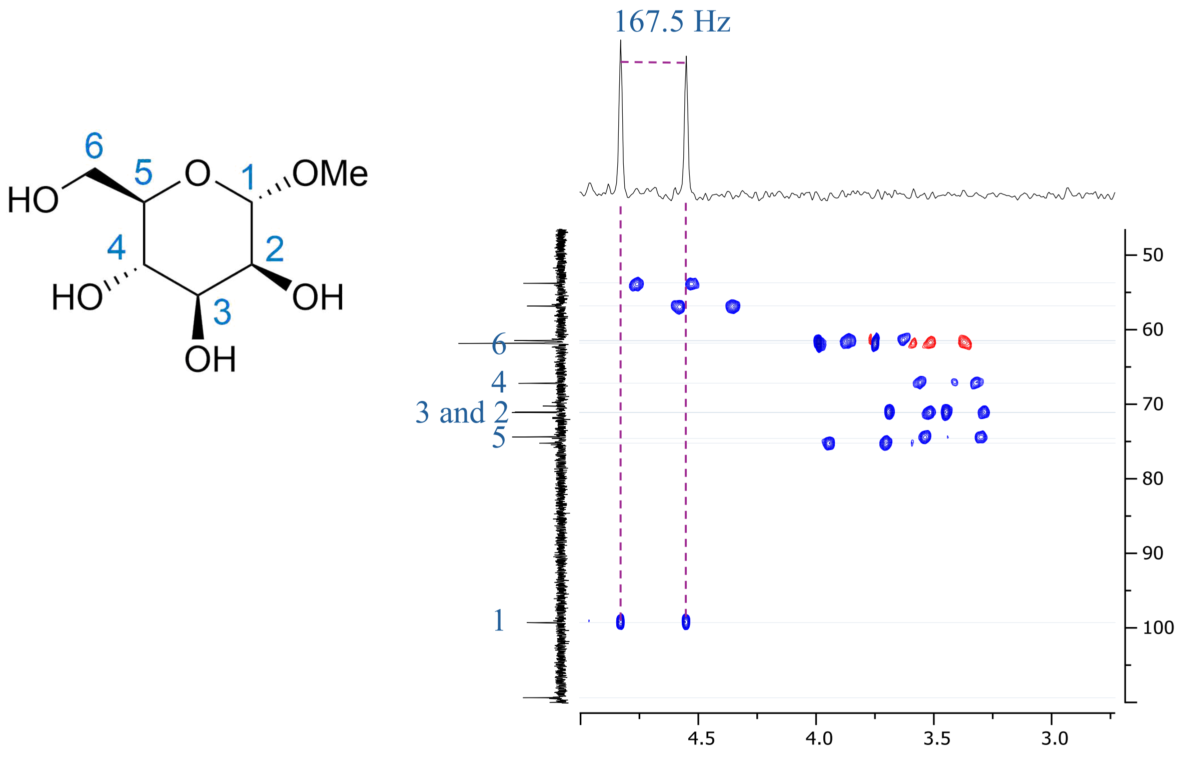


**Figure S47**. Sections of coupled 2D HSQC spectra (DMSO-$d_6$ at 298 K) of (a) compound **7** and (b) methyl α-D-mannopyranoside containing cross-peaks of the mannose residue[58]. Projections in $^1$H dimension are generated from cross peaks of mannose C1 show the splitting corresponding to the size of $^1J_{C,H}$ coupling constants.

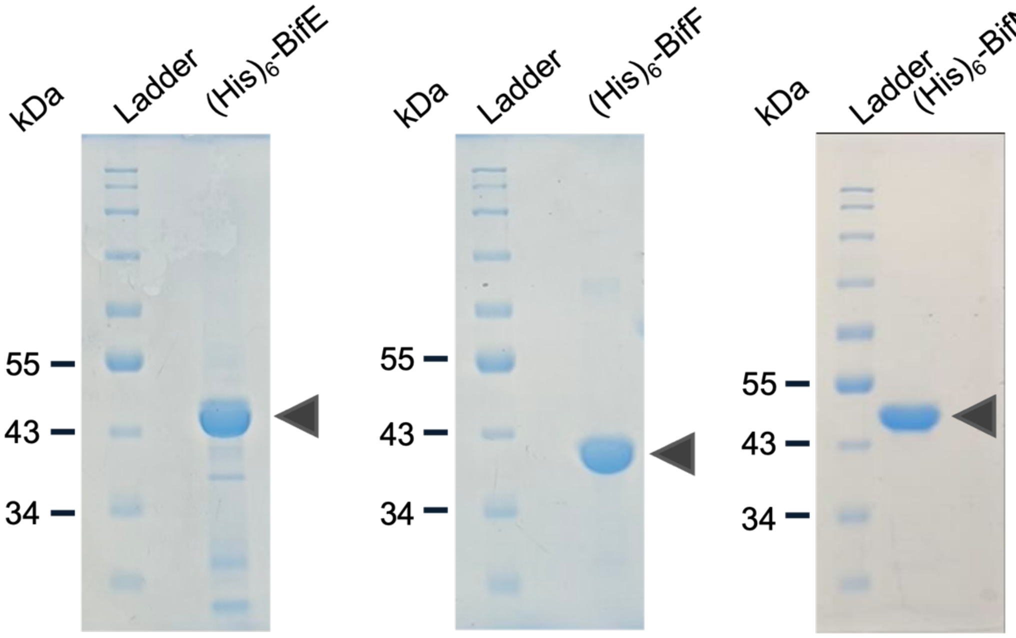


**Figure S48.** Purification and analysis of recombinant proteins. SDS-PAGE analysis of purified $(His)_6$-BifE, $(His)_6$-BifF, $(His)_6$-BifN as indicated by the black triangle.

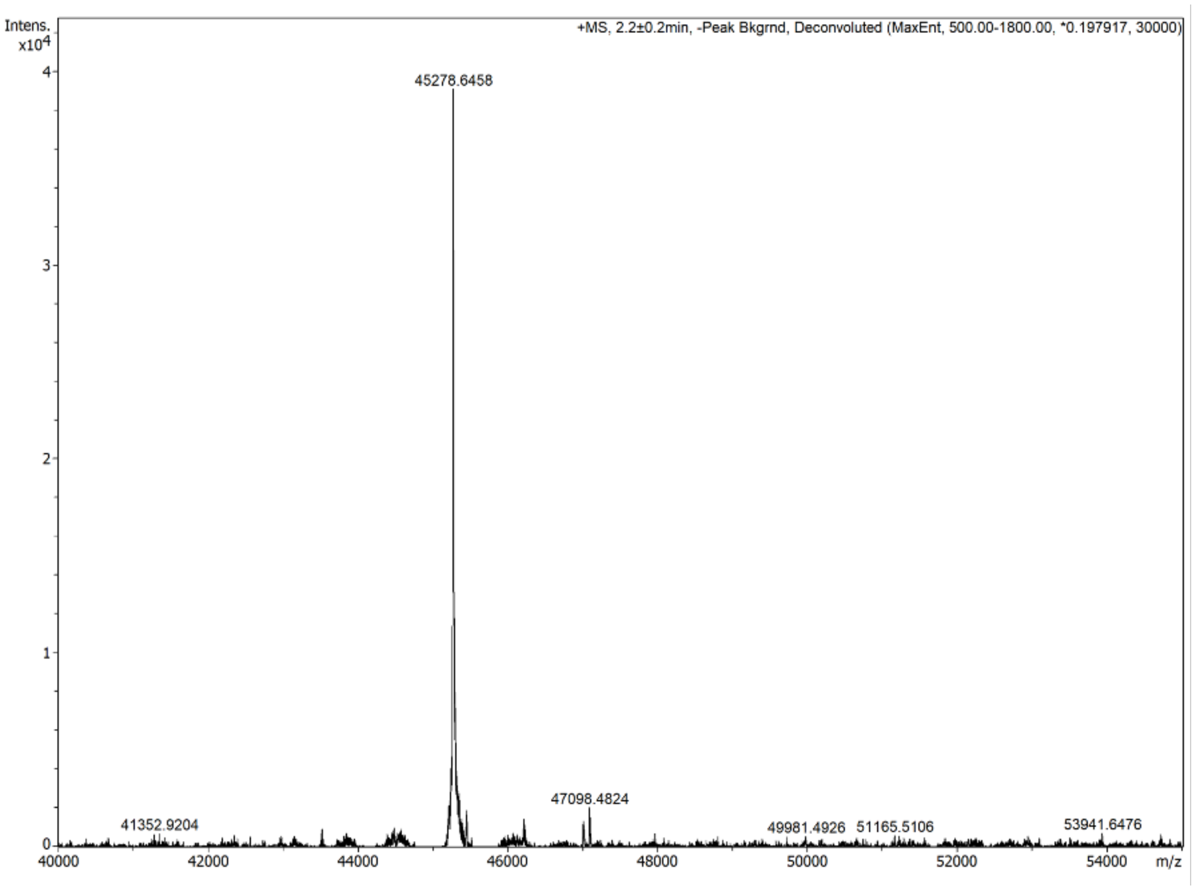


**Figure S49**. Deconvoluted intact mass spectrometry for $(His)_6$-BifE. The found *m/z* of 45,278.6458 Da corresponds to $(His)_6$-BifE minus its N-terminal methionine (Calc, *m/z* 45,45,278.34 Da).

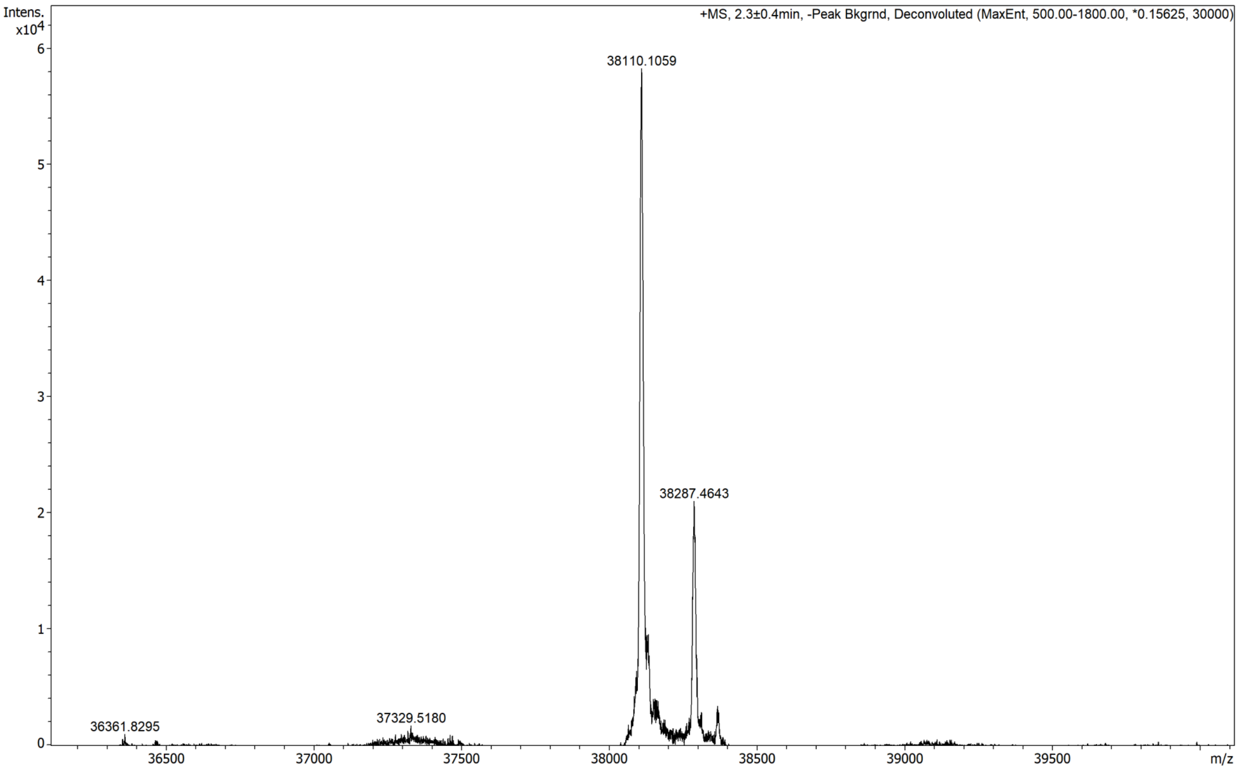


**Figure S50.** Deconvoluted intact mass spectrometry for $(His)_6$-BifF. The found *m/z* of 38,110.1059 Da corresponds to $(His)_6$-BifF minus its N-terminal methionine (Calc *m/z* 38,110.11). The second prominent compound with *m/z* 38,287.4643 Da corresponds to $(His)_6$-BifF minus its N-terminal methionine and harbouring an *N*-gluconoylated hexahistidine tag[53] (Calc *m/z* 38,288.17 Da).

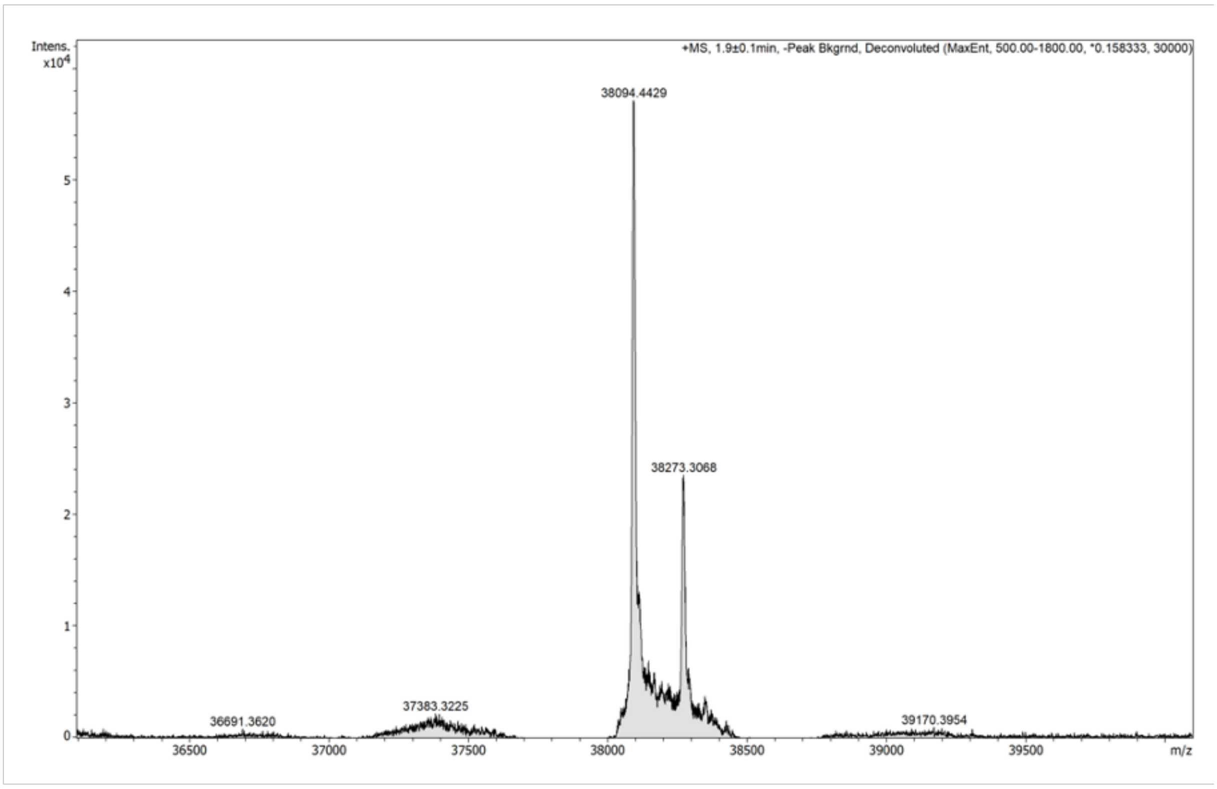


**Figure S51.** Deconvoluted intact mass spectrometry for $(His)_6$-BifN. The found *m/z* of 38094.4429 Da corresponds to $(His)_6$-BifN minus its N-terminal methionine (Calc *m/z* 38,094.69). The second prominent compound with *m/z* 38,271.5178 Da corresponds to $(His)_6$-BifN minus its N-terminal methionine and harbouring an *N*-gluconoylated hexahistidine tag[53] (Calc *m/z* 38,272.75 Da).

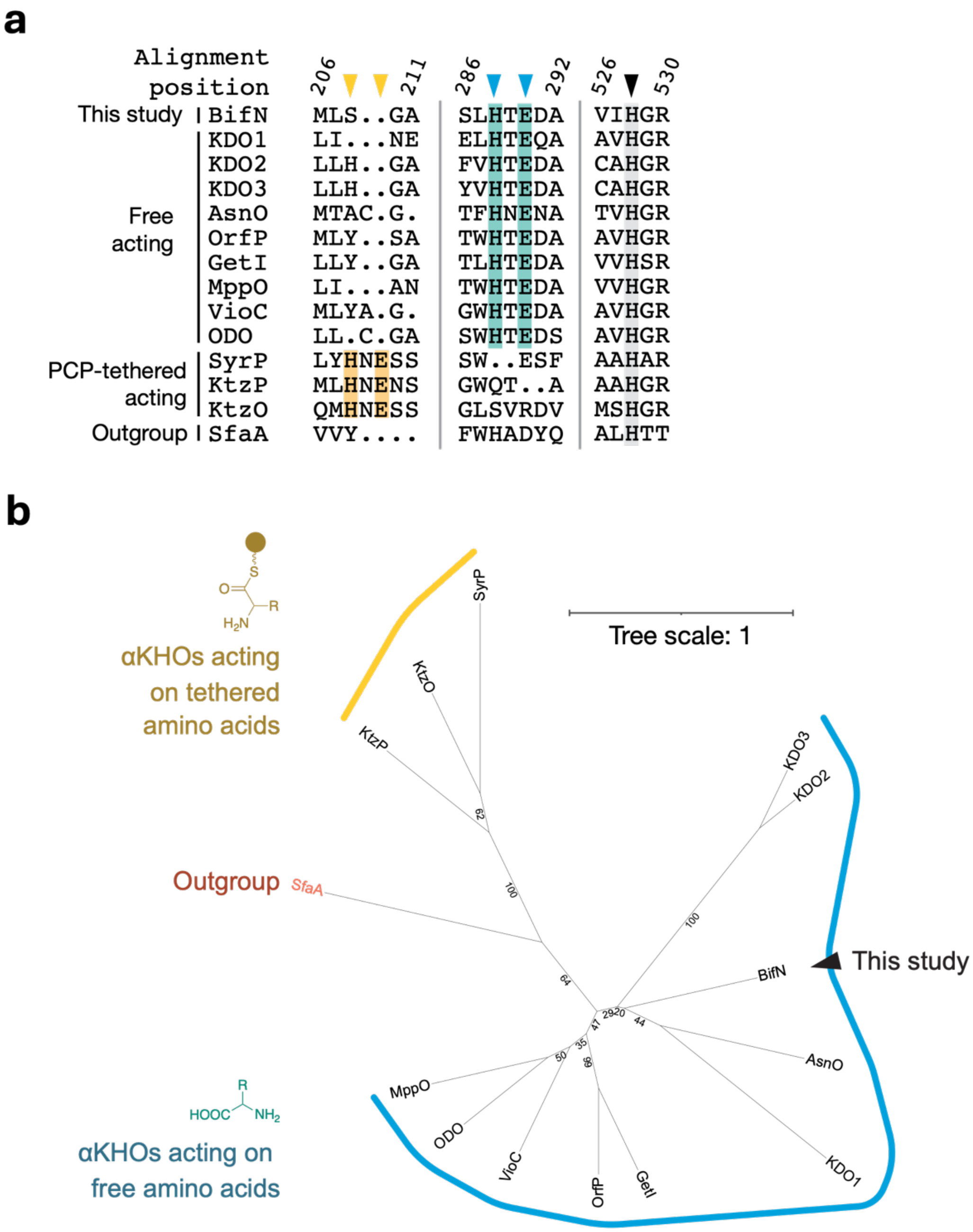


**Figure S52.** Bioinformatic analysis of experimentally characterised α-ketoglutarate-dependent hydroxylases (αKHOs). (a) Excerpt of a Muscle amino acid alignment of BifN with experimentally characterised αKHOs. Arrows indicate the His-Glu-His catalytic triad. Shading represents amino acid conservation. Protein sequences and the full alignment are available at 10.6084/m9.figshare.27958413. (b) An unrooted approximate maximum-likelihood tree of the αKHOs aligned in (a). Bootstrap support values are shown at nodes. Colours indicate αKHOs that act on free amino acid substrate(s) or those tethered to peptidylcarrier protein domains. SfaA, an α-ketoglutarate-dependent isonitrilase was included as an outgroup. The tree is available in Nerwrick format at 10.6084/m9.figshare.27958413.

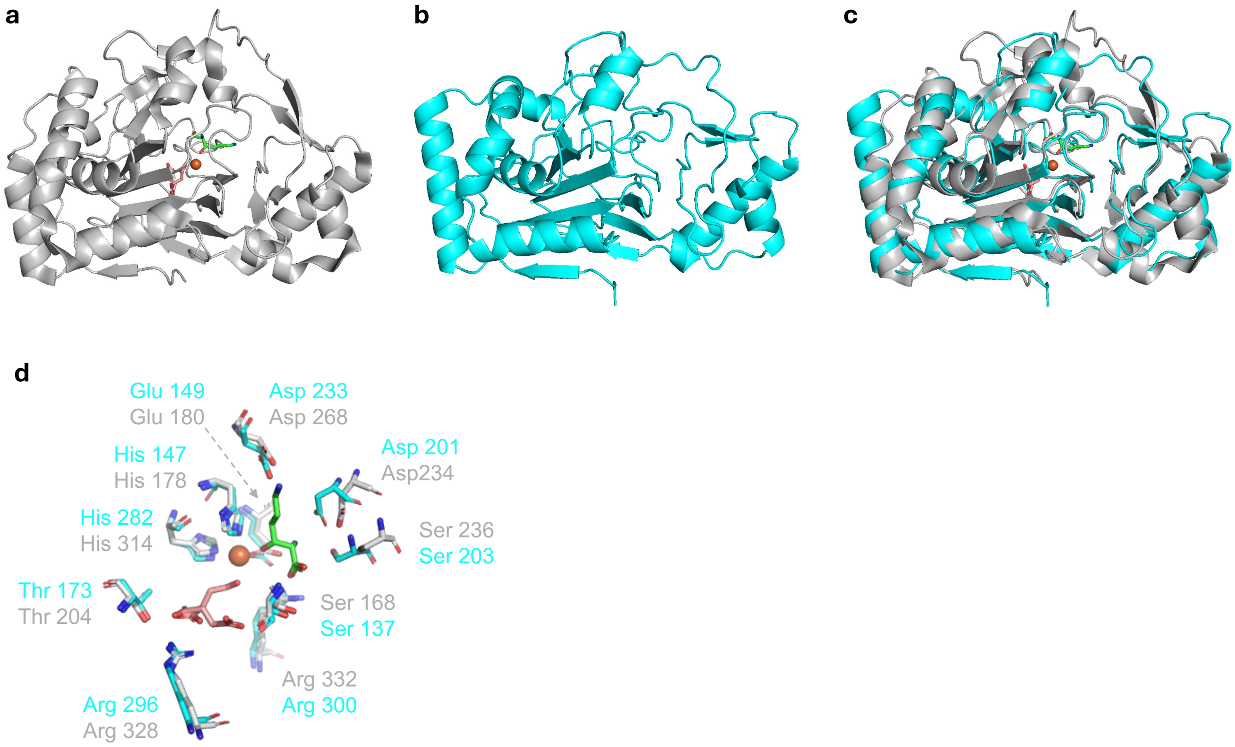


**Figure S53.** Structural analysis of BifN. (a) Crystal structure of KDO1 (PDB ID 6F6J) in complex with $Fe^{2+}$, succinate and (3*S*)-3-OH-ʟ-Lys product. (b) AlphaFold3 structural model of BifN. (c) Superimposition of KDO1 (grey) and BifN (cyan) using PyMol 2.5.4. (d) Comparison of the KDO1 (light grey) and BifN (cyan) active sites. For (a), (c) and (d) $Fe^{2+}$ (orange) and succinate (salmon) and (3*S*)-3-OH-ʟ-Lys (green).

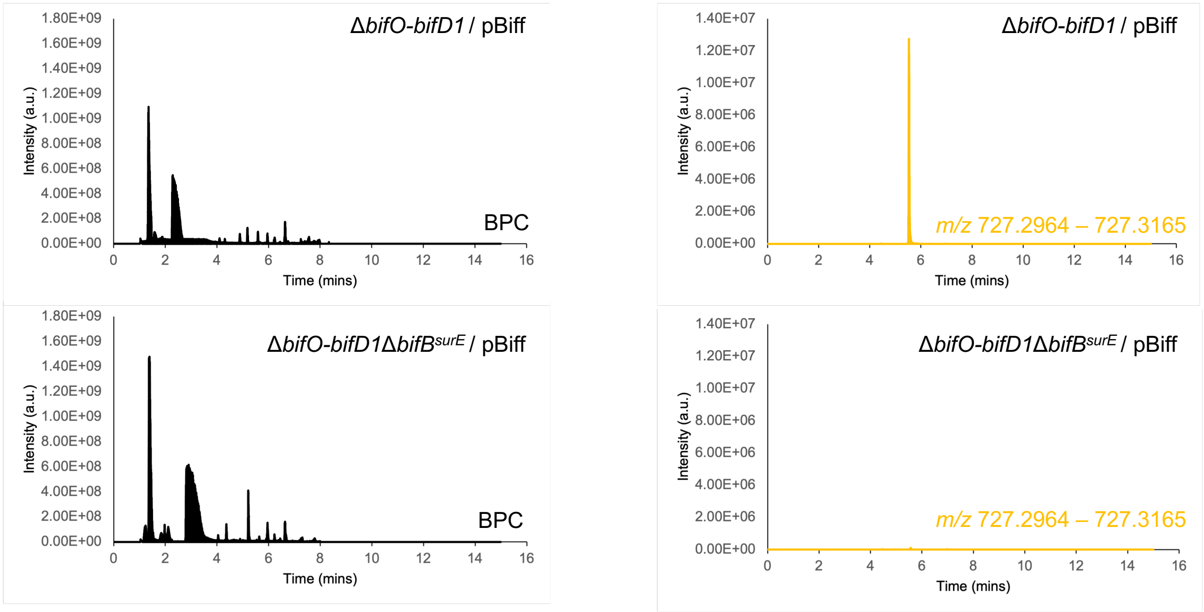


**Figure S54.** BifBSurE is required for biffamycin A production. Chemical extracts were prepared from the indicated strains and displayed is the base peak chromatogram (BPC, left, black) or exracted ion chromatogram (EIC, right, orange) for the monoisotopic mass corresponding to the $[M+H]^+$ adduct for 1 ($C_{32}H_{47}ClN_6O_{11}$) ± 0.01.

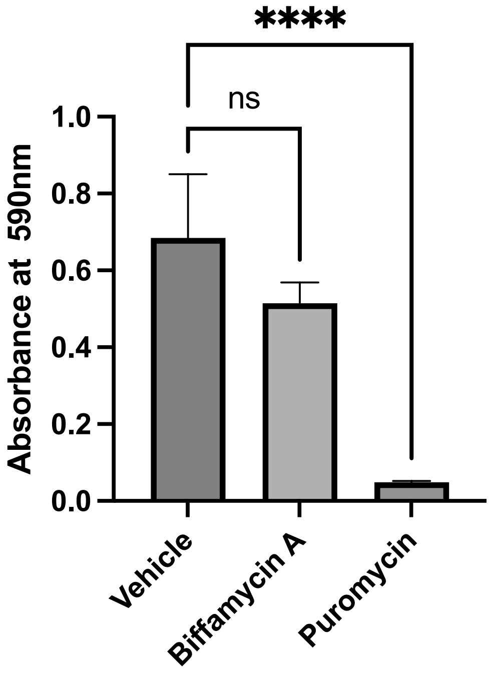


**Figure S55.** Biffamycin A is not cytotoxic to HEK293 cells. HEK293 cells were treated for 48 hr with either biffamycin A, puromycin or empty vehicle (sterile water) followed by crystal violet staining biomass., after which cellular protein content was quantified by incubation with MTT reagent and measuring absorbance at 527nm. Statistical significance was assessed by one-way ANOVA. Error bars denote the standard error of the mean; **** = $P$ < 0.00001, ns = no significance.

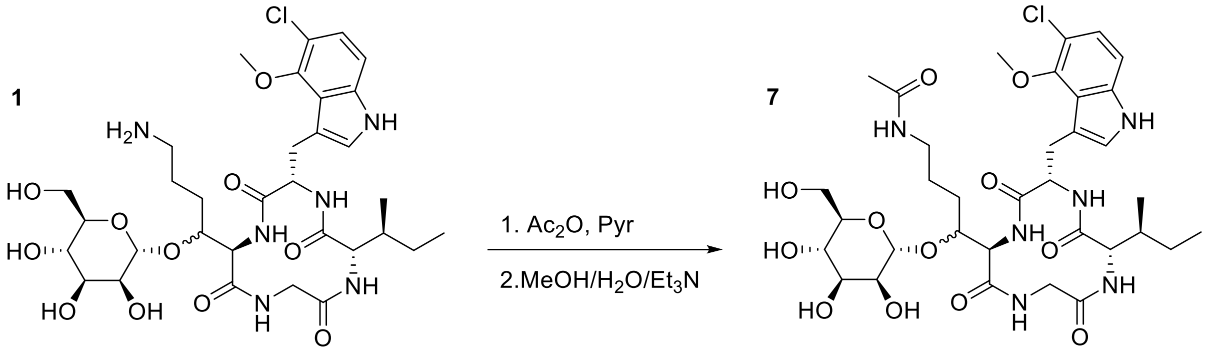


**Figure S56.** Synthetic scheme for *N*-acetyl biffamycin A.

**Table S8.** Bacterial strains and plasmids used in this study.

| Strain/plasmid | Description [a] | Reference |
|---|---|---|
| ***Streptomyces albidoflavus*** | | |
| S4 | Wild type *S. albidoflavus* S4 | [62] |
| Δ5 | Derivative of S4 harbouring mutations in biosynthetic pathways for: antimycin, candicidin, surugamide, fredericamycin and albaflavenone. | [12] |
| Δ5Δ*bifO-bifD1* | Derivative of Δ5 in which *bifD1D2EFGHIJKLMNO* were replaced with a hygromycin resistant cassette with an outwardly firing *ermE** promoter to drive transcription of *bifCBA;* $Hyg^R$ | This study |
| Δ5Δ*bifO-bifD1 attB* ΦC31::pBiff | Derivative of S4 Δ5Δ*bifO-bifD1* harbouring the pBiff plasmid containing constitutively expressed *bifD1D2EFHIJKNOP;* $Hyg^R$, $Apr^R$ | This study |
| ***Streptomyces coelicolor*** | | |
| M1152 | Derivative *S. coelicolor* M145 harbouring mutations in the biosynthetic pathways for: actinorhodin, undecylprodigiosin, calcium-dependent antibiotic and coelimycin and in *rpoB*[C1298T] and *rpsL*[A262G, C271T]. | [64] |
| M1152 *attB* ΦC31::pSET152ermEp | Derivative of M1152 harbouring pSET152ermEp at the ΦC31 *attB* site; $Apr^R$ | This study |
| M1152 *attB* ΦC31::pBifEFIJK | Derivative of M1152 harbouring pBifEFIJK at the ΦC31 *attB* site; $Apr^R$ | This study |
| M1152 *attB* ΦC31::pBifIJK | Derivative of M1152 harbouring pBifIJK at the ΦC31 *attB* site; $Apr^R$ | This study |
| M1152 *attB* ΦC31::pBifEF | Derivative of M1152 harbouring pBifEF at the ΦC31 *attB* site; $Apr^R$ | This study |
| M1152 *attB* ΦC31::pBifEIJK | Derivative of M1152 harbouring pBifEIJK at the ΦC31 *attB* site; $Apr^R$ | This study |
| M1152 *attB* ΦC31::pBifFIJK | Derivative of M1152 harbouring pBifFIJK at the ΦC31 *attB* site; $Apr^R$ | This study |
| ***Escherichia coli*** | | |
| BL21(DE3) | Host for heterologous protein production | Novagen |
| C41(DE3) | Host for heterologous protein production | Sigma-Aldrich |
| ET12567 | Non-methylating host for transfer of DNA into *Streptomyces* species; $Cam^R$ | [54] |
| GB0R-red | Host for RecET recombination | [53] |
| IMHA 659045 | Clinical isolate | Cubist |
| NEB5α | General cloning host | New England Biolabs |
| XL10(Gold) | General cloning host | Agilent Technologies |
| **Other organisms** | | |
| *Pseudomonas aeruginosa* PA14 | Clinical isolate | [55] |
| *Streptococcus pyogenes* ATCC19615 | Clinical isolate | ATCC |
| *Staphylococcus aureus* USA300 | Clinical isolate | [56] |
| *Staphylococcus aureus* VRSA-5 | Clinical isolate | [57] |
| *Mycobacterium smegmatis* ATCC101 | Type strain for *Mycobacterium smegmatis* | ATCC |
| **Cosmids and plasmids** | | |
| CosBif | Supercos1 derivative whose insert spans *bifO-bifC*; $Kan^R$, $Carb^R$ | [51] |
| pET28a | Commercial protein expression vector; $Kan^R$ | Novagen |
| pET28a-bifE | Derivative of pET28a harbouring *bifE* cloned into the NdeI-HindIII sites; $Kan^R$ | This study |
| pET28a-bifF | Derivative of pET28a harbouring *bifF* cloned into the NdeI-HindIII sites; $Kan^R$ | This study |
| pBifEFIJK | Derivative of pSET152ermEp harbouring constitutively expressed *bifEFIJK*; ΦC31 integrase, *attP* site, $Apr^R$ | This study |
| pBifIJK | Derivative of pSET152ermEp harbouring constitutively expressed *bifIJK*; ΦC31 integrase, *attP* site, $Apr^R$ | This study |
| pBifEF | Derivative of pSET152ermEp harbouring constitutively expressed *bifEF*; ΦC31 integrase, *attP* site, $Apr^R$ | This study |
| pBifEIJK | Derivative of pSET152ermEp harbouring constitutively expressed *bifEIJK*; ΦC31 integrase, *attP* site, $Apr^R$ | This study |
| pBifFIJK | Derivative of pSET152ermEp harbouring constitutively expressed *bifFIJK*; ΦC31 integrase, *attP* site, $Apr^R$ | This study |
| pBiff | Derivative of pSET152ermEp harbouring constitutively expressed *bifD1D2EFHIJKNOP;* ΦC31 integrase, *attP* site, $Apr^R$ | This study |
| pSET152*ermE*p | pSET152 derivative containing *ermE**p cloned into the EcoRV-EcoRI sites; $Apr^R$ | [55] |
| pUC19-pHygp | Derivative of pUC19 harbouring a hygromycin resistance gene flanked by FRT sites and divergently firing *rpsL(XC)* and ermE* promoters; $Carb^R$, $Hyg^R$ | [52] |
| pUZ8002 | Encodes conjugation machinery for mobilization of plasmids from *E. coli* to *Streptomyces;* $Kan^R$ | [54] |

[a] Carb, carbenicillin; Apr, apramycin; Hyg, hygromycin, Kan, kanamycin; Cam, chloramphenicol; *oriT*, origin of conjugal transfer

**Table S9.** Oligonucleotides used in this study.

| Name | Sequence (5´-3´)[a] | Description |
|---|---|---|
| DCLV179 | tgaccatatgCTGCCCAGCGCGCTG | PCR: Amplification of the BifF coding sequence |
| DCLV180 | gcataagcttTCATCGCTTGACGCCTGTGACCAG | PCR: Amplification of the BifF coding sequence |
| JEC221 | cgccgggcgtttttatctagaGATGCAGGCGACACCCAG | PCR: Amplfiication of homology directed repair arms for Cas9 deletion of BifB$^{SurE}$ coding sequence |
| JEC222 | ctggagcagggtgacctg**tca**CGTGTCCTCCTGCCGG | PCR: Amplfiication of homology directed repair arms for Cas9 deletion of BifB$^{SurE}$ coding sequence (bold text represents artificial stop codon to terminate BifB before SurE domain) |
| JEC223 | ccggcaggaggacacg**tga**CAGGTCACCCTGCTCCAG | PCR: Amplfiication of homology directed repair arms for Cas9 deletion of BifB$^{SurE}$ coding sequence (bold text represents artificial stop codon to terminate BifB before SurE domain) |
| JEC224 | tttacggttcctggcctctagaGATCGTCGTGAACGCGGTG | PCR: Amplfiication of homology directed repair arms for Cas9 deletion of BifB$^{SurE}$ coding sequence |
| JEC225 | acgcGGTAGTAGGTCAGCGCCGAG | CRISPR/Cas9 sgRNA for BifB$^{SurE}$ deletion |
| JEC226 | aaacCTCGGCGCTGACCTACTACC | CRISPR/Cas9 sgRNA for BifB$^{SurE}$ deletion |
| JEC245 | GACAAGGAACTGGTGCACGAC | PCR: Amplification of deleted BifB$^{SurE}$ locus |
| JEC246 | GAAGAGCCCGACGACACC | PCR: Amplification of deleted BifB$^{SurE}$ locus |
| MWB124 | gagtagacgacggagacgtaATGGAAACCGCACCGGCGTC | PCR: CPEC assembly of pBifEIJK |
| MWB125 | gacgccggtgcggtttccatTACGTCTCCGTCGTCTACTC | PCR: CPEC assembly of pBifEIJK |
| MWB126 | CGAAGCAGGGTTATGCAGCGGAAAAGATCCGTCGACCTGC | PCR: CPEC assembly of PCR products into pSET152ermEp |
| MWB127 | GCAGGTCGACGGATCTTTTCCGCTGCATAACCCTGCTTCG | PCR: CPEC assembly of PCR products into pSET152ermEp |
| MWB135 | gatcgcaggtgcacgcggtcgatcGCCCTGCAGGCGGAAGTCA GGTAG | PCR: CPEC assembly of pBifEF |
| MWB136 | ctacctgacttccgcctgcagggcGATCGACCGCGTGCACCTG CGATC | PCR: CPEC assembly of pBifEF |
| MWB137 | ggtcacaggcgtcaagcgatgagACTAGTATCGATGAATTCGT AATC | PCR: CPEC assembly of pBifFIJK |
| MWB138 | gattacgaattcatcgatactagtcTCATCGCTTGACGCCTGT GACC | PCR: CPEC assembly of pBifFIJK |
| RFS184 | CCATTATTATCATGACATTAA | CosBif insert end-sequencing |
| RFS185 | GTCCGTGGAATGAACAATGG | CosBif insert end-sequencing |
| RFS764 | gagtagacgacggagacgtaGTGACCCACGAGGGATCTCCGGG GGCAGGACCGGTACCGG | PCR: Amplification of P-Hyg-P recombineering cassette for activating *bifCBA* |
| RFS766 | cgccgagcaggaactggtgc | PCR: Amplification of *bifCBA* recombineered locus |
| RFS772 | gccggttggtaggatcgtctagaacaggaggccccatatgGCT AACAGCGTGGTGATTGT | PCR: Amplification of *bifKJI* for assembly into pSET152ermEp for construction of pBiff |
| RFS773 | gctatgacatgattacgaattcatcgatactagtggtaccTCA GCGCACGCTCCAGGAGG | PCR: Amplification of *bifKJI* for assembly into pSET152ermEp for construction of pBiff |
| RFS774 | gctcagggagtacgcctcctggagcgtgcgctgaggtaccGCC CTGCAGGCGGAAGTCAG | PCR: Amplification of *rpsL(XC)*p for assembly with *bifFE* for construction of pBiff |
| RFS775 | acacaccgtagacggccttctccagcgcgctgggcagcatTAC GTCTCCGTCGTCTACTC | PCR: Amplification of *rpsL(XC)*p for assembly with *bifFE* for construction of pBiff |
| RFS776 | cggccggtctccgtgagctcgagtagacgacggagacgtaATG CTGCCCAGCGCGCTGGA | PCR: Amplification of *bifFE* for assembly with *rpsL(XC)*p for construction of pBiff |
| RFS777 | gaaacagctatgacatgattacgaattcatcgatactagtTCA GCGGAACGCGAATCCCA | PCR: Amplification of *bifFE* for assembly with *rpsL(XC)*p for construction of pBiff |
| RFS778 | ctggtgtggacggggtggtgagggccgggggccggtccggatt ccggggatccgtcgacc | PCR: Amplification of P-Hyg-P recombineering cassette for activating *bifCBA* |
| RFS779 | ccgttctacttctggctgc | PCR: Amplification of *bifCBA* recombineered locus |
| RFS786 | GTATCCAAGGTGTACGGGTCG | PCR: Amplification of partial *bifD1D2* for assembly with gBlock for construction of pBiff |

| | | |
|---|---|---|
| RFS787 | aatttcacacaggaaacagctatgacatgattacGAATTCTCA CTCACGCCCCCCGATGG | PCR: Amplification of partial *bifD1D2* for assembly with gBlock for construction of pBiff |
| RFS788 | ctgctgggattcgcgttccgctgaactagtAGCCCGACCCGAG CACGCGC | PCR: Amplification of *ermE**p for assembly with *bifN* for construction of pBiff |
| RFS789 | gggttcgatgacaaaccttttcacacgaacCATATGGGGCCTC CTGTTCT | PCR: Amplification of *ermE**p for assembly with *bifN* for construction of pBiff |
| RFS790 | GTTCGTGTGAAAAGGTTTGT | PCR: Amplification of *bifN* for assembly with *ermE**p for construction of pBiff |
| RFS791 | TCAGACGACGACGCGCGACC | PCR: Amplification of *bifN* for assembly with *ermE**p for construction of pBiff |
| RFS792 | accgaggcccggtcgcgcgtcgtcgtctgaGCCCTGCAGGCGG AAGTCAG | PCR: Amplification of *rpsL(XC)*p for assembly with *bifH* for construction of pBiff |
| RFS793 | ctccgggtcgtcgaacgggttgacggtcatTACGTCTCCGTCG TCTACTC | PCR: Amplification of *rpsL(XC)*p for assembly with *bifH* for construction of pBiff |
| RFS794 | ATGACCGTCAACCCGTTCGA | PCR: Amplification of *bifH* for assembly with *rpsL(XC)*p for construction of pBiff |
| RFS795 | gaaacagctatgacatgattacgaattcatcgatactagtTCA GCGCCGTCCCATGGCCA | PCR: Amplification of *bifH* for assembly with *rpsL(XC)*p for construction of pBiff |
| RFS800 | cctggtgatggccatgggacggcgctgaactagtatcgatAGC CCGACCCGAGCACGCGC | PCR: Amplification of gBlock containing partial *bifD2* for assembly with remaining *bifD2D1* sequence for construction of pBiff |
| RFS801 | GCAGTTCGATACTCACCCCG | PCR: Amplification of gBlock containing partial *bifD2* for assembly with remaining *bifD2D1* sequence for construction of pBiff |
| RFS802 | ggctccggtggatgccatcgggggcgtgagtgagaattcGCC CTGCAGGCGGAAGTCAG | PCR: Amplification of *rpsL(XC)*p-*bifO* for assembly with *bifP* for construction of pBiff |
| RFS812 | tatatacatatgGAAACCGCACCGGCGTCCTT | PCR: amplification of the BifE coding sequence |
| RFS813 | tatataaagcttTCAGCGGAACGCGAATCCCA | PCR: amplification of the BifE coding sequence |
| RFS814 | tatatacatatgGTTCGTGTGAAAAGGTTTGT | PCR: amplification of the BifN coding sequence |
| RFS815 | tatataaagcttTCAGACGACGACGCGCGACC | PCR: amplification of the BifN coding sequence |
| RFS854 | ctcggcgccacgggccgggcggcgtccgcggcgtgctgtcTCA GCGGTAGAGGAACCGGT | PCR: Amplification of *rpsL(XC)*p-*bifO* for assembly with *bifP* for construction of pBiff |
| RFS855 | ccgcccggcccgtggcgccgagcgcagccctggaggacagATG GCGACGACGACCCTGGA | PCR: Amplification of *bifP* for construction of pBiff |
| RFS856 | aatttcacacaggaaacagctatgacatgattactacgtaTCA GGTCAGTGAGCAGCCCT | PCR: Amplification of *bifP* for construction of pBiff |
| gBlock | agcccgacccgagcacgcgccggcacgcctggtcgatgtcgga ccggagttcgaggtacgcggcttgcaggtccaggaaggggacg tccatgcgagtgtccgttcgagtggcggcttgcgcccgatgct agtcgcggttgatcggcgatcgcaggtgcacgcggtcgatctt gacggctggcgagaggtgcggggaggatctgaccgacgcggtc cacacgtggcaccgcgatgctgttgtgggcacaatcgtgccgg ttggtaggatcgtctagaacaggaggccccatatgaggagaac ttcgtcgcgtaaggcgccagttgactggtcgattcgc**ctc**gat tccgtatccaaggtgtacgggtcggaaaagagtccggttcagg ccctggacggggtgagtatcgaactgc | Synthetic DNA for changing the TTA codon for Leu17 in *bifD1* to CTC Leu17 (bold) |

[a]Red colour text indicates a restriction site; lowercase text indicates sequence does not anneal to the template; uppercase text indicates sequence anneals to template.
